# Herbarium records provide reliable phenology estimates in the understudied tropics

**DOI:** 10.1101/2022.08.19.504574

**Authors:** Daniel S. Park, Goia M. Lyra, Aaron M. Ellison, Rogério Katsuhito Barbosa Maruyama, Débora dos Reis Torquato, Renata C. Asprino, Benjamin I. Cook, Charles C. Davis

**Affiliations:** Department of Biological Sciences, Purdue University, West Lafayette, Indiana, 47906 USA; Purdue Center for Plant Biology, Purdue University, West Lafayette, Indiana, 47906 USA; Department of Organismic and Evolutionary Biology | Harvard University Herbaria, Harvard University, Cambridge, Massachusetts, 02138 USA; Programa de Pós-Graduação em Biodiversidade e Evolução, Instituto de Biologia, Universidade Federal da Bahia, Salvador, Bahia, 40170-115 Brasil; Harvard Forest, Harvard University, Petersham, Massachusetts, 01366 USA; Sound Solutions for Sustainable Science, Boston, Massachusetts, 02135 USA; Programa de Pós-Graduação em Botânica, Universidade Estadual de Feira de Santana, Novo Horizonte Feira de Santana, Bahia, 44036-900 Brasil; NASA Goddard Institute for Space Studies, New York, NY, 10025 USA; Ocean and Climate Physics, Lamont-Doherty Earth Observatory, Palisades, NY, 10964 USA

**Keywords:** Brazil, climate, citizen science, field survey, herbaria, natural history collections, neotropics, phenology

## Abstract

Plant phenology has been shifting dramatically in response to climate change, a shift that may have significant and widespread ecological consequences. Of particular concern are tropical biomes, which represent the most biodiverse and imperiled regions of the world. However, compared to temperate floras, we know little about phenological responses of tropical plants because long-term observational datasets from the tropics are sparse.

Herbarium specimens have greatly increased our phenological knowledge in temperate regions, but similar data have been underutilized in the tropics and their suitability for this purpose has not been broadly validated. Here, we compare phenological estimates derived from field observational data (i.e., plot surveys) and herbarium specimens at various spatial and taxonomic scales to determine whether specimens can provide accurate estimations of reproductive timing and its spatial variation.

Here we demonstrate that phenological estimates from field observations and herbarium specimens coincide well. Fewer than 5% of the species exhibited significant differences between flowering periods inferred from field observations versus specimens regardless of spatial aggregation. In contrast to studies based on field records, herbarium specimens sampled much larger geographic and climatic ranges, as has been documented previously for temperate plants, and effectively captured phenological responses across varied environments.

Herbarium specimens are verified to be a vital resource for closing the gap in our phenological knowledge of tropical systems. Tropical plant reproductive phenology inferred from herbarium records are widely congruent with field observations, suggesting that they can (and should) be used to investigate phenological variation and their associated environmental cues more broadly across tropical biomes.

## Introduction

Shifts in plant phenology–the timing of life-history events–are among the most iconic biological responses to climatic change and have widespread consequences for individual taxa and critical ecosystem processes (Willis et al. 2008; Polgar and Primack 2011; Richardson et al. 2013; Keenan et al. 2014;). Of particular concern are tropical biomes, which represent the most biodiverse and imperiled regions of the world (Kreft and Jetz 2007; Jenkins et al. 2013; Mora et al. 2013; Raven et al. 2020). However, there is comparatively little information on the phenological responses of tropical plants to climate change (Pau et al. 2011; Cook et al. 2012; Davis and Ellison 2018; Abernethy et al. 2018; Davis et al. 2022). Along these lines, most of our knowledge regarding the patterns, cues, and mechanisms of plant phenology comes from temperate biomes in North America, western Europe, and northeast Asia (Wolkovich et al. 2014; Park et al. 2021; Davis et al. 2022). Our understanding of tropical plant phenology has been especially limited by the overall paucity of long-term observational data and the overwhelming diversity of tropical species and their varied phenological behaviors (Abernethy et al. 2018; Davis et al. 2022). Tropical systems comprise diverse climates, from aseasonal rainforests to seasonally dry forests and grasslands. Moreover, various reproductive phenological strategies coexist in the tropics, including near complete synchrony and total asynchrony; sub- and supra-annual flowering; and short bursts of activity as well as continuous reproduction (Medway 1972; Frankie et al. 1974; Augspurger 1983; Bronstein and Patel 1992; Van Schaik et al. 1993; Newstrom et al. 1994; Galetto et al. 2000; Sakai 2002). Although these strategies may differ among and within species, tropical biomes as a whole do not exhibit a regular marked and reliable annual resting season in terms of plant reproductive activity as is common in temperate systems (Borchert 1996; Boulter et al. 2006; Zalamea et al. 2011; Morellato et al. 2013; Mendoza et al. 2017; Davis and Ellison 2018; Staggemeier et al. 2020; Davis et al. 2022). This has contributed to a ‘temperate phenological paradigm’, which we recently argued has been a major obstacle for understanding tropical phenology (Davis et al. 2022). To better understand such patterns in the tropics, we require more and better long-term records with greater spatiotemporal and taxonomic sampling.

Herbarium collections comprise large geographical, temporal, and taxonomic depth, and have been used to great effect in temperate zone investigations of phenology (Zohner and Renner 2014; Davis et al. 2015; Willis et al. 2017a; Willis et al. 2017b; Gallinat et al. 2018; Park et al. 2019; Park et al. 2021a). Herbarium specimens also have shown promise in studies of tropical plant phenology but have seen comparatively little use and have yet to be applied broadly (Borchert 1996; Boulter et al. 2006; Zalamea et al. 2011; Davis et al. 2018; Fava et al. 2019; Lima et al. 2021; Davis et al. 2022). On this front, Davis et al. (2022) recently demonstrated the likely utility of applying massive herbarium data for resolving tropical phenology, especially in Brazil. Several promising findings were identified in this effort, namely, that i.) phenological variation is great across the tropics, ii.) some biomes are much more sampled than others (e.g., Caatinga, Cerrado, Amazonia, and Atlantic Forest), and herbarium-based phenological observations are most abundant after only 1960, and that iii.) precipitation is a likely crucial factor for phenological cueing. In addition, the ongoing digitization and online mobilization of herbarium specimens have made them more widely available than ever before, and the onset of Digitization 2.0 *sensu* Hedrick et al. (2020) - the analysis solely of digitized collections - has enabled the efficient extraction of phenological information from specimens at a massive scale (Willis et al. 2017b; Hedrick et al. 2020; Davis et al. 2020; Park et al. 2019, 2021a).

Here, we harness a recently published dataset (doi to be provided upon acceptance) of high resolution phenological data scored from more than 32,000 freely available online specimens from Brazil and contrast these with direct field observation records to examine the phenological patterns of 24 phylogenetically diverse species across four major tropical biomes. To determine if herbarium specimens adequately represent flowering times in the tropics, we explicitly test whether phenological data inferred from herbarium specimens differ from data collected from field surveys. In summary, we demonstrate that herbarium specimens provide reliable information in the tropics across a broad range of taxa.

## Materials and Methods

We searched the literature and collected field observational phenological data for 24 species spanning diverse angiosperm clades (Bignoniaceae, Chrysobalanaceae, Fabaceae and Malpighiaceae; Table S1). These species were chosen for their broad representation in the neotropics, and availability of digitized images of herbarium specimens. After discarding studies with missing data (i.e., gaps in phenological observations), we were left with observational data from nine studies (23 species) across Brazil. Phenological observations were available as presence/absence information of flowering/fruiting events per month/species. These field observations spanned four diverse tropical biomes, Amazonia, Atlantic Forest, Caatinga, and Cerrado. Amazonia comprises the largest tropical rainforest in the world, with factors such as altitude, vegetation cover and inundation patterns driving the formation of diverse communities therein (Ferreira et al. 2015). The Atlantic Forest comprises dense, mixed and seasonally deciduous and semi-deciduous forests as well as mangroves and restinga vegetation (Duarte et al. 2014). The seasonally dry tropical forests in Caatinga and the neotropical savanna of Cerrado can experience up to 10 months of drought a year (Terra et al. 2018) and are counted among the most endangered biomes on the planet (Hoekstra et al. 2004). However, Cerrado benefits from the presence of the headwaters of the three largest hydrographic basins in South America (Amazonian, São Francisco and Prata). Cerrado vegetation is also characterized by the marked presence of adaptations to fire (Terra et al. 2018).

We simultaneously gathered digitized specimen images and associated metadata for 4638 specimens across these 23 species from a variety of online aggregators, including REFLORA (Forzza et al. 2016), SpeciesLink, iDigBio, and Tropicos. Each species was represented by at least 100 unique herbarium specimens. Citizen-scientists hired through Amazon’s Mechanical Turk service (MTurk; https://www.mturk.com/) counted the number of buds, flowers, and fruits to assess peak flowering time using the CrowdCurio interface following Willis et al. (2017b).

Crowdworkers were required to discern and quantify the different organs present on a test specimen with at least 80% accuracy across three trials before they could participate in the actual tasks. To provide an estimate of reliability, each image set scored by a single crowdworker included a single duplicate image randomly selected from the others (Williams et al. 2017). Species with reproductive organs that were difficult to discern were further examined by experts (i.e., the authors of this study). We estimated the consistency score for each participant based on the data for each image set by dividing the absolute difference in counts for each organ by the total count of that specimen across the two duplicate specimens and subtracting this value from 1 (1 – (|count1 – count2| / (count1 + count2)) (Williams et al. 2017; Park et al. 2019). Consistency scores range from zero (unreliable/inconsistent) to one (reliable/consistent). Participants who reported no organs on one sheet and a non-zero number of the same organ on the duplicate sheet were assigned a reliability score of zero for that organ (i.e., the lowest reliability score). Each specimen was examined by at least three people and we weighted their counts of each organ by their consistency scores and averaged them to obtain a single set of quantifications per specimen. These quantifications were used to separately infer the start, end, and length of flowering and fruiting periods at the municipality, state, biome, and country level using the data collected therein. Flowering and fruiting periods were defined as the period of time between the earliest and latest observations of flowering and fruiting individuals, respectively. Although tropical systems comprise both seasonal and aseasonal climates, plant reproductive activity occurs year-round (Morellato et al. 2013; Mendoza et al. 2017). Given the non-resting nature of tropical systems, we determined the start and end of flowering and fruiting periods from a circular distribution of collection dates (Morellato et al. 2010; Staggemeier et al. 2020; Davis et al. 2022) of specimens with at least a single flower or fruit present (≥ 1) using the *circular* package (Agostinelli and Lund 2022) in R v3.6.3 (R Core Team 2017). Likewise, we determined the start, end, and length of flowering and fruiting periods from a circular distribution of field observation dates.

To determine whether phenological inference from specimen data differed from field surveys, we compared the phenological period of each species as observed in the field with the circular 95% highest posterior density interval of flowering/fruiting periods inferred from herbarium collection dates within the same spatial category (i.e., municipality, state, biome, and country). Where there was no overlap between the 95% highest posterior density interval of the specimen collection dates of a species and the field observed phenological period in the same spatial category, we concluded that the two were significantly different at p < 0.05. The circular 95% highest posterior density interval of specimen collection dates was calculated using the *hpd_est_circ* function in the R package *bpnreg* (Cremers 2018). We also applied linear mixed models to examine the effects of these two methods of phenological inference on the inferred length of flowering/fruiting periods at each spatial scale. Data source (herbarium specimen or field observation) was also included as a fixed effect, and species identity entered the models as random effects. Analyses were conducted using the *lme4* package (Bates et al. 2015) in R v3.6.3 (R Core Team 2017). In total, comparisons of field survey and specimen derived phenological information were made across 23 species spanning 4 biomes, 6 states, and 5 municipalities in Brazil (Fig. S1). On average, municipalities were 2,839 ± 4,791 km^2^, states 532,276 ± 612,210 km^2^, biomes 2,042,371 ± 1,526,713 km^2^ in size, and the area of Brazil is approximately 8,515,767 km^2^.

Climate data, including maximum air temperature and total precipitation, were downloaded from CHELSA (https://chelsa-climate.org/) at 30 arc second resolution and resampled at the municipality level for each year (Karger et al. 2017). These data were used to compare the sampling of climate space between the two data categories–field versus herbarium observations of phenology–and to demonstrate how herbarium specimens may be used to investigate the environmental drivers of tropical phenology.

## Results

Herbarium specimens represented a much broader sampling across geographic and climatic space than field observational data (Fig 1). No observational studies were outside the geographic and climatic range represented by herbarium collections of the same species. Most flowering herbarium specimens were collected in the same months (Fig. 2a) and temporal period (Fig. 2b) in which flowering was observed in monthly field surveys, especially when specimens were collected in the same municipality as the observations. The degree of overlap between flowering periods estimated from field surveys and specimens varied across species and spatial scale (Appendix S1). However, less than 5% of the species in our dataset exhibited significant differences between flowering periods inferred from observations or specimens, regardless of spatial aggregation (p < 0.05). Flowering periods inferred from herbarium specimens did not differ from those inferred from field observations in the same municipality. The proportion of species with significant differences between flowering periods inferred from field versus herbarium observations within the same state, biome, and country were 4.8%, 4.2%, and 4.5% respectively.

**Figure 1.**
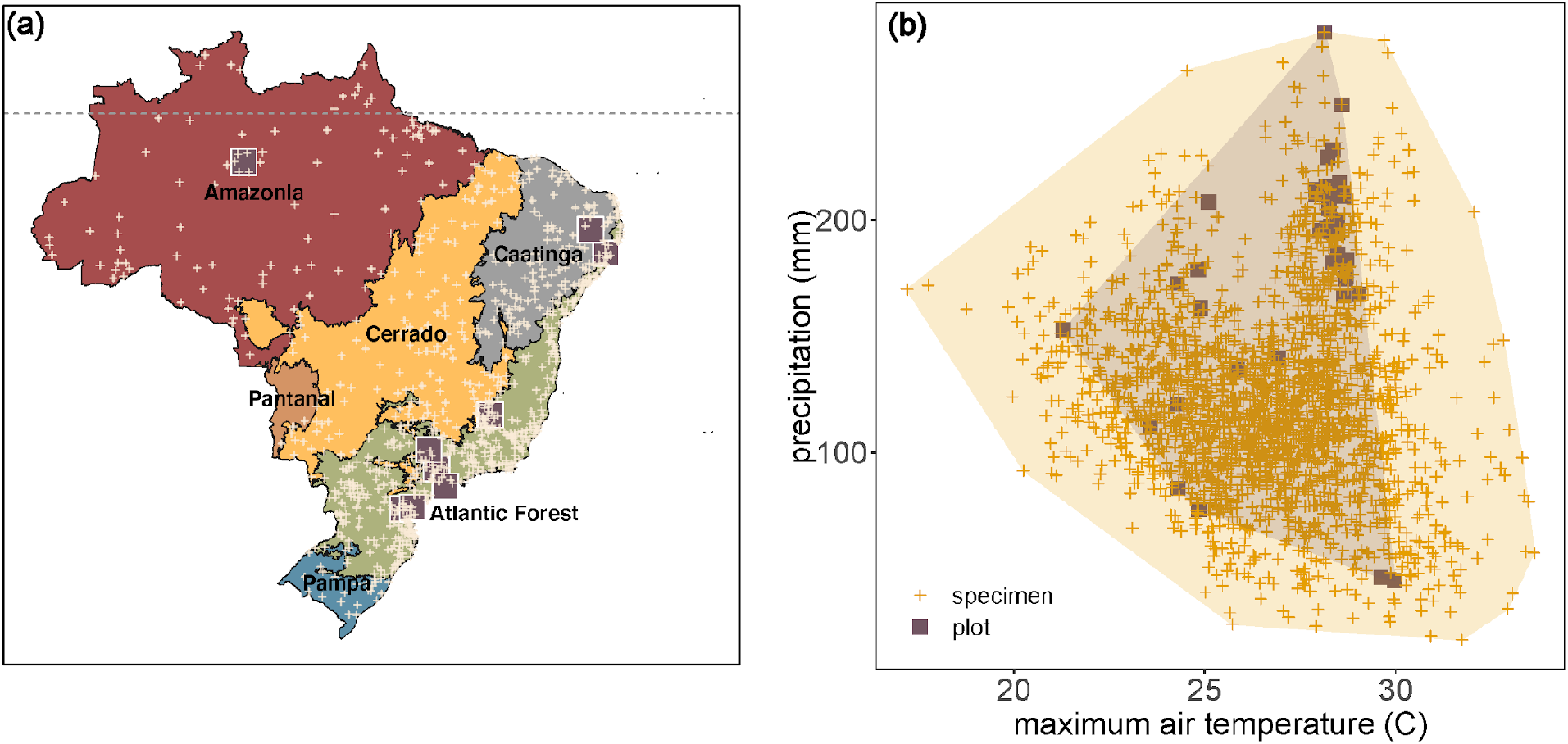
Distribution of field versus herbarium observations of phenology. Field observations of plant phenology (purple squares) and herbarium specimen collection locations (yellow crosses) in geographic (a) and climatic space (b). Each biome is depicted in a different color on panel (a). Shaded polygons in panel (b) are convex hulls encompassing all data points from each source, and the x and y axes refer to average monthly maximum air temperature and precipitation of the year/location of collection/observation respectively.

**Figure 2.**
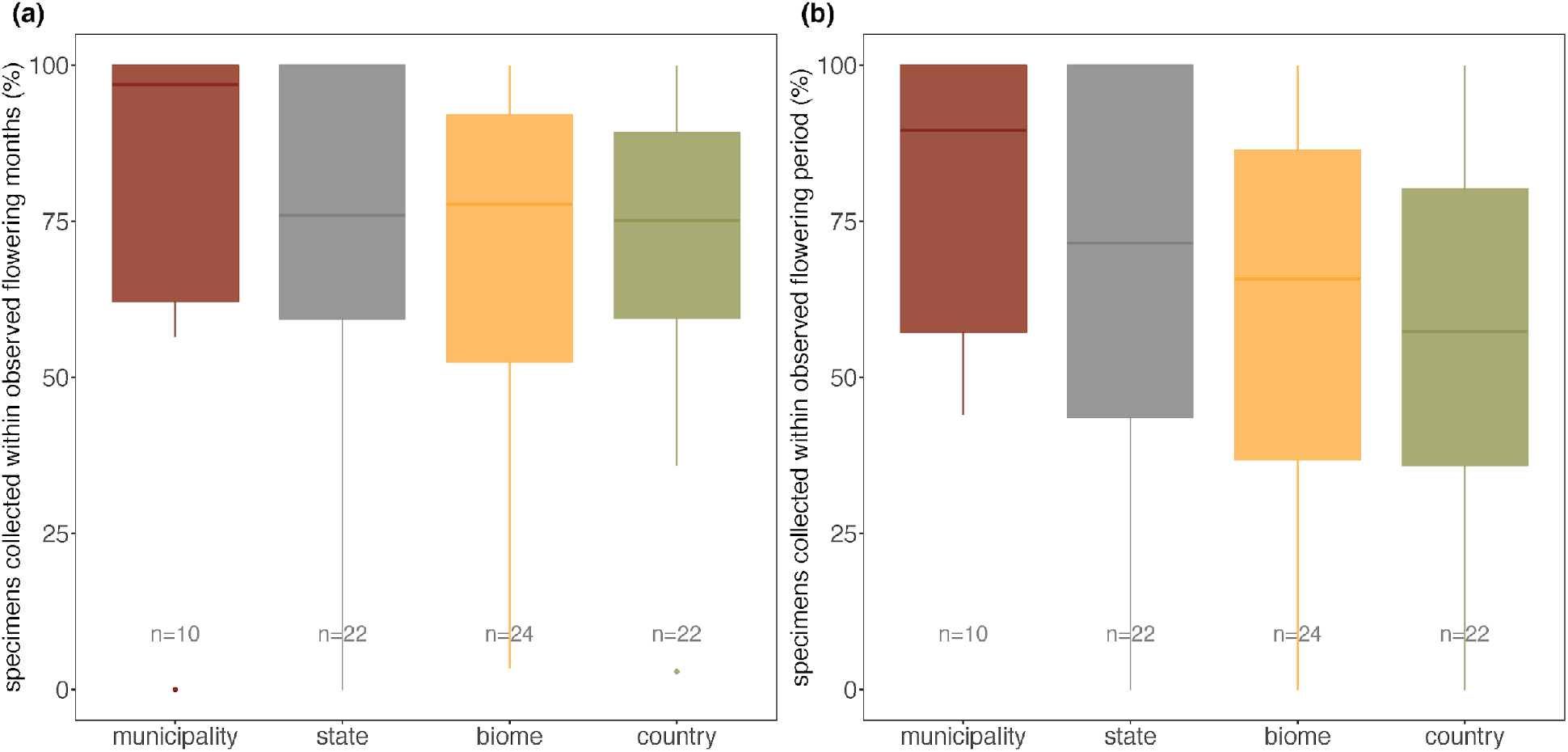
Proportion of herbarium specimens with flowers collected within the same months (a) and periods (b) of flowering observations in field surveys from the same municipality, state, biome, and country. The number of data points at each spatial scale are listed beneath each box and whisker plot. These species were compared across five municipalities, six states, four biomes, and the entire country of Brazil.

The duration of flowering period inferred from specimens was positively correlated with field observations, especially at smaller spatial scales (Fig. 3). Fruiting durations inferred from herbarium specimens were positively correlated with field observed durations as well, but not significantly so, regardless of spatial scale (Fig. S2). There was no significant difference in the length of flowering and fruiting periods inferred from field observations and herbarium specimens at the municipality and state levels (Table S2). However, flowering and fruiting periods inferred from herbarium specimens were significantly longer at the biome and country scales. Along these lines, species could exhibit phenological variation across their ranges not captured by the more narrow geographic focus of the field observations. For instance, our crowdsourcing results suggested *Chamaecrista desvauxii* (Collad.) Killip flowering phenology may differ substantially among Amazonia, Caatinga, Cerrado, and Mata Atlântica (Fig. 4). Overall similar patterns were observed for fruiting phenology, but the discrepancies between estimates derived from herbarium specimens and field observations tended to be larger (Appendix S1).

**Figure 3.**
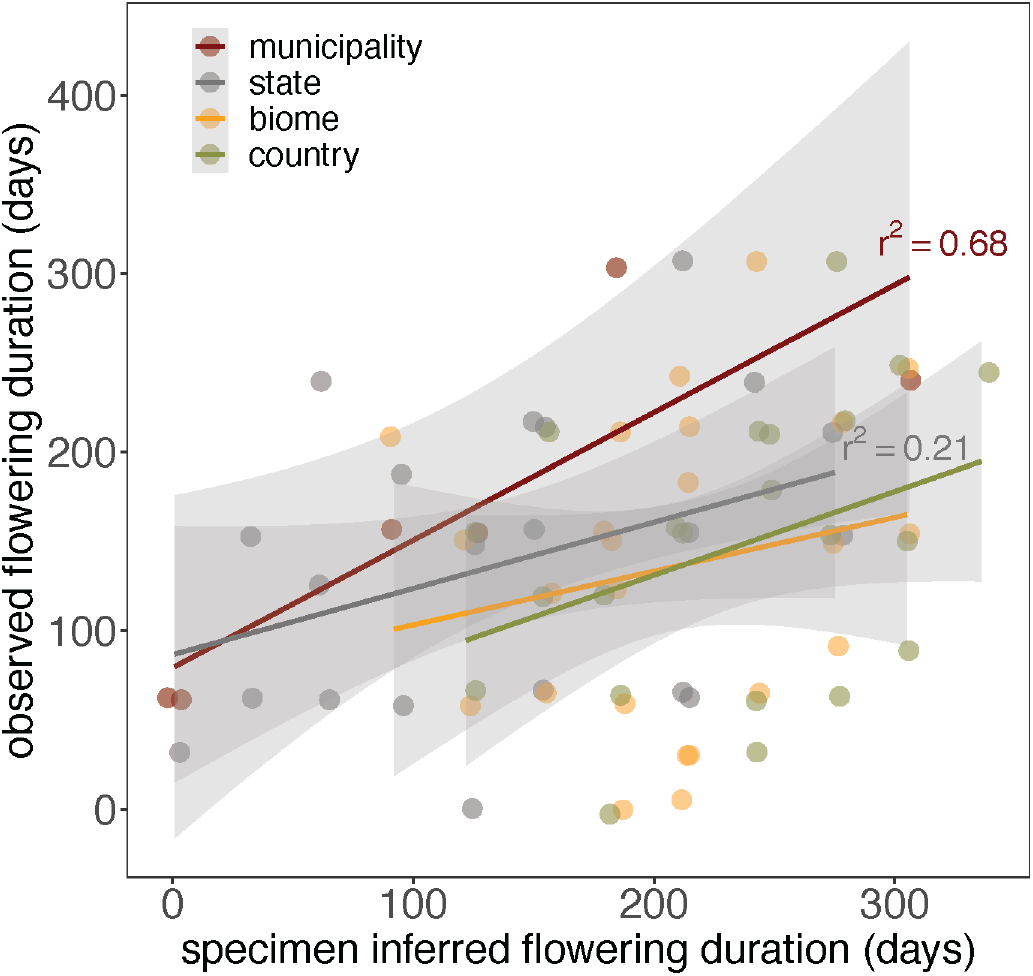
Comparison of flowering period inferred from field versus herbarium observations at varying spatial scales. R^2^ values are depicted only for significant correlations (p < 0.05).

**Figure 4.**
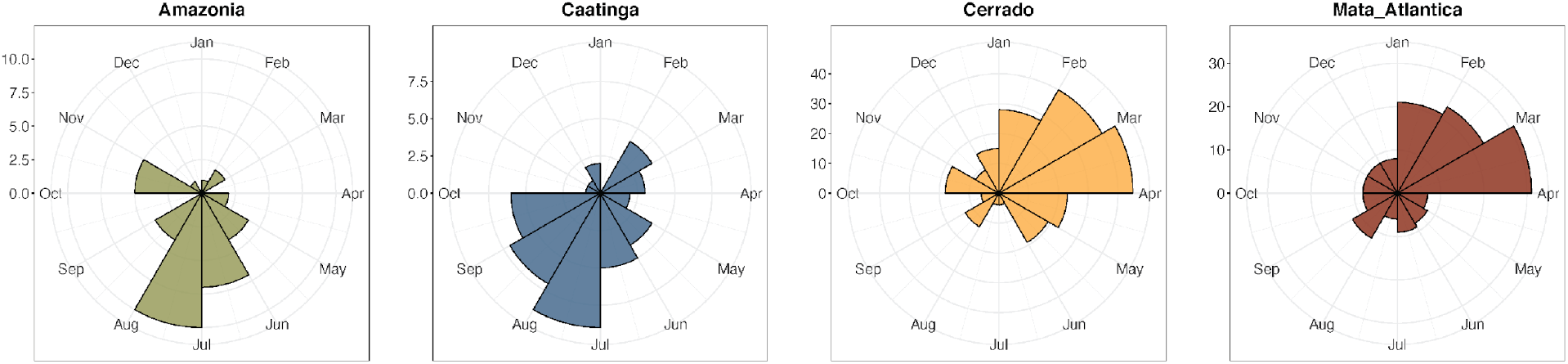
Histogram of *Chamaecrista desvauxii* flowering specimens collected in four biomes. Circular diagrams depict the time of collection in angles.

## Discussion

Tropical biomes are both simultaneously the most biodiverse and most threatened by anthropogenic change (Raven et al. 2020). To understand the effects and consequences of global change on processes that shape tropical ecosystems, such as plant reproductive phenology, we require data that span vast spatial, temporal, and taxonomic scales. Yet direct field observations of tropical phenology are rare (Abernethy et al. 2018; Davis et al. 2022). The utility of herbarium specimens to bridge this impasse to investigate tropical phenology has not been assessed broadly. Although such assessments have been attempted for temperate biomes, temporal and environmental cues for phenological states are likely to differ between temperate and tropical regions (Borchert et al. 2005; Davis et al. 2022). Our results indicate that herbarium specimens are indeed informative for phenological research in the tropics, yet need to be explored cautiously.

### Herbarium specimens provide reliable estimates of tropical phenology

Leaf phenology is by far the most common subject of studies examining the effects of climate change on plant phenology, especially at large scales (Park et al. 2021b). However, flowering and fruiting phenologies are key to understanding the range of individual responses, the fitness and survival of plant species, and the potential of cascading trophic changes within ecosystems (Ting et al. 2008; Willis et al. 2008; Polansky and Boesch 2013; Butt et al. 2015; Morellato et al. 2016; Mendoza et al. 2017). Our results demonstrate high congruence between reproductive phenology inferred from field observations and digitized herbarium specimens at every scale we analyzed. Despite the potential spatial biases of herbarium collections (Daru et al. 2018), both the geographic and climatic ranges represented by herbarium collections were substantially larger, and fully encompassed the range of observational data. As they do in temperate regions, herbarium specimens represent phenological observations across a much wider spatial extent and variety of tropical biomes and environments (Davis et al. 2015). Further, the herbarium specimens in our dataset comprised phenological information spanning more than 60 years, demonstrating that these specimens can help address the paucity of historic and long-term observational datasets in the tropics (see also Davis et al. 2022). Indeed, most of the field observations we reviewed were relatively recent (1988 – 2008), whereas many specimens were collected before the onset of what we now recognize as substantial global warming. We note that although specimens span greater temporal scales than most field observations, they do not necessarily comprise regular, repeated sampling of the same locale through time. Thus, while we may infer coarse temporal trends from specimens and complement the results from field observations, they do not replace the need for regularly censused, long-term observational investigations.

As with flowering, estimates of fruiting phenology were generally congruent between specimens and field observations, but to a lesser degree. We suggest four reasons for the weaker congruence of fruiting estimates. First, most herbarium specimens tend to be of plants in flower, resulting in fewer fruiting specimens from which to infer phenology. This can result in greater uncertainty and stochasticity in specimen-derived estimates of fruiting phenology. Second, fruits in some tropical species may persist on the parental plant well beyond maturation, across months or even years. Thus, it can be difficult to determine how old a fruit is from a digitized specimen, which can greatly complicate the interpretation of these scores. Third, smaller fruits may be confused with leaf/or flower buds during scoring (Fig. 5), reducing the accuracy of fruiting phenology estimates (see also Willis et al. 2017b and Davis et al. 2020) for variation across the scores of different phenophases).

**Figure 5.**
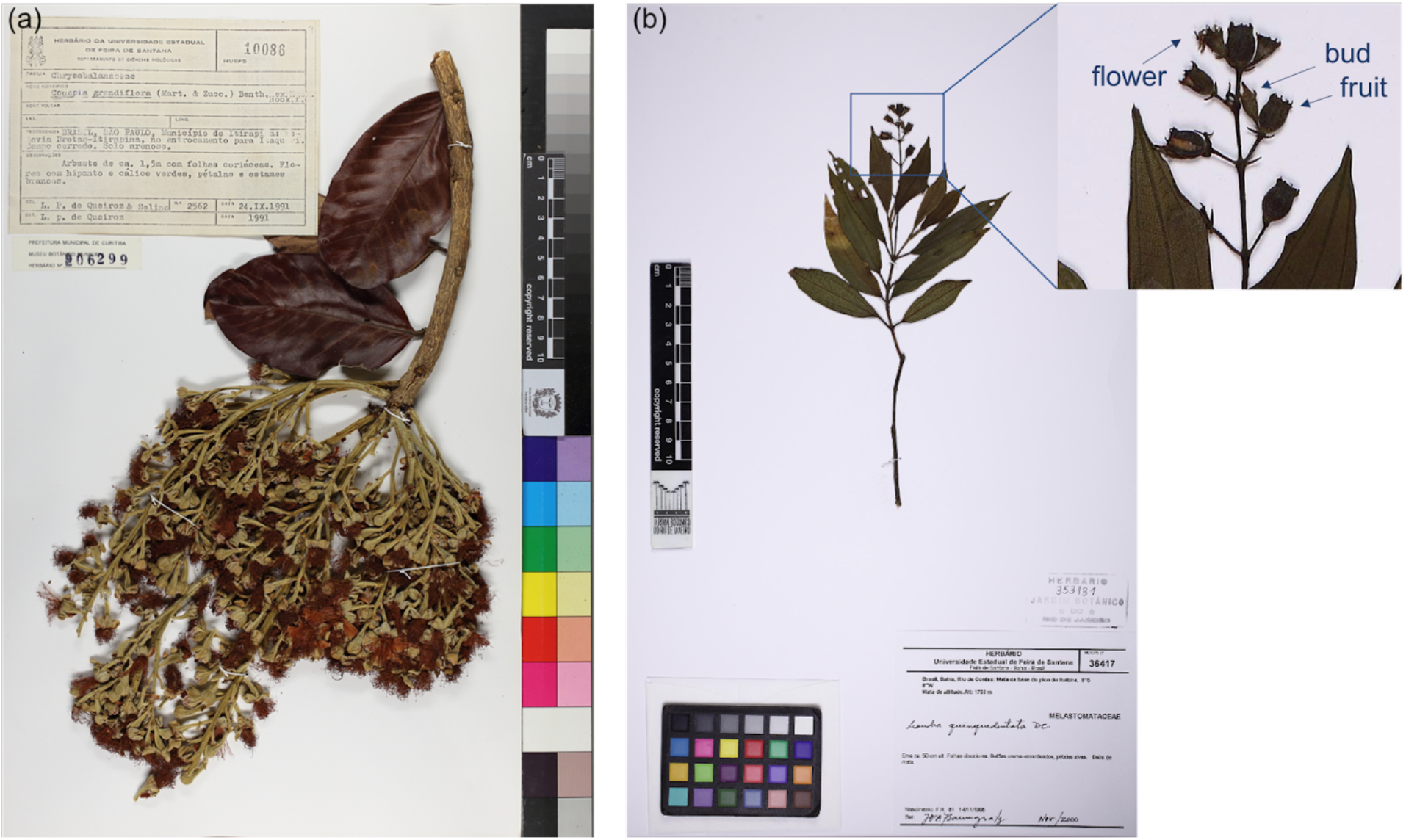
Specimens of *Couepia grandiflora* (a) and *Leandra quinquedentata* (b), illustrating overlap and similarity among reproductive organs. Images are from the Reflora Virtual Herbarium under CC BY-SA 4.0.

### The promise of herbarium specimens for determining phenological cues in the tropics

Numerous studies have documented phenological patterns in the tropical regions of the world, and factors such as precipitation, insolation, and photoperiod have been suggested to influence these events (Borchert 1996; Borchert et al. 2005; Borchert et al. 2015; Calle et al. 2010). However, most of these did not attempt to test the environmental or physiological drivers of these patterns (Abernethy et al. 2018). Herbarium specimens have been used to investigate the drivers of plant phenology in temperate regions, and similar approaches could potentially be applied to tropical systems (Davis and Ellison 2018; Davis et al. 2022). For example, herbarium specimens of the monkey’s comb, *Amphilophium crucigerum* (L.) L.G.Lohmann (Bignoniaceae) imply the same climate – flowering phenology relationships as field observations (Fig. 6). *A. crucigerum* tends to flower earlier in wetter (and slightly cooler) climates in both field surveys and herbarium specimens. Although the general lack of observational data prevents us from making more concrete inferences, this demonstrates the promise of using herbarium specimens to investigate environmental drivers of tropical phenology.

**Figure 6.**
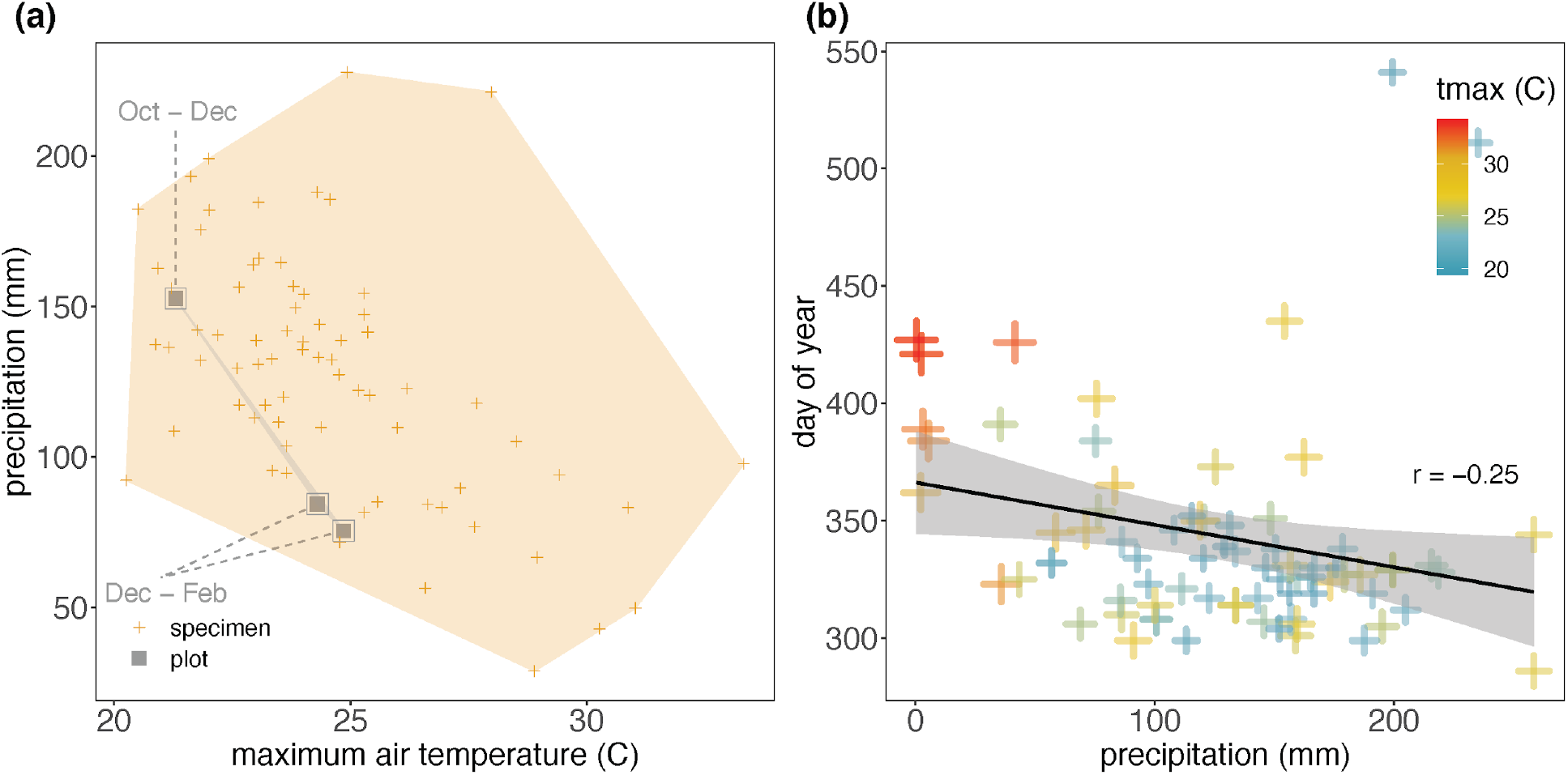
*Amphilophium crucigerum* phenology and its relationship with climate. (a) Flowering phenology field observations (grey squares) versus herbarium specimens (yellow crosses) are depicted in climate space. Maximum air temperature and precipitation represent monthly averages of the year/location of collection/observation respectively. Crosses represent specimens and grey squares represent surveyed plots. Grey text insets indicate the flowering periods of this species as surveyed in each plot. (b) The relationship between day of year flowering and mean September-October precipitation inferred from herbarium specimens. Both field observation data and specimen data indicated that *A. crucigerum* flowering begins in October and persists into the following year.

The phenological timing of species and their responses to climate have been demonstrated to vary substantially across latitude, both within and among species (Park et al. 2019; Park et al. 2021b). Although these studies focused on temperate floras, we expect similar responses may exist in tropical biomes as well, as the ranges of numerous taxa span multiple tropical biomes (Lucresia et al. 2021). For instance, *Banisteriopsis pubipetala* (A. Juss.) Cuatrec. (Malpighiaceae) has been independently observed to flower from July to December in Cerrado (Batalha and Mantovani 2000), while flowering may be restricted to October in Mata Atlântica (Morellato and Leitao-Filho 1996). However, it is difficult to assess phenological variation across their ranges from existing, geographically restricted field observations. As in temperate regions, herbarium specimens collected from across species’ ranges can be used effectively to explore how their phenological responses vary across space and time. Along these lines, the decrease in congruence we observe between phenological timing inferred from field observations and herbarium specimens at larger spatial scales (e.g., country) may reflect the presence of widespread phenological variation across species’ ranges in the tropics.

### Limitations, caveats, and ways forward

Herbarium specimens represent a non-random, non-comprehensive sampling of phenological events (Davis et al. 2015; Willis et al. 2017a; Daru et al. 2018) because they are rarely collected for phenological studies and reflect gaps and biases related to collector behavior. Taxonomic challenges, including misidentified specimens and species complexes, can mislead results of all phenological studies, but especially those based on digital specimen images and particularly for tropical plants, which are notoriously misidentified (Goodwin et al. 2015). For instance, we found disparities in phenological information inferred from specimens and field observations in *Banisteriopsis variabilis* B.Gates (Malpighiaceae) and *Licania heteromorpha* Benth (Chrysobalanaceae) (Appendix S1; see also Davis et al. 2022). As their specific epithets suggest, both taxa are morphologically variable and may constitute species complexes (Prance 1972). However, taxonomic difficulty can affect field-based studies as well, and collections-based studies have increasingly found ways to detect and account for such biases (Park et al. 2019; Belitz et al. 2020). Moreover, these instances potentially represent opportunities to explore cryptic species that may be delimited by phenology.

Inferring phenophases from digital images can be difficult for certain taxa. For instance, most species of *Miconia* and *Leandra* (Melastomataceae), and most species of Chrysobalanaceae possess tiny buds and flowers, often clumped and/or overlapping in specimens (Fig. 5). Fruits may be harder to recognize in several species, as they can be confused with flower or leaf buds. Quantifying reproductive organs and assessing the phenological stage of such taxa can be difficult when the physical specimens are not available for direct examination. Thus, digital specimen-based investigations of phenology may not be appropriate for all taxa. Still, though time consuming, expert botanists are able to discern and quantify different reproductive organs of these taxa from high-resolution digital images as we demonstrate here. Further, advances in machine-learning applications for phenological research are making the automatic extraction of data from digitized specimens with equal or better accuracy than non-expert humans increasingly feasible (Lorieul et al. 2019; Davis et al. 2020; Goëau et al. 2020).

Inferring phenological phenomena from herbarium specimens faces additional challenges in tropical biomes. Some species can reproduce multiple times during a single year, while others may reproduce supra-annually (Bawa et al. 2003; Engel and Martins 2005; Rojas-Robles and Stiles 2009). For example, *Licania octandra* var. *pallida* Prance (Chrysobalanaceae) flowered four times in 25 years of field observations in the Reserva Ducke (Amazon Rainforest) from 1970 to 1994 (Ruiz and Alencar 1999). Though the density of flowering specimen collections suggest that several species may exhibit multiple flowering peaks throughout the year (e.g., *Tabebuia aurea* (Bignoniaceae), *Hirtella racemosa* (Chrysobalanaceae), *Byrsonima crassifolia* (Malpighiaceae) and *Mimosa somnians* (Fabaceae); Appendix S2), such patterns cannot always be reliably inferred from herbarium specimens, especially for taxa and phenological stages that have not been well collected. Indeed, the discrepancies between herbarium specimen versus field observational records derived phenological estimates tended to be larger for fruits (Table S2; Appendix S1). Finally, herbarium specimens often comprise only part of a larger plant, and thus may not always reflect the general phenological stage of an entire individual. This issue may be exacerbated in tropical ecosystems due to the comparative abundance of tree species, though reproductive materials are often prioritized for collection.

Nonetheless, our study shows that tropical plant reproductive phenology inferred from herbarium records are widely congruent with field observations and demonstrates that herbarium specimens can be effectively used to assess patterns and mechanisms of plant phenological responses in the tropics. In particular, with theoretical and methodological advances that have made collections-based phenological studies increasingly efficient, herbarium specimens are positioned to be a vital resource for closing the gap in our phenological knowledge in tropical biomes (Davis and Ellison 2018; Davis et al. 2022). Such efforts will be critical to enhance our ability to predict how plant assemblages in the tropics will respond to an increasingly changing climate and implement mitigation strategies.

## Author Contributions

CCD conceived the initial idea for the project which was refined with discussions with the entire author group; CCD supervised the study; CCD, DSP, GML, DRT, RCA, and RKBM collected data; AME and DSP analyzed the data; DSP and GML drafted the first version of the manuscript and all authors contributed significantly to subsequent revisions.

## Acknowledgements

We thank Nádia Roque for hosting part of this research in her lab at the Universidade Federal da Bahia, Brazil. Funding for this research was provided by a Climate Change Solutions Fund grant from Harvard University. We additionally acknowledge funding from the U.S. National Science Foundation (NSF) grant NSF-DEB 1754584 to CCD, DSP, and AME, and from the ADBC program of the NSF to CCD (awards 1208835 [NEVP], 1702322 [SoRo], 1802209 [PoE], and 1902078 [TORCH]).

