## Supplementary material for "Herbarium records provide reliable phenology estimates in the understudied tropics": Table S2

**Table S2**. Results of linear models analyzing the effect of data source (field observation or herbarium specimen) on inferred flowering and fruiting periods.

| phenophase | scale | Estimate | Std. Error | df | t value | Pr(>\|t\|) | effect |
|---|---|---|---|---|---|---|---|
| flowering | municipality | -39.9526 | 22.4362 | 6.2345 | -1.7807 | 0.1234 | source-specimen |
| flowering | state | -0.4078 | 18.2277 | 22.4857 | -0.0224 | 0.9823 | source-specimen |
| flowering | biome | 72.3333 | 17.9472 | 25.5897 | 4.0303 | 0.0004 | source-specimen |
| flowering | country | 87.1818 | 16.9156 | 21.0000 | 5.1539 | 4.17E-05 | source-specimen |
| fruiting | municipality | -14.6988 | 10.4664 | 5.0500 | -1.4044 | 0.2186 | source-specimen |
| fruiting | state | 6.6518 | 23.8456 | 17.0959 | 0.2790 | 0.7836 | source-specimen |
| fruiting | biome | 52.6842 | 18.2941 | 18.0000 | 2.8799 | 0.0100 | source-specimen |
| fruiting | country | 83.2105 | 19.8293 | 18.0000 | 4.1963 | 0.0005 | source-specimen |
