## Supplementary material for "Herbarium records provide reliable phenology estimates in the understudied tropics": Fig. S1

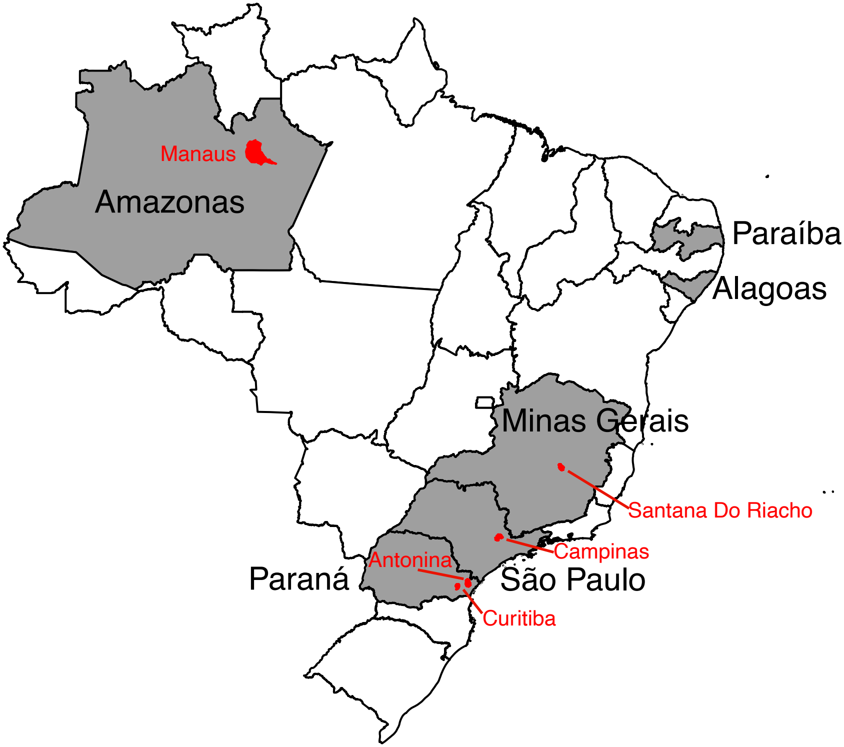


Figure S1. States and municipalities with both direct field survey and specimen dervided phenological information. Relevant states and municipalities are shaded in grey and red, respectively.
