## Appendix S2 for "Herbarium records provide reliable phenology estimates in the understudied tropics"

The following figures represent histogram of flowering and fruiting specimens collected for each species examined across six tropical biomes. Circular diagrams depict the time of collection in angles and sampling intensity in height. Empty spaces indicate the absence of specimens collected from the corresponding biome.

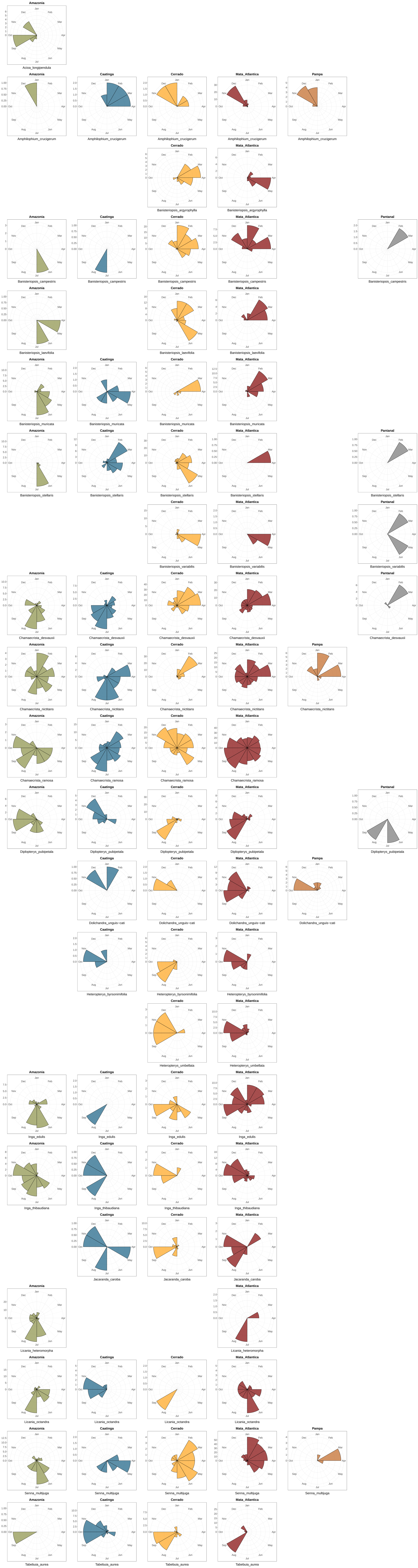
Amazonia
Acioa_longipendula
Amazonia
Amphilophium_crucigerum
Caatinga
Amphilophium_crucigerum
Cerrado
Amphilophium_crucigerum
Mata_Atlantica
Amphilophium_crucigerum
Pampa
Amphilophium_crucigerum
Cerrado
Banisteriopsis_argyrophylla
Mata_Atlantica
Banisteriopsis_argyrophylla
Amazonia
Banisteriopsis_campestris
Caatinga
Banisteriopsis_campestris
Cerrado
Banisteriopsis_campestris
Mata_Atlantica
Banisteriopsis_campestris
Pantanal
Banisteriopsis_campestris
Amazonia
Banisteriopsis_laevifolia
Cerrado
Banisteriopsis_laevifolia
Mata_Atlantica
Banisteriopsis_laevifolia
Amazonia
Banisteriopsis_muricata
Caatinga
Banisteriopsis_muricata
Cerrado
Banisteriopsis_muricata
Mata_Atlantica
Banisteriopsis_muricata
Amazonia
Banisteriopsis_stellaris
Caatinga
Banisteriopsis_stellaris
Cerrado
Banisteriopsis_stellaris
Mata_Atlantica
Banisteriopsis_stellaris
Pantanal
Banisteriopsis_stellaris
Cerrado
Banisteriopsis_variabilis
Mata_Atlantica
Banisteriopsis_variabilis
Pantanal
Banisteriopsis_variabilis
Amazonia
Chamaecrista_desvauxii
Caatinga
Chamaecrista_desvauxii
Cerrado
Chamaecrista_desvauxii
Mata_Atlantica
Chamaecrista_desvauxii
Pantanal
Chamaecrista_desvauxii
Amazonia
Chamaecrista_nictitans
Caatinga
Chamaecrista_nictitans
Cerrado
Chamaecrista_nictitans
Mata_Atlantica
Chamaecrista_nictitans
Pampa
Chamaecrista_nictitans
Amazonia
Chamaecrista_ramosa
Caatinga
Chamaecrista_ramosa
Cerrado
Chamaecrista_ramosa
Mata_Atlantica
Chamaecrista_ramosa
Amazonia
Diplopterys_pubipetala
Caatinga
Diplopterys_pubipetala
Cerrado
Diplopterys_pubipetala
Mata_Atlantica
Diplopterys_pubipetala
Pantanal
Diplopterys_pubipetala
Caatinga
Dolichandra_unguis−cati
Cerrado
Dolichandra_unguis−cati
Mata_Atlantica
Dolichandra_unguis−cati
Pampa
Dolichandra_unguis−cati
Caatinga
Heteropterys_byrsonimifolia
Cerrado
Heteropterys_byrsonimifolia
Mata_Atlantica
Heteropterys_byrsonimifolia
Cerrado
Heteropterys_umbellata
Mata_Atlantica
Heteropterys_umbellata
Amazonia
Inga_edulis
Caatinga
Inga_edulis
Cerrado
Inga_edulis
Mata_Atlantica
Inga_edulis
Amazonia
Inga_thibaudiana
Caatinga
Inga_thibaudiana
Cerrado
Inga_thibaudiana
Mata_Atlantica
Inga_thibaudiana
Caatinga
Jacaranda_caroba
Cerrado
Jacaranda_caroba
Mata_Atlantica
Jacaranda_caroba
Amazonia
Licania_heteromorpha
Mata_Atlantica
Licania_heteromorpha
Amazonia
Licania_octandra
Caatinga
Licania_octandra
Cerrado
Licania_octandra
Mata_Atlantica
Licania_octandra
Amazonia
Senna_multijuga
Caatinga
Senna_multijuga
Cerrado
Senna_multijuga
Mata_Atlantica
Senna_multijuga
Pampa
Senna_multijuga
Amazonia
Tabebuia_aurea
Caatinga
Tabebuia_aurea
Cerrado
Tabebuia_aurea
Mata_Atlantica
Tabebuia_aurea

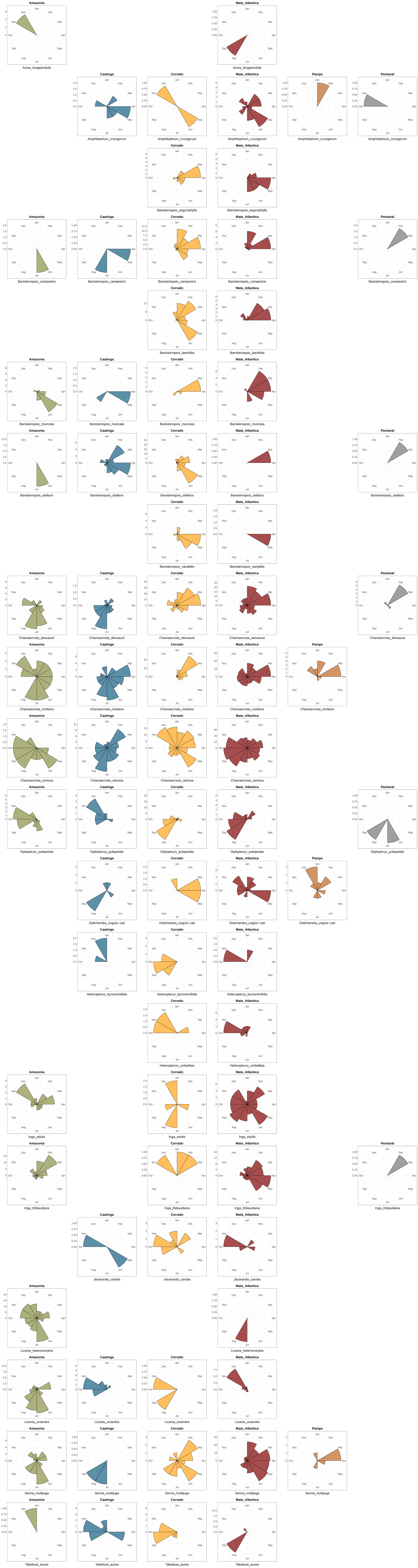
Amazonia
Acioa_longipendula
Mata_Atlantica
Acioa_longipendula
Caatinga
Amphilophium_crucigerum
Cerrado
Amphilophium_crucigerum
Mata_Atlantica
Amphilophium_crucigerum
Pampa
Amphilophium_crucigerum
Pantanal
Amphilophium_crucigerum
Cerrado
Banisteriopsis_argyrophylla
Mata_Atlantica
Banisteriopsis_argyrophylla
Amazonia
Banisteriopsis_campestris
Caatinga
Banisteriopsis_campestris
Cerrado
Banisteriopsis_campestris
Mata_Atlantica
Banisteriopsis_campestris
Pantanal
Banisteriopsis_campestris
Cerrado
Banisteriopsis_laevifolia
Mata_Atlantica
Banisteriopsis_laevifolia
Amazonia
Banisteriopsis_muricata
Caatinga
Banisteriopsis_muricata
Cerrado
Banisteriopsis_muricata
Mata_Atlantica
Banisteriopsis_muricata
Amazonia
Banisteriopsis_stellaris
Caatinga
Banisteriopsis_stellaris
Cerrado
Banisteriopsis_stellaris
Mata_Atlantica
Banisteriopsis_stellaris
Pantanal
Banisteriopsis_stellaris
Cerrado
Banisteriopsis_variabilis
Mata_Atlantica
Banisteriopsis_variabilis
Amazonia
Chamaecrista_desvauxii
Caatinga
Chamaecrista_desvauxii
Cerrado
Chamaecrista_desvauxii
Mata_Atlantica
Chamaecrista_desvauxii
Pantanal
Chamaecrista_desvauxii
Amazonia
Chamaecrista_nictitans
Caatinga
Chamaecrista_nictitans
Cerrado
Chamaecrista_nictitans
Mata_Atlantica
Chamaecrista_nictitans
Pampa
Chamaecrista_nictitans
Amazonia
Chamaecrista_ramosa
Caatinga
Chamaecrista_ramosa
Cerrado
Chamaecrista_ramosa
Mata_Atlantica
Chamaecrista_ramosa
Amazonia
Diplopterys_pubipetala
Caatinga
Diplopterys_pubipetala
Cerrado
Diplopterys_pubipetala
Mata_Atlantica
Diplopterys_pubipetala
Pantanal
Diplopterys_pubipetala
Caatinga
Dolichandra_unguis−cati
Cerrado
Dolichandra_unguis−cati
Mata_Atlantica
Dolichandra_unguis−cati
Pampa
Dolichandra_unguis−cati
Caatinga
Heteropterys_byrsonimifolia
Cerrado
Heteropterys_byrsonimifolia
Mata_Atlantica
Heteropterys_byrsonimifolia
Cerrado
Heteropterys_umbellata
Mata_Atlantica
Heteropterys_umbellata
Amazonia
Inga_edulis
Cerrado
Inga_edulis
Mata_Atlantica
Inga_edulis
Amazonia
Inga_thibaudiana
Cerrado
Inga_thibaudiana
Mata_Atlantica
Inga_thibaudiana
Pantanal
Inga_thibaudiana
Caatinga
Jacaranda_caroba
Cerrado
Jacaranda_caroba
Mata_Atlantica
Jacaranda_caroba
Amazonia
Licania_heteromorpha
Mata_Atlantica
Licania_heteromorpha
Amazonia
Licania_octandra
Caatinga
Licania_octandra
Cerrado
Licania_octandra
Mata_Atlantica
Licania_octandra
Amazonia
Senna_multijuga
Caatinga
Senna_multijuga
Cerrado
Senna_multijuga
Mata_Atlantica
Senna_multijuga
Pampa
Senna_multijuga
Amazonia
Tabebuia_aurea
Caatinga
Tabebuia_aurea
Cerrado
Tabebuia_aurea
Mata_Atlantica
Tabebuia_aurea
