## Appendix S1 for "Herbarium records provide reliable phenology estimates in the understudied tropics"

Each following plot illustrates the flowering/fruiting period of a species as inferred by plot observations (thick/faint lines) and that inferred from herbarium specimens (thin/bold lines) collected in the same municipality (red), state (grey), biome (yellow), and country (green). The start and end of flowering and fruiting periods were inferred from a circular distribution of observation/collection dates of individuals in flower or fruit.

### *Acioa longipendula*

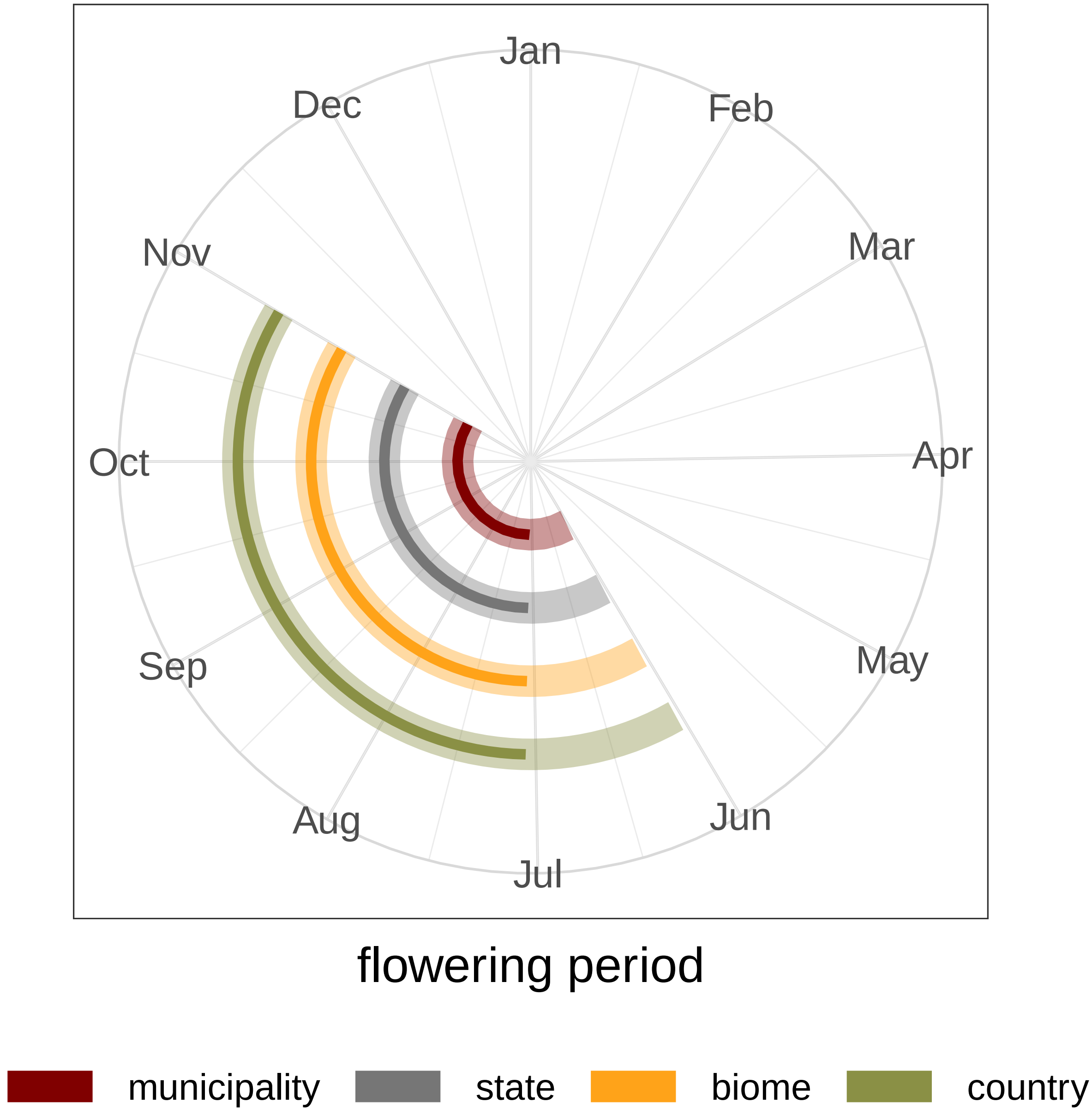

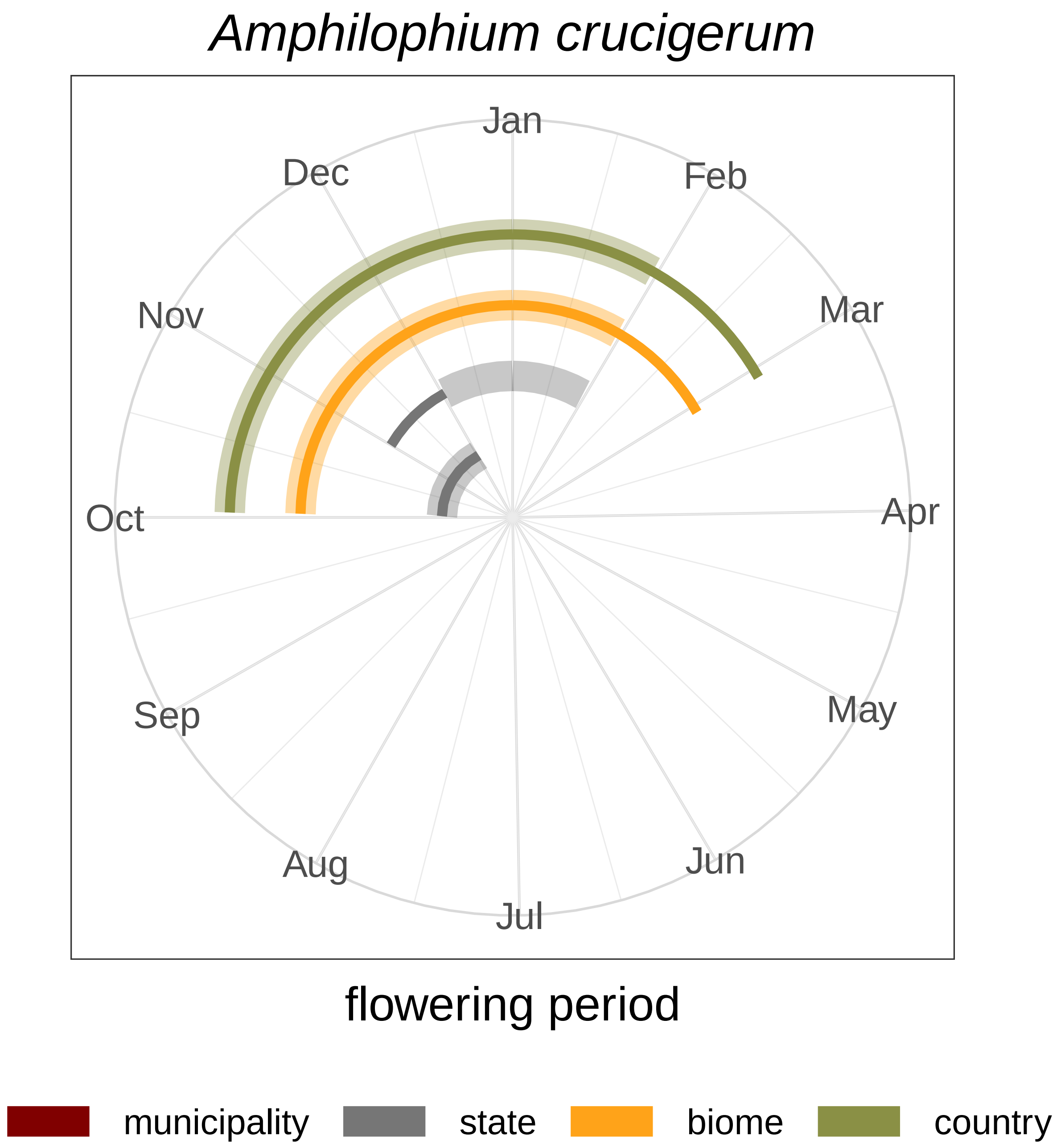

Amphilophium crucigerum
Jan
Feb
Mar
Apr
May
Jun
Jul
Aug
Sep
Oct
Nov
Dec
flowering period
municipality
state
biome
country

### *Banisteriopsis argyrophylla*

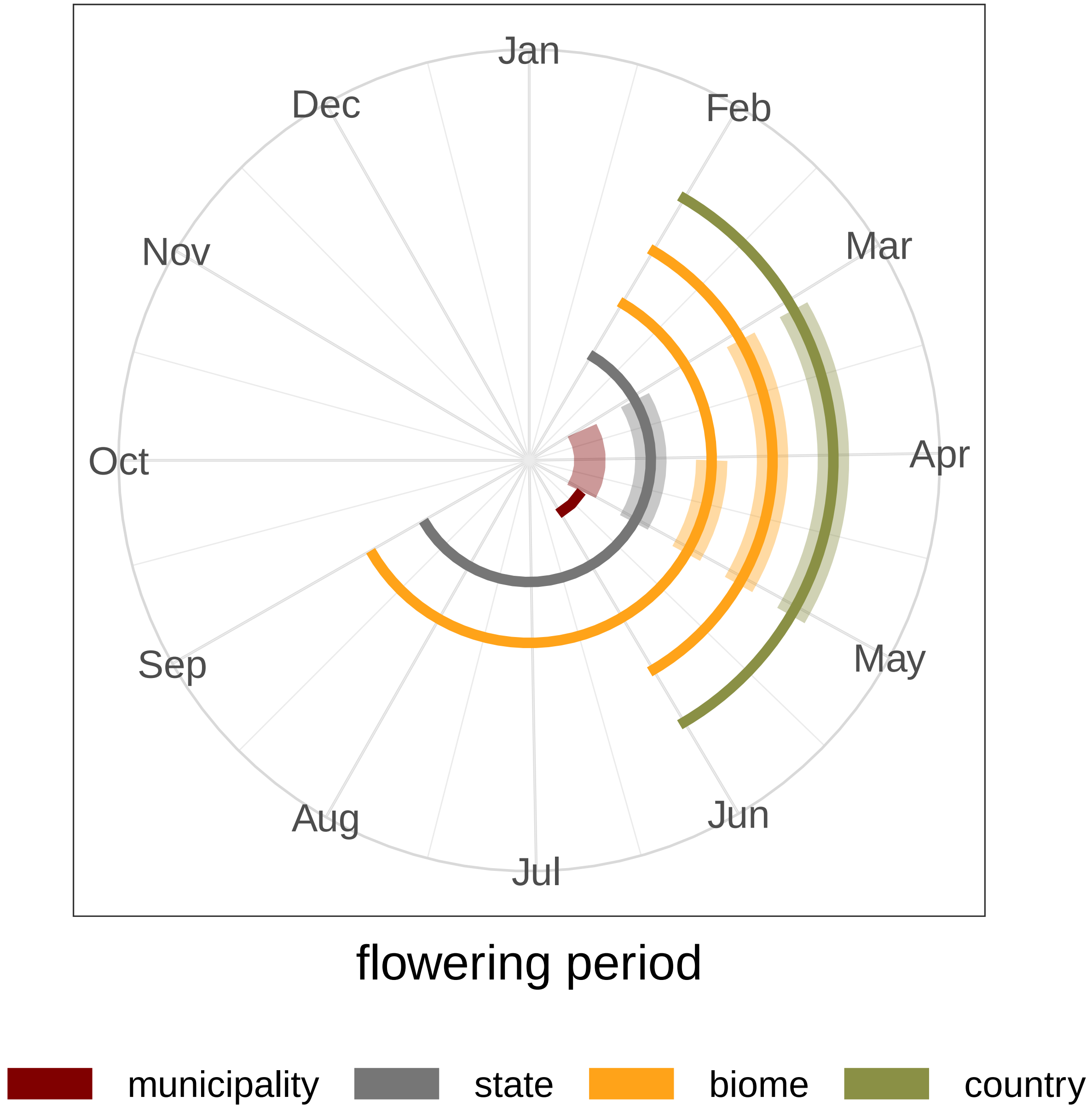

### *Banisteriopsis campestris*

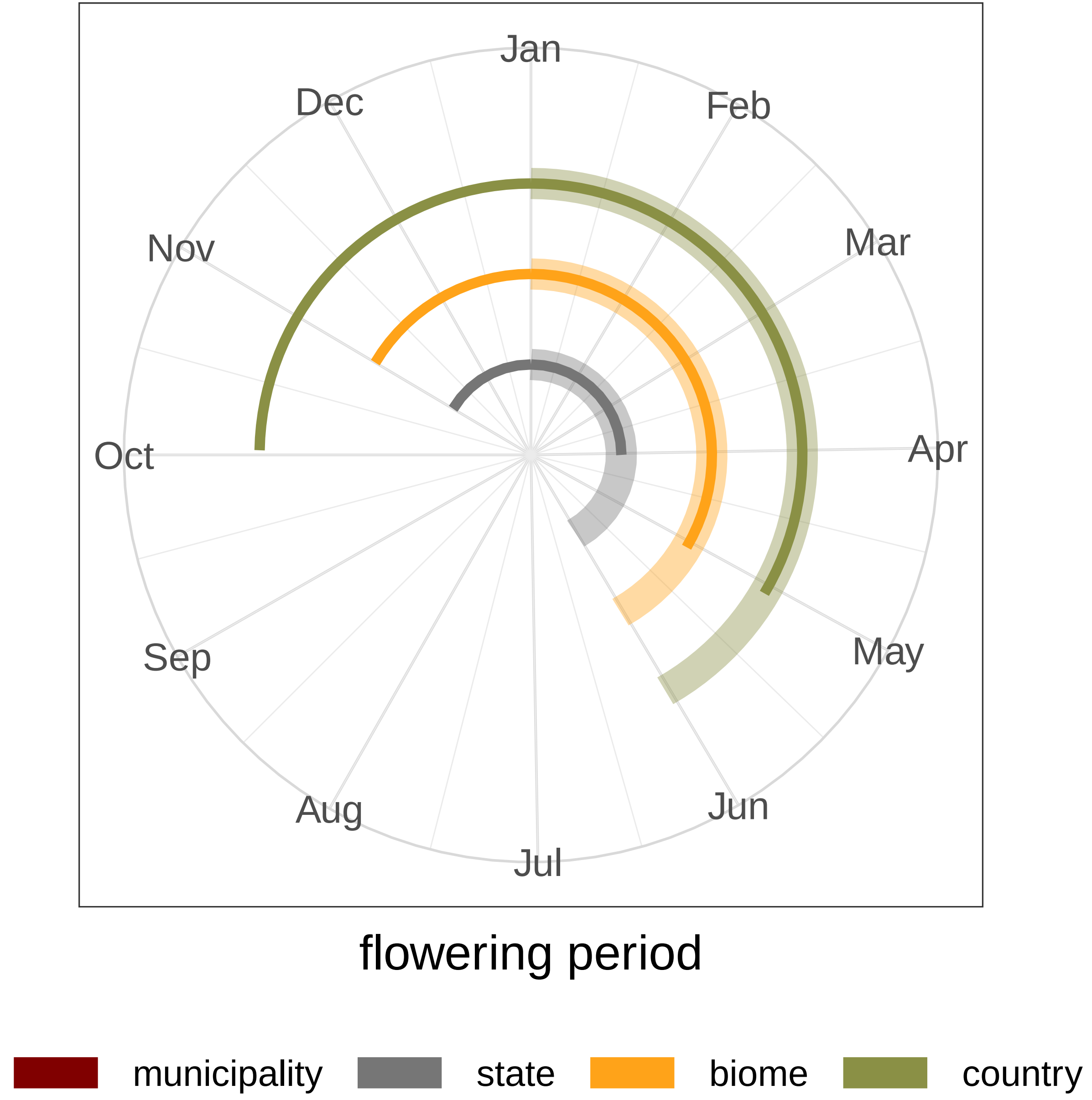

### *Banisteriopsis laevifolia*

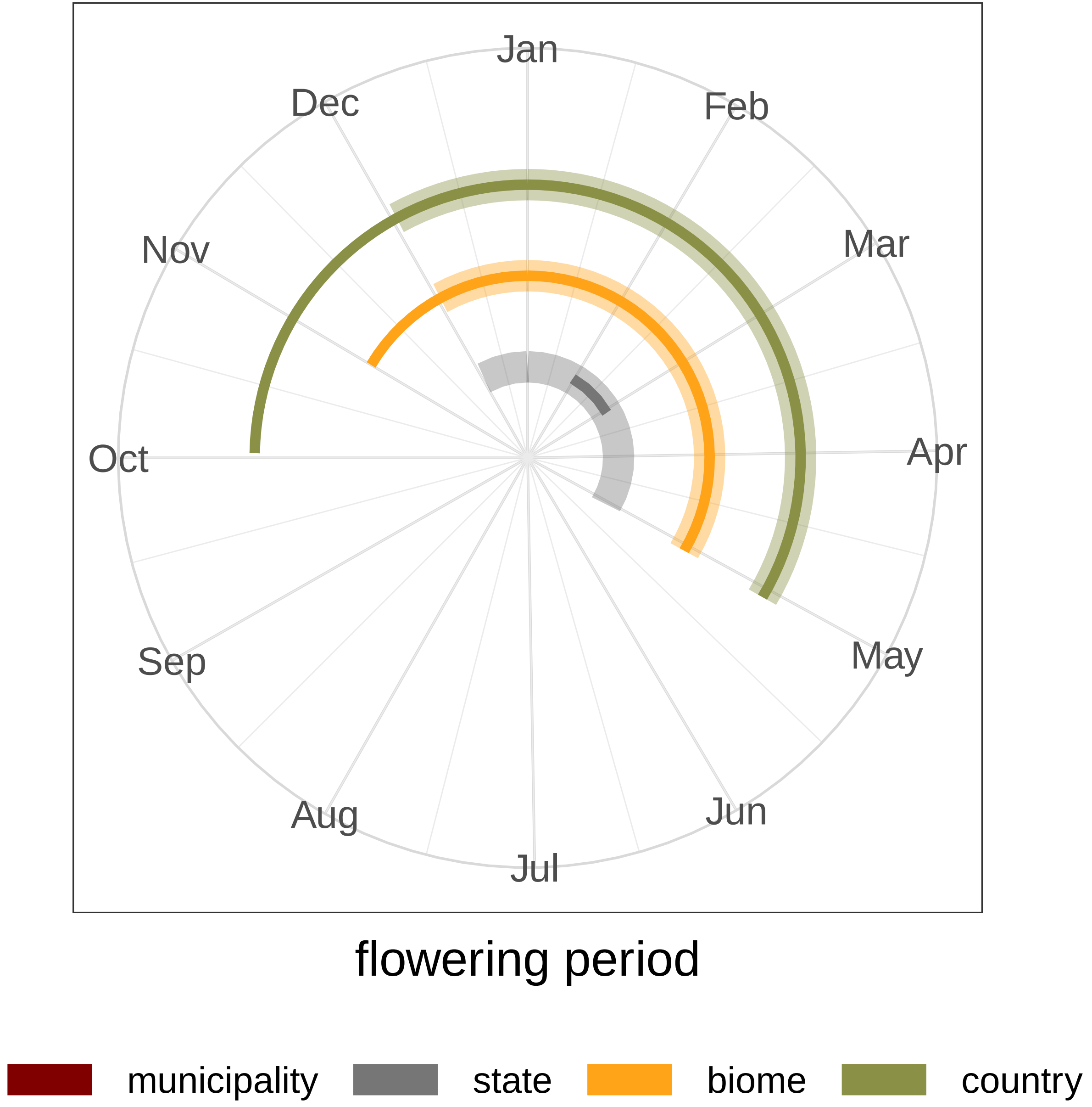

### *Banisteriopsis muricata*

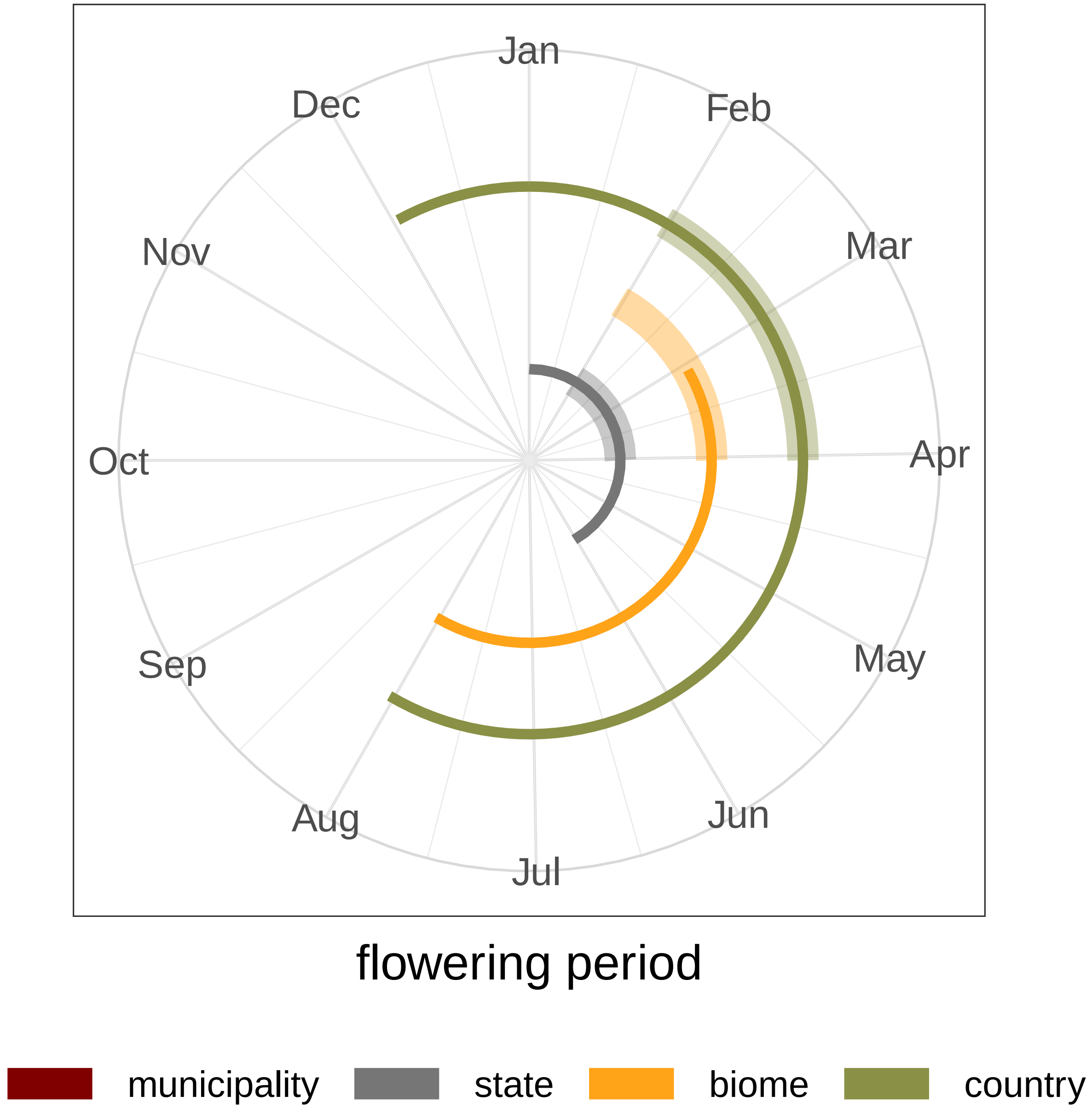

### *Banisteriopsis stellaris*

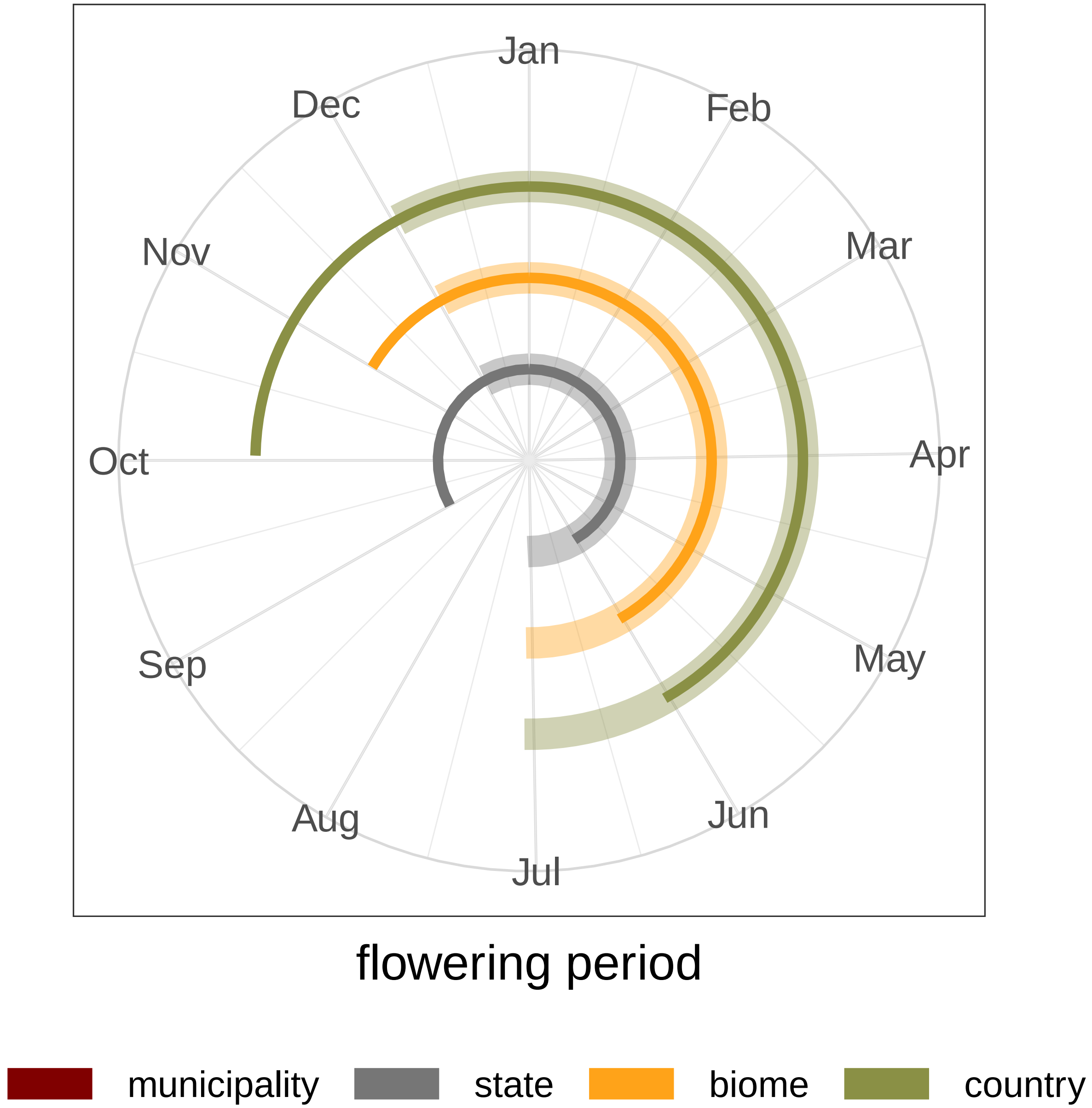

### *Banisteriopsis variabilis*

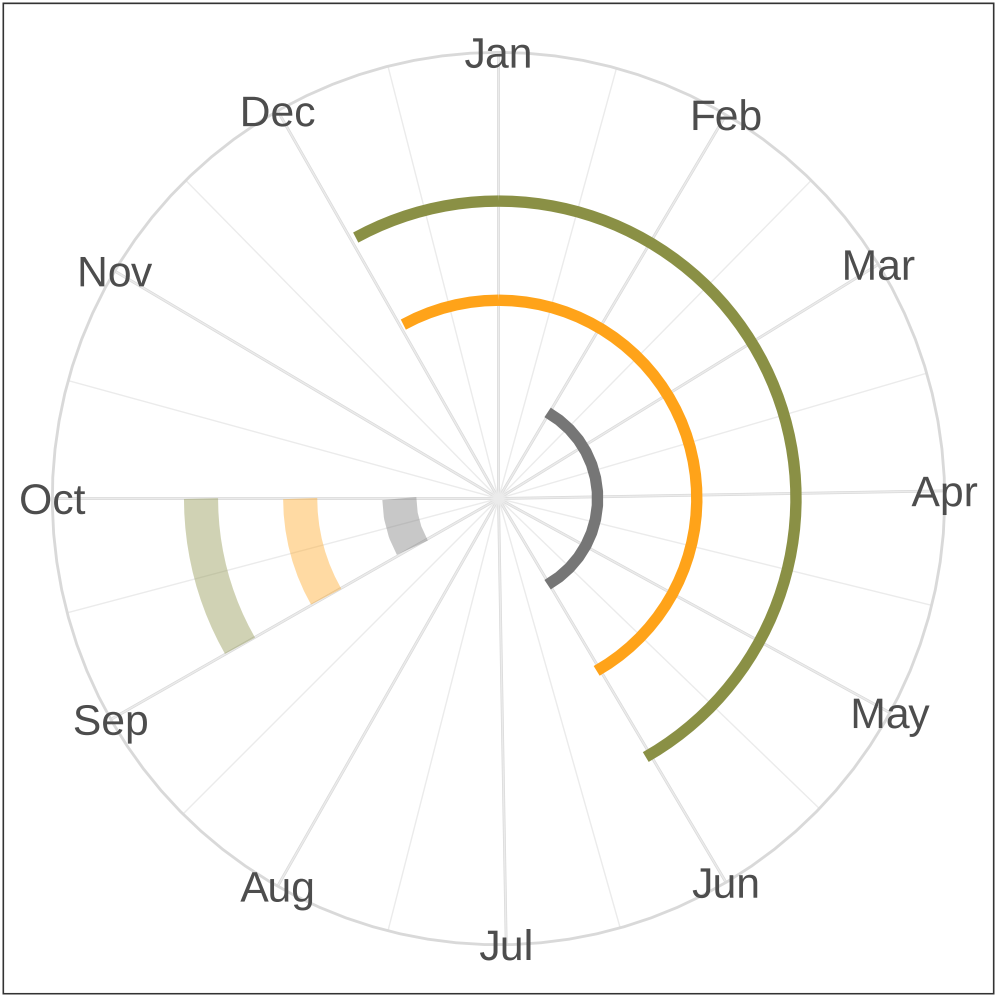


flowering period

### *Chamaecrista desvauxii*

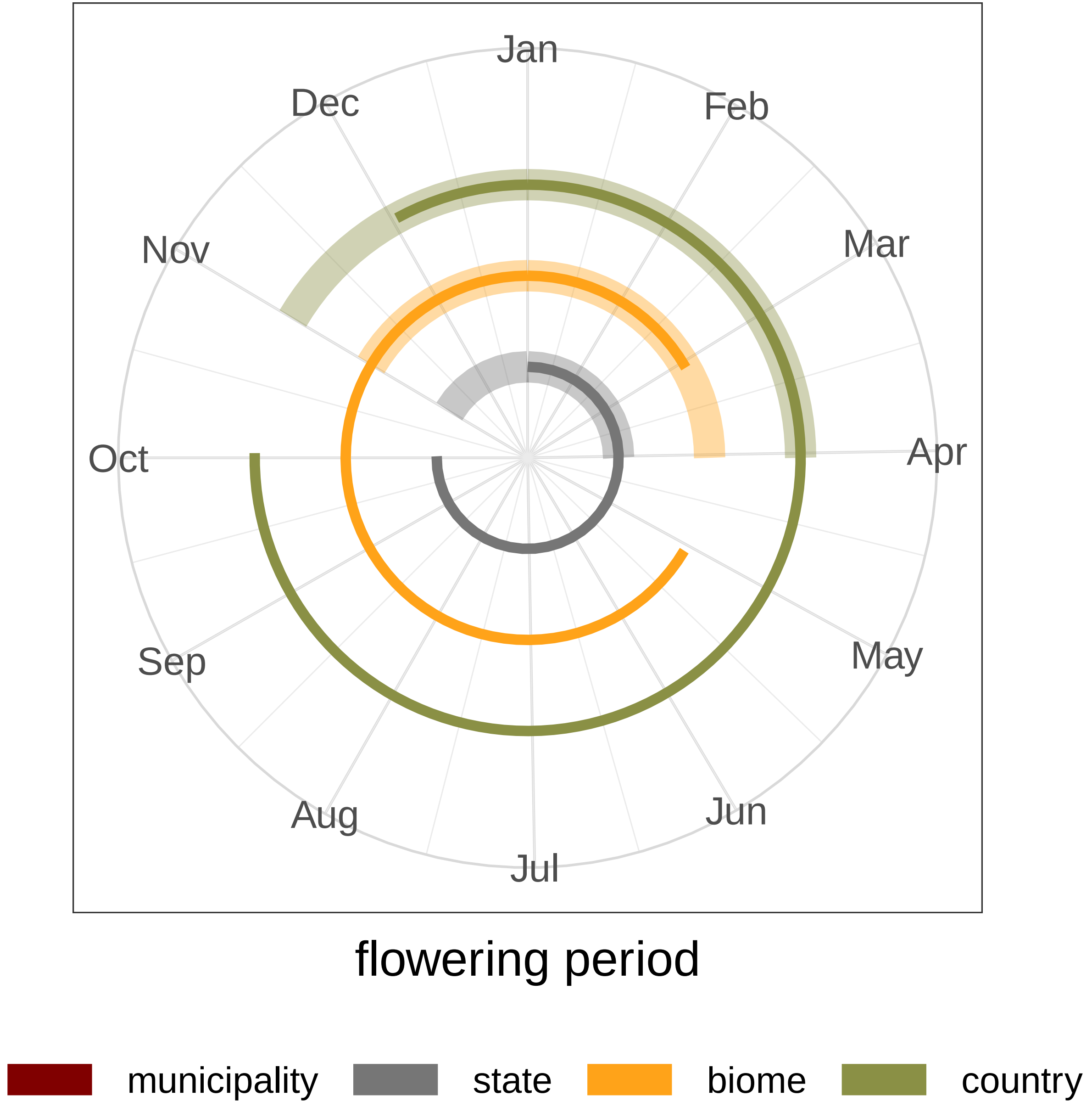

### *Chamaecrista nictitans*

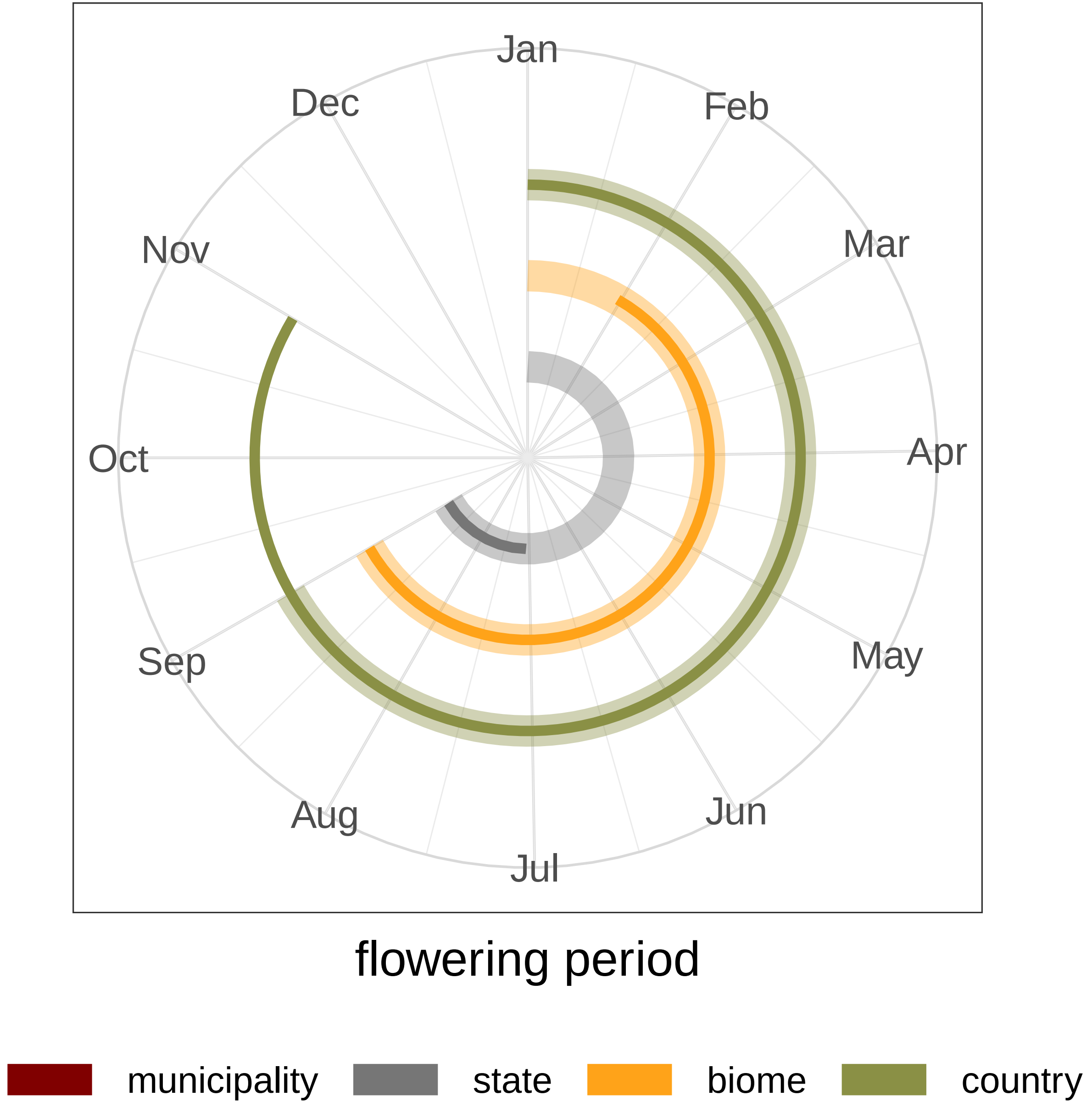

### *Chamaecrista ramosa*

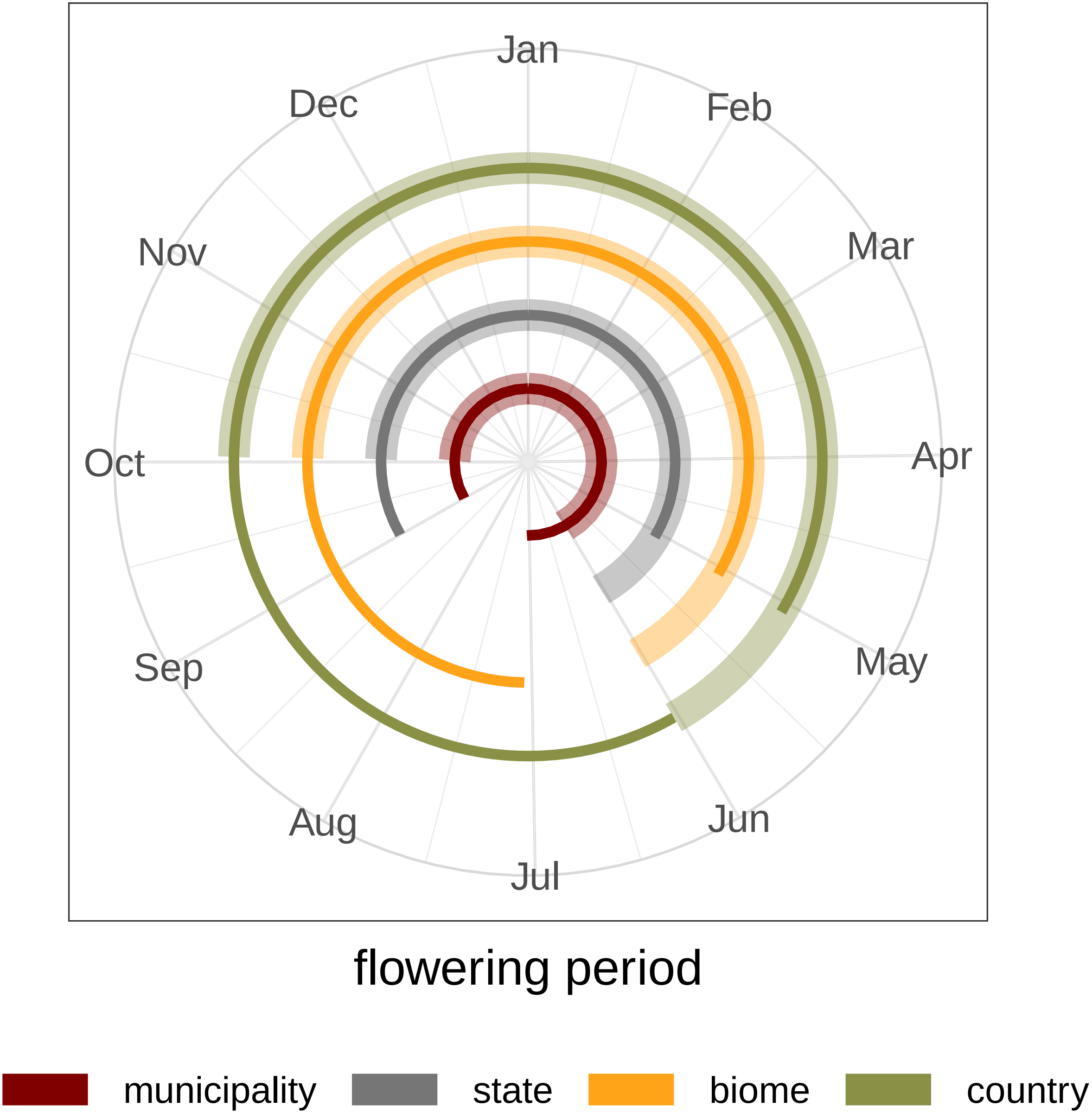

### *Diplopterys pubipetala*

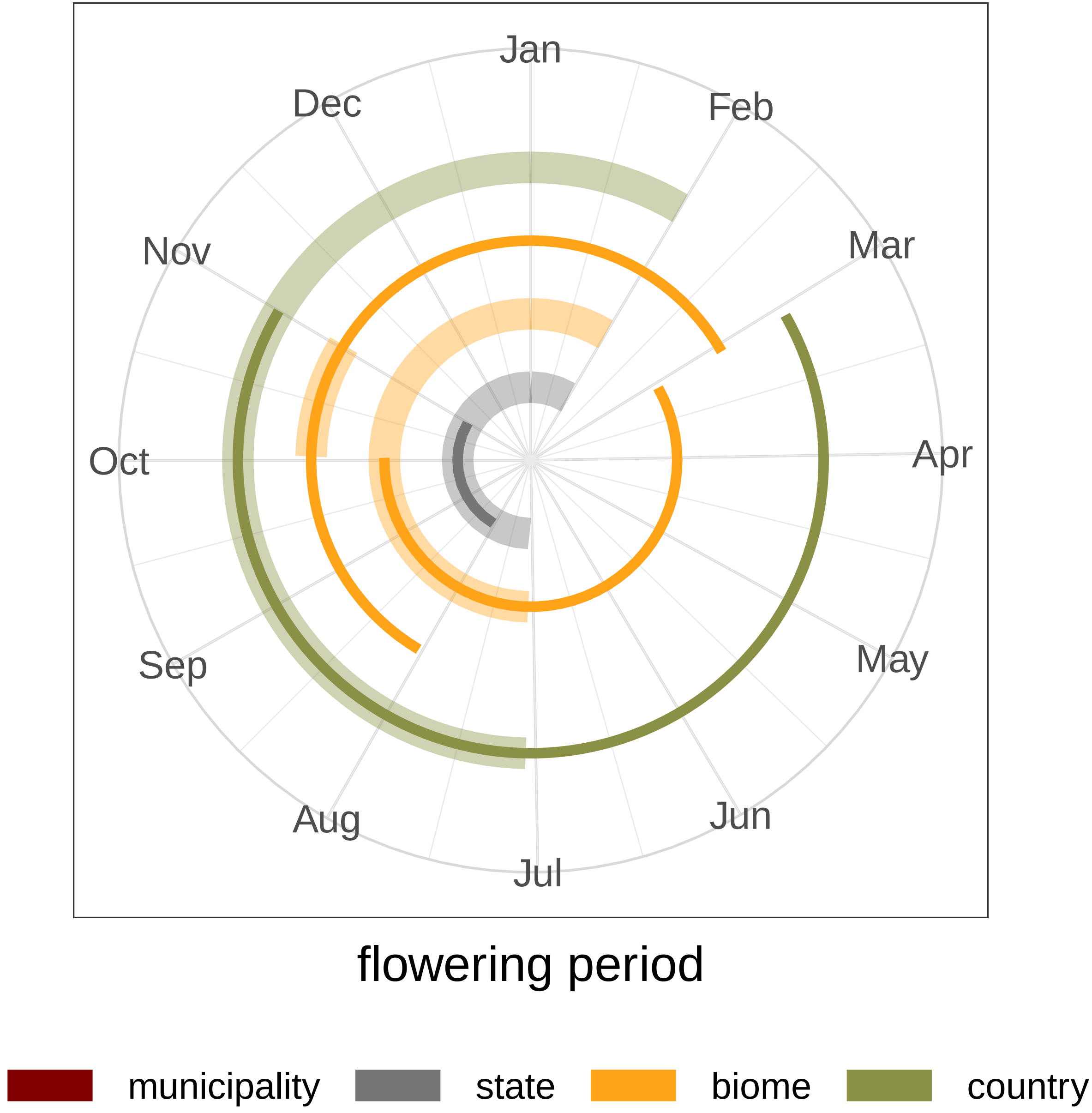

### *Dolichandra unguis−cati*

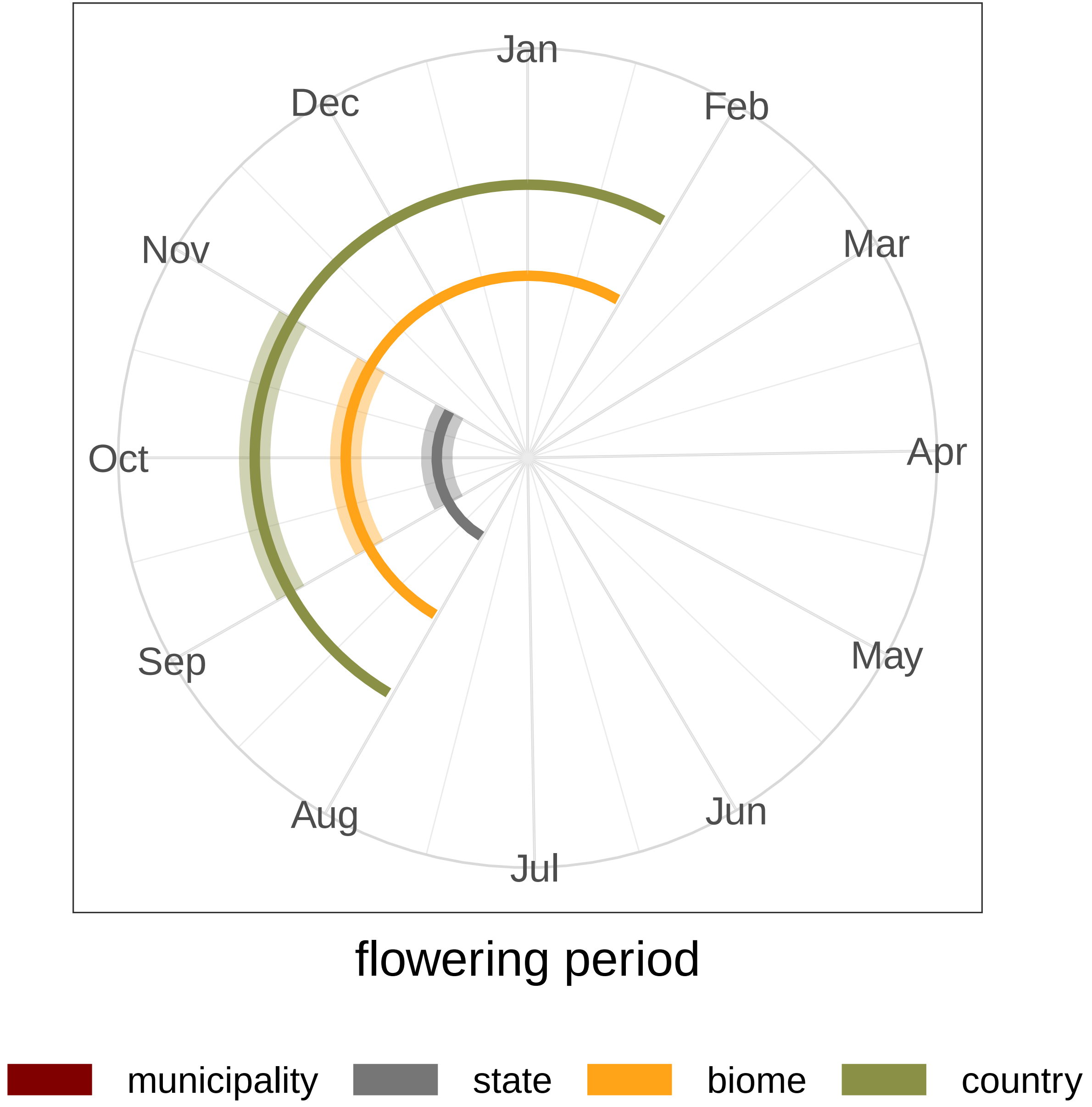

### *Heteropterys byrsonimifolia*

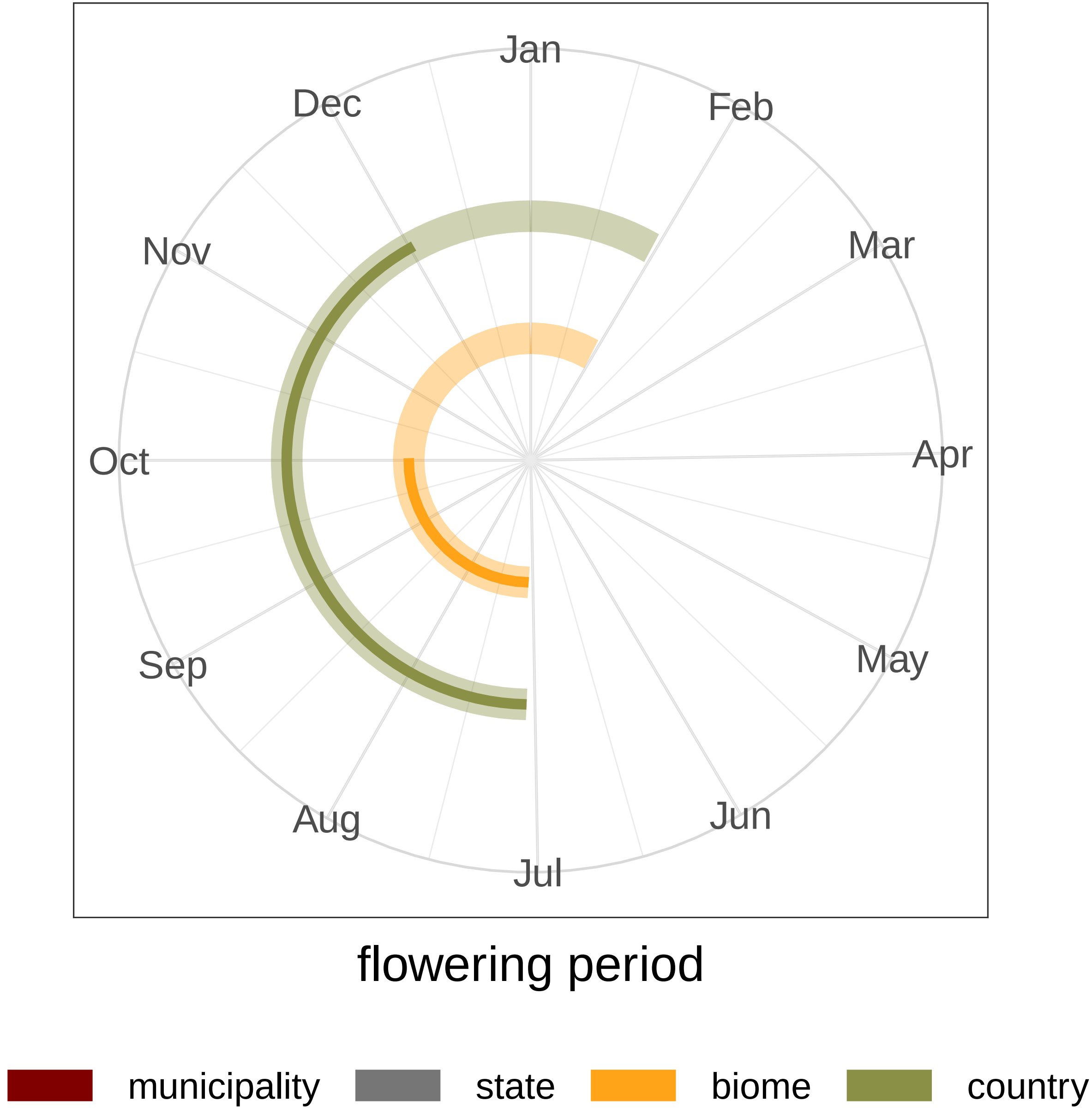

### *Heteropterys umbellata*

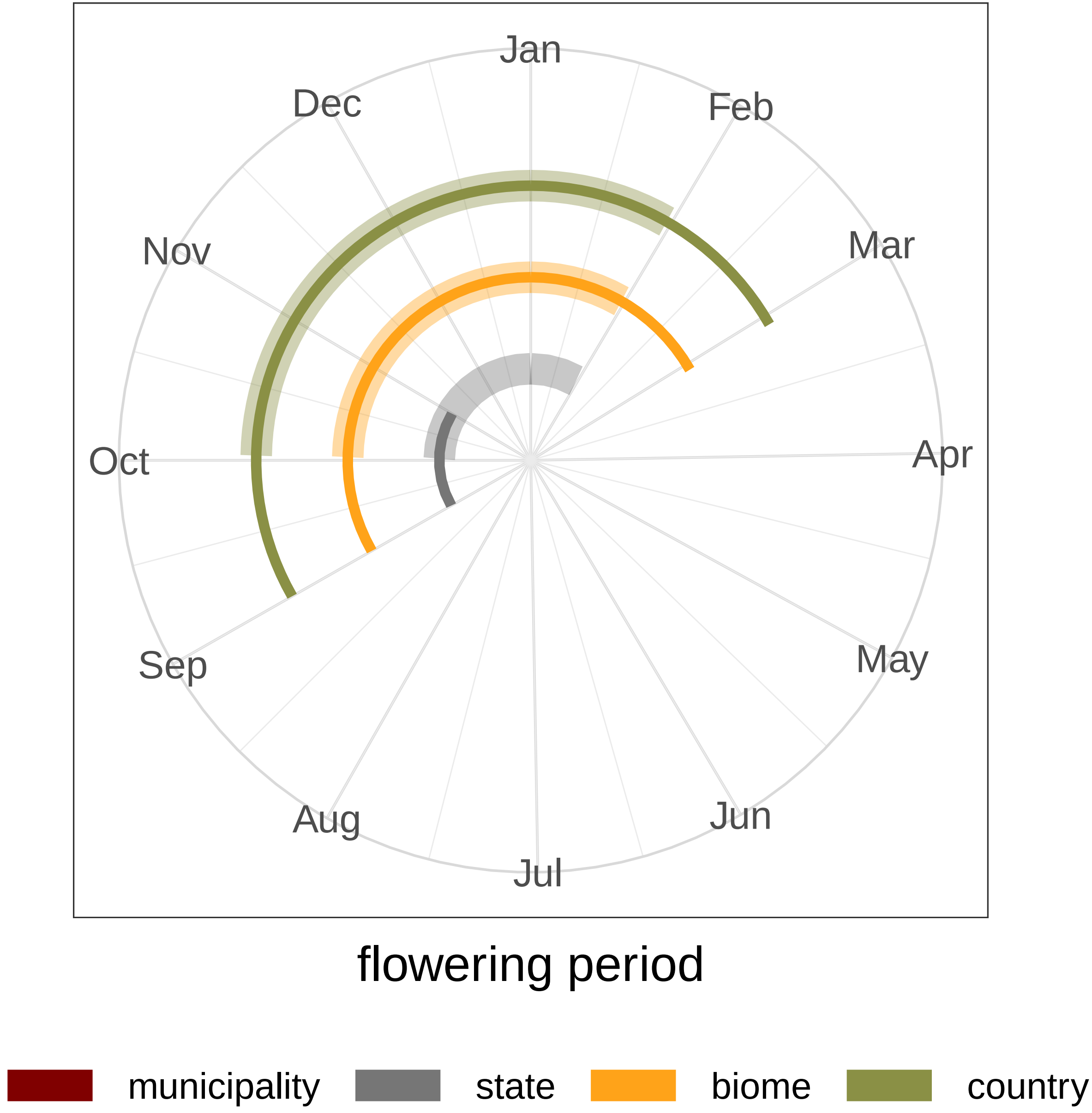

### *Inga edulis*

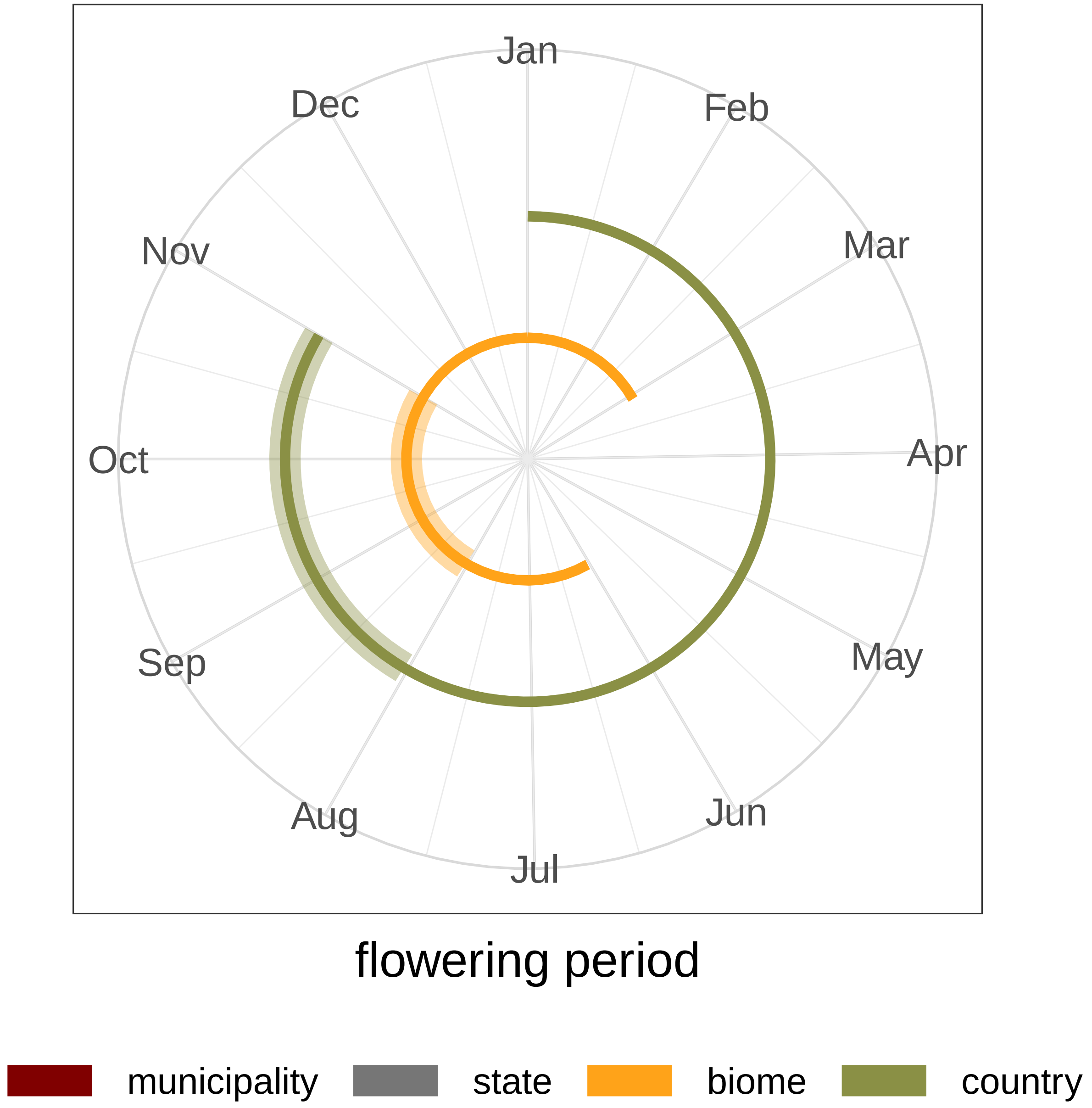
Jan
Feb
Mar
Apr
May
Jun
Jul
Aug
Sep
Oct
Nov
Dec
flowering period
municipality
state
biome
country

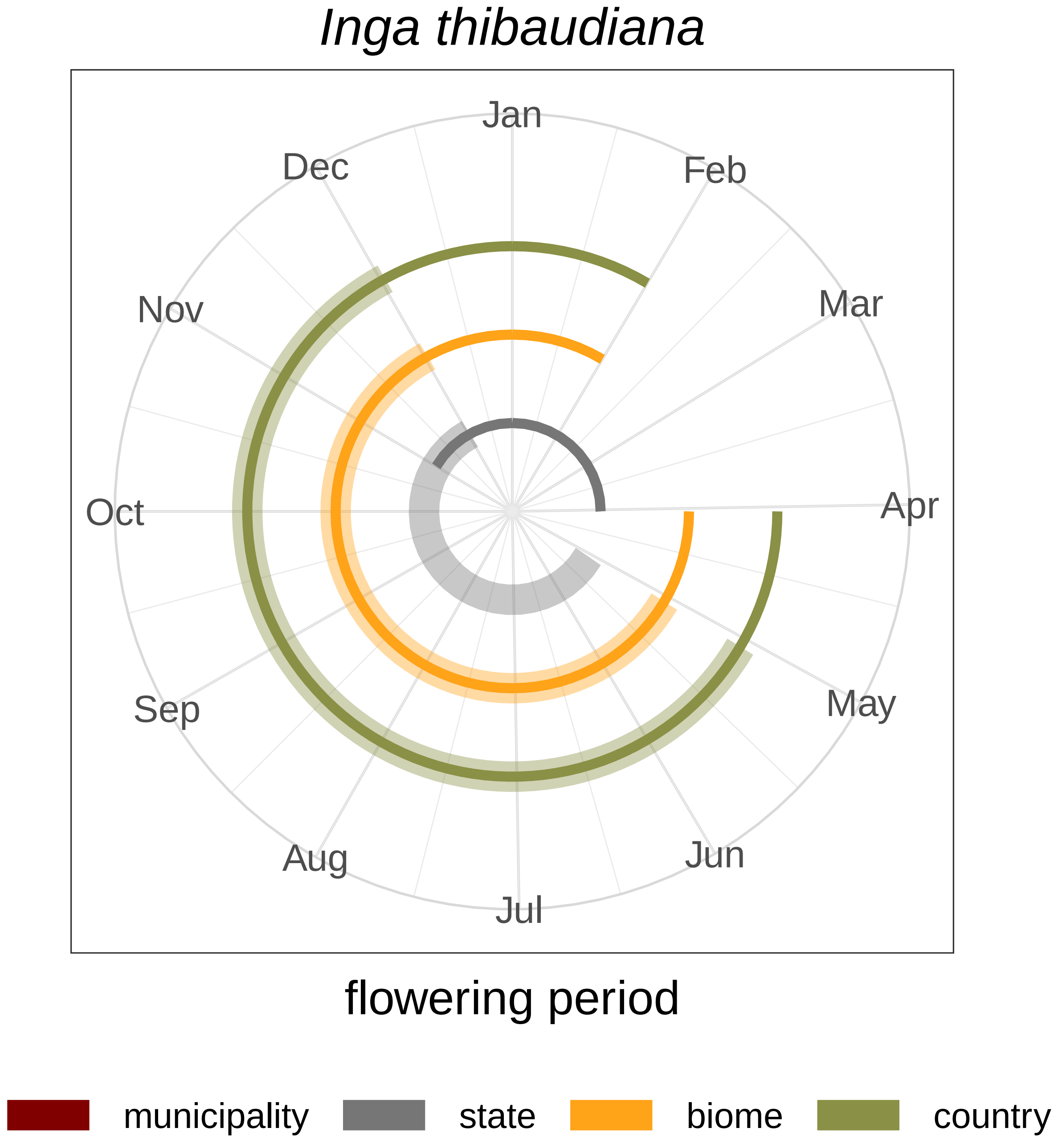

Inga thibaudiana
Jan
Feb
Mar
Apr
May
Jun
Jul
Aug
Sep
Oct
Nov
Dec
flowering period
municipality
state
biome
country

### *Jacaranda caroba*

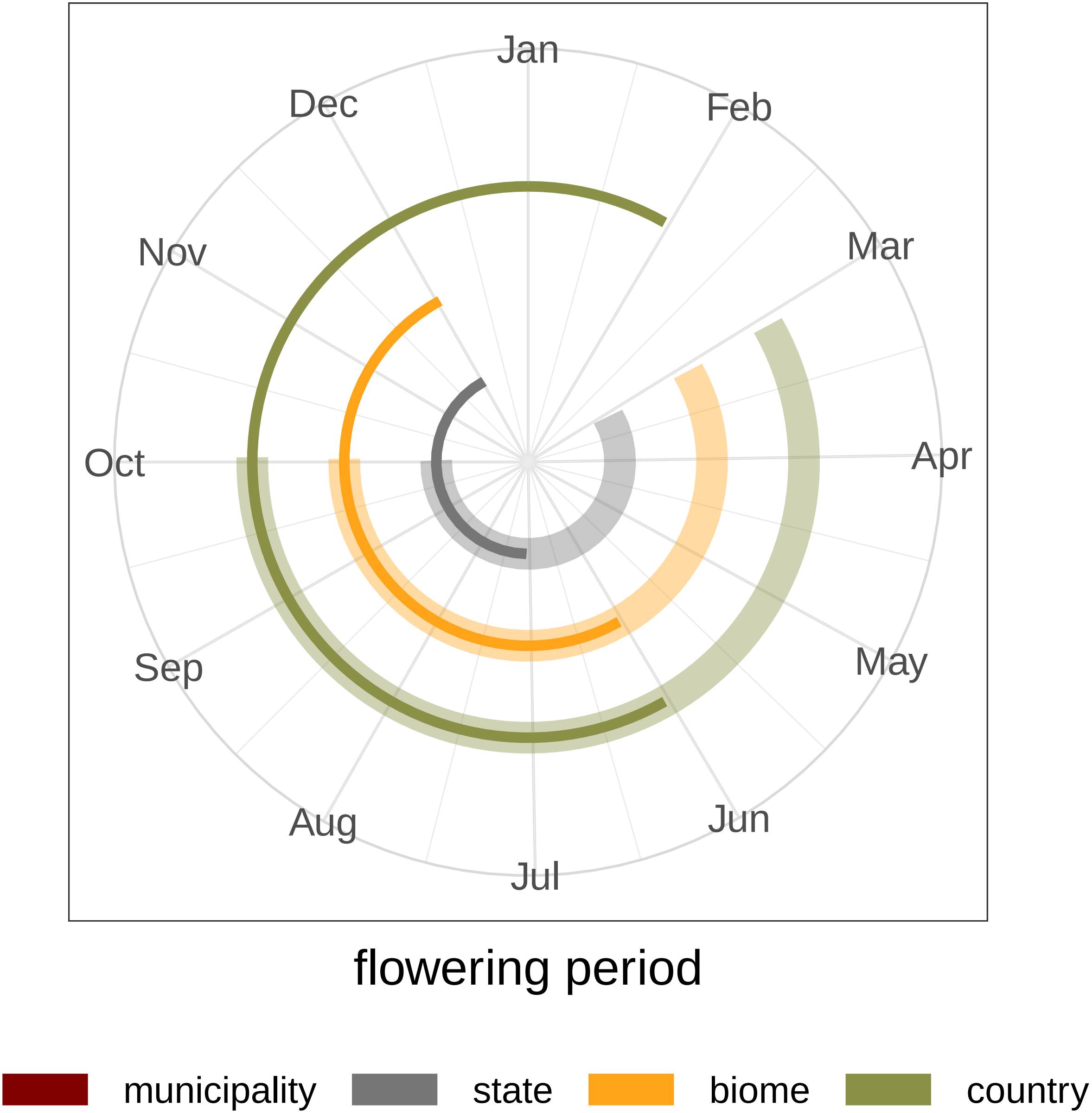

### *Licania heteromorpha*

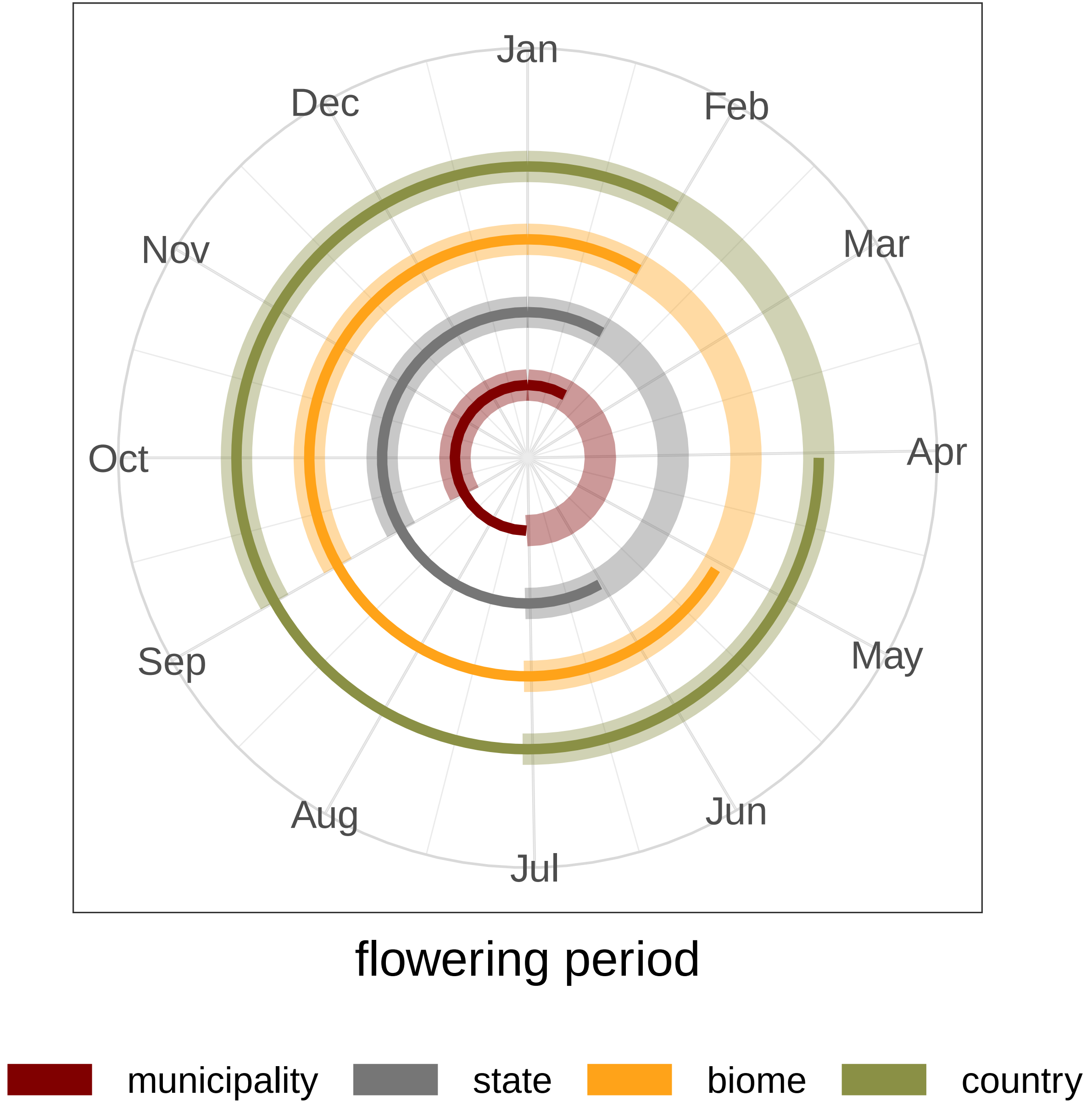

### *Licania octandra*

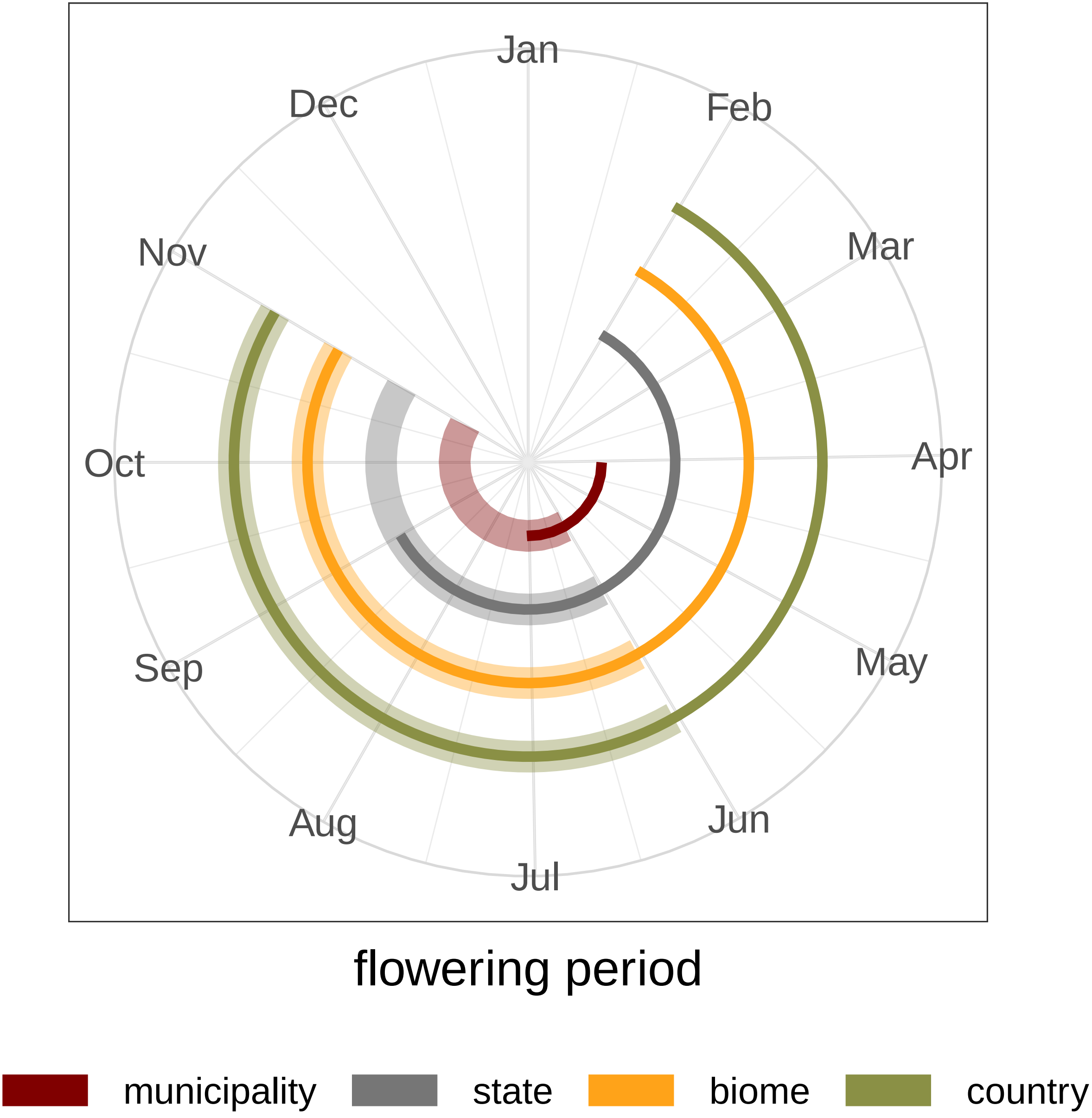

### *Senna multijuga*

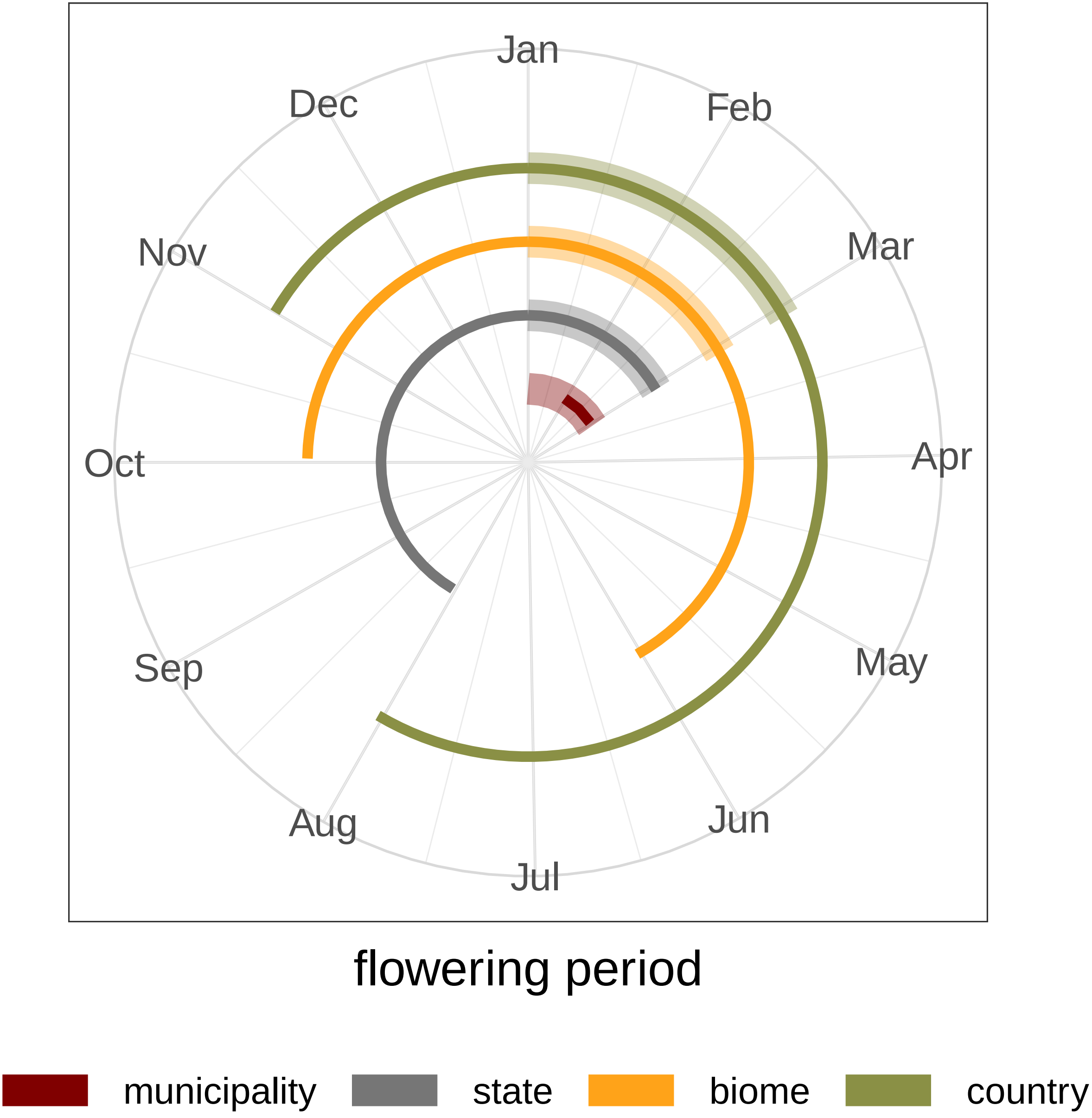

### *Tabebuia aurea*

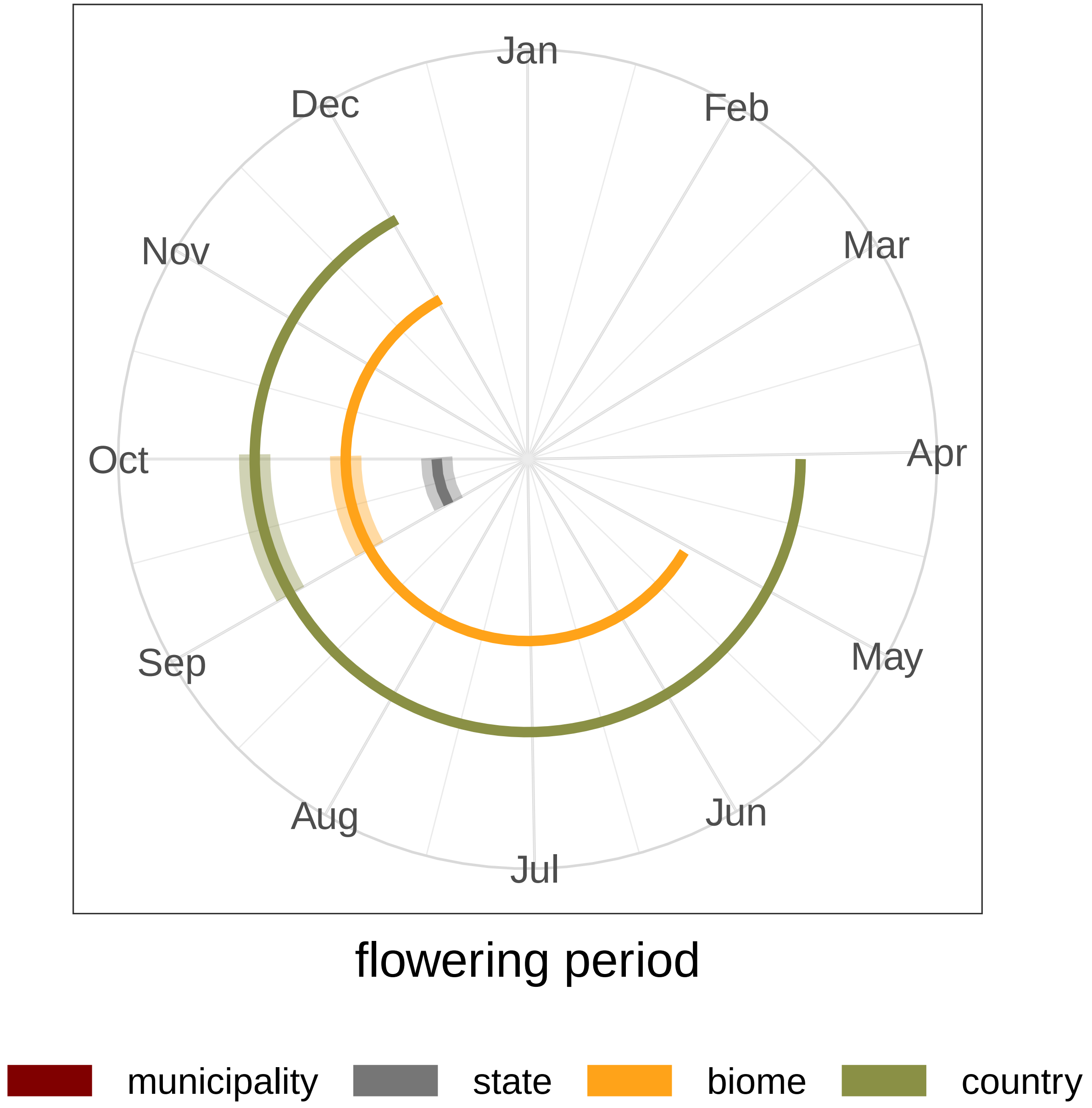

### *Acioa longipendula*

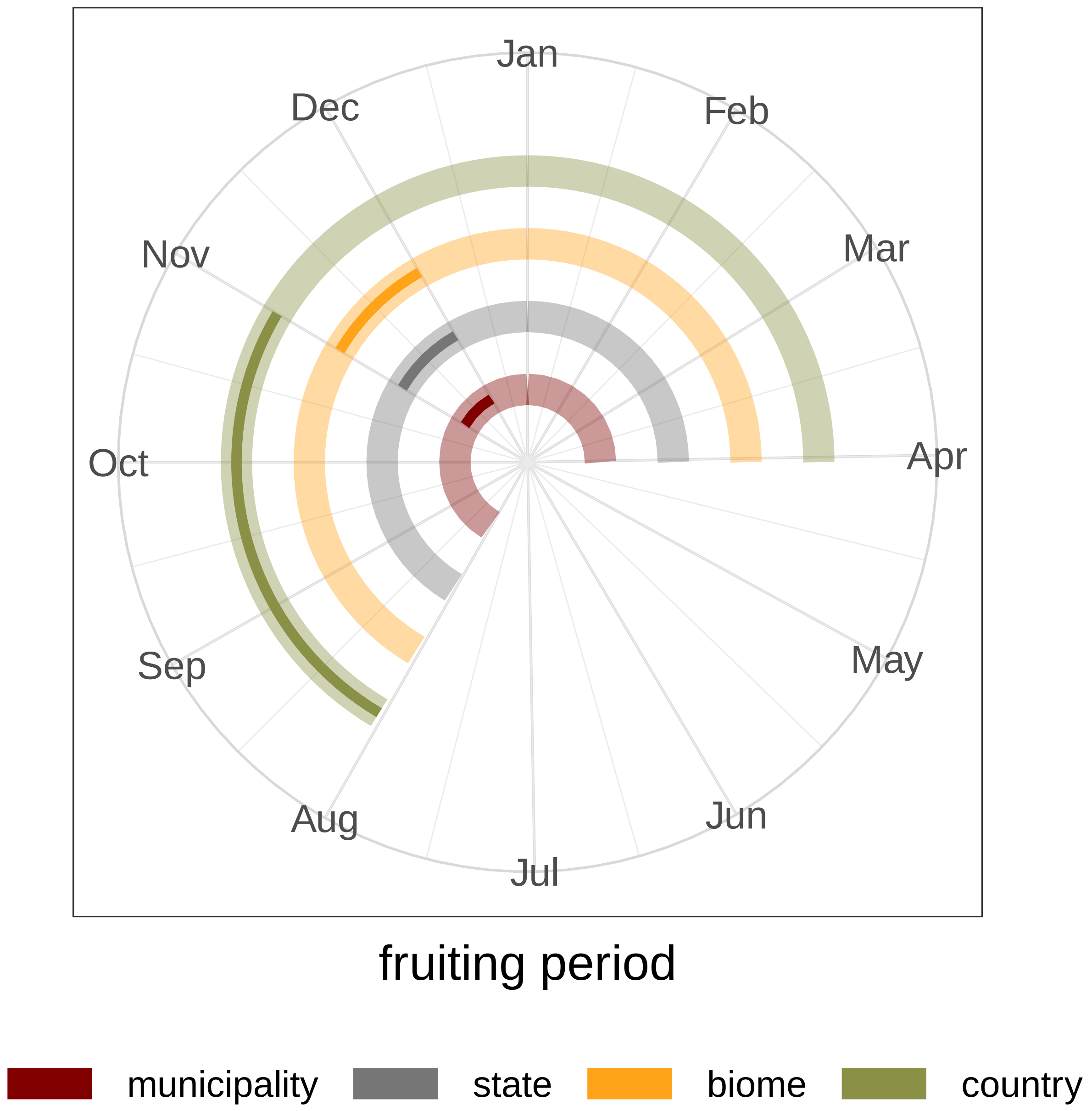

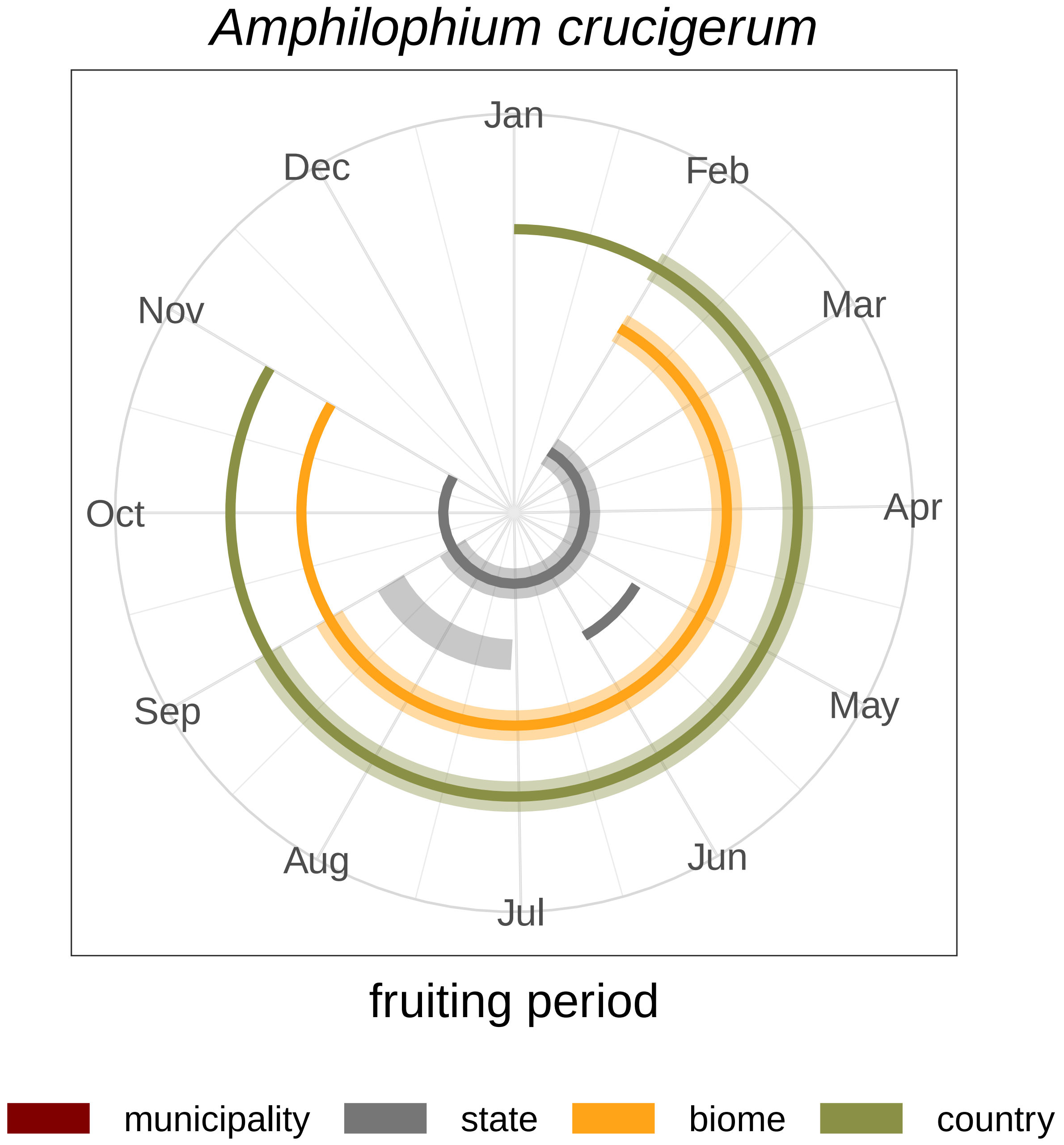

*Amphilophium crucigerum*
Jan
Feb
Mar
Apr
May
Jun
Jul
Aug
Sep
Oct
Nov
Dec
fruiting period
municipality
state
biome
country

### *Anemopaegma chamberlaynii*

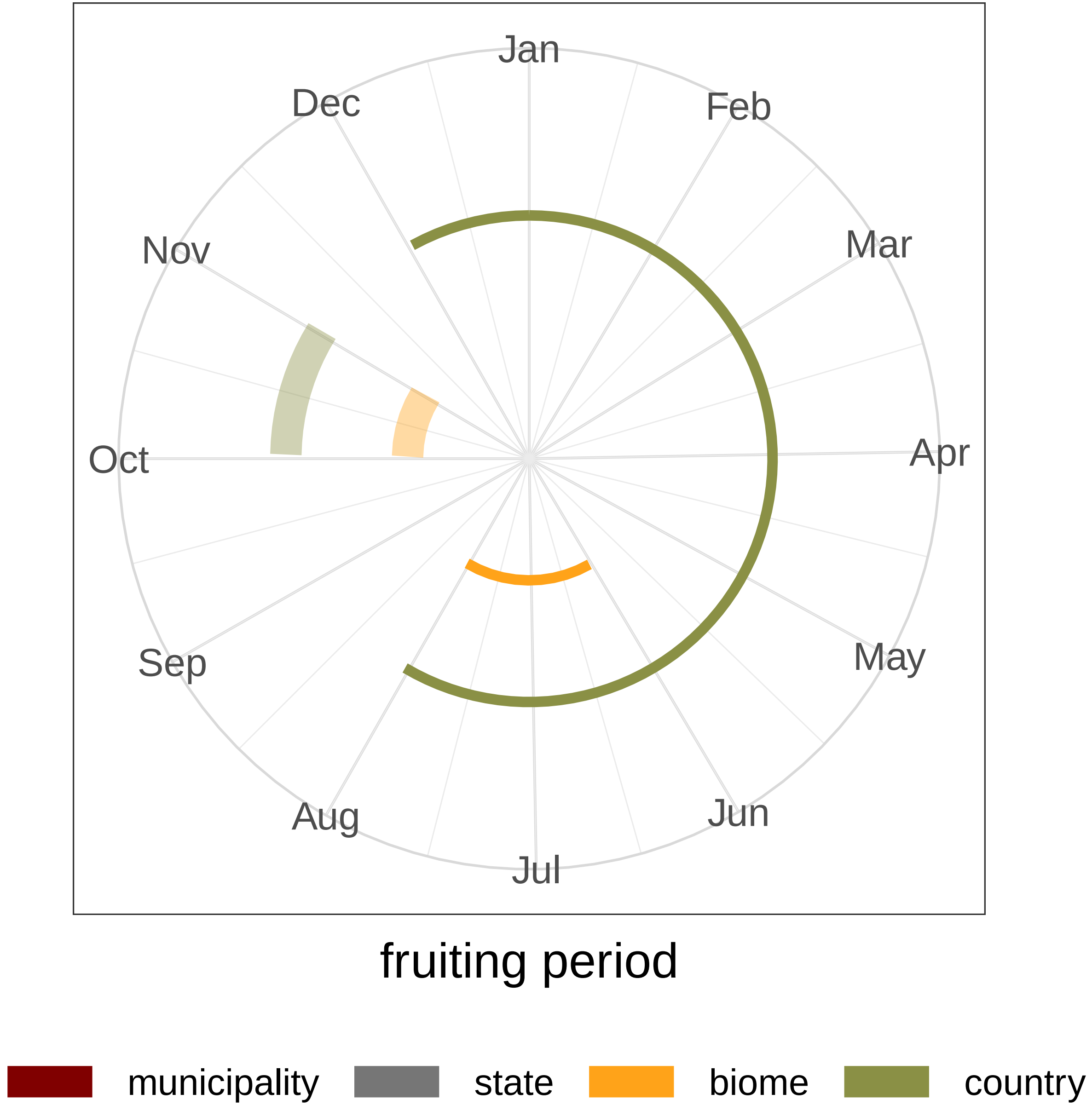

### *Banisteriopsis argyrophylla*

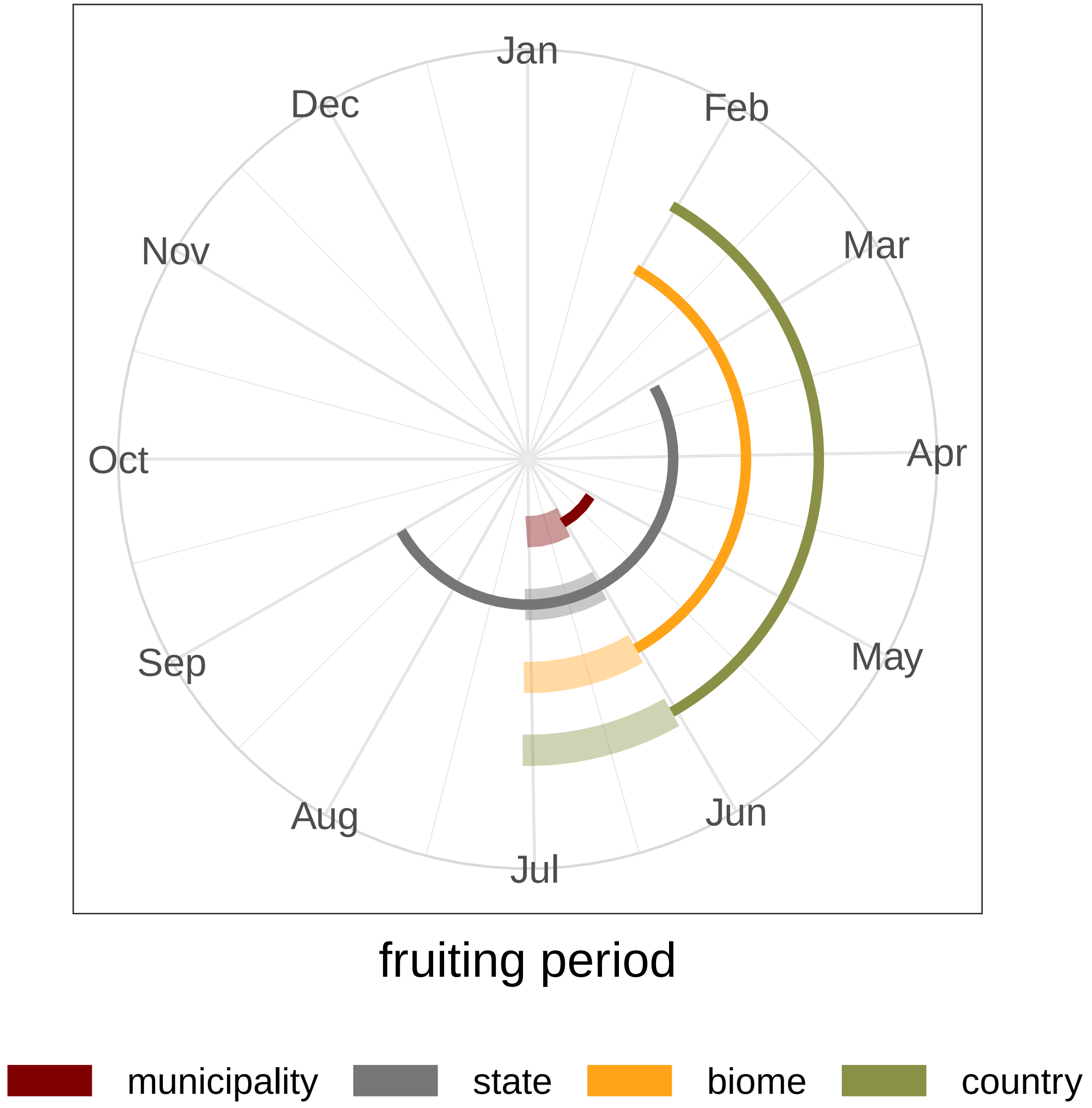

### *Banisteriopsis campestris*

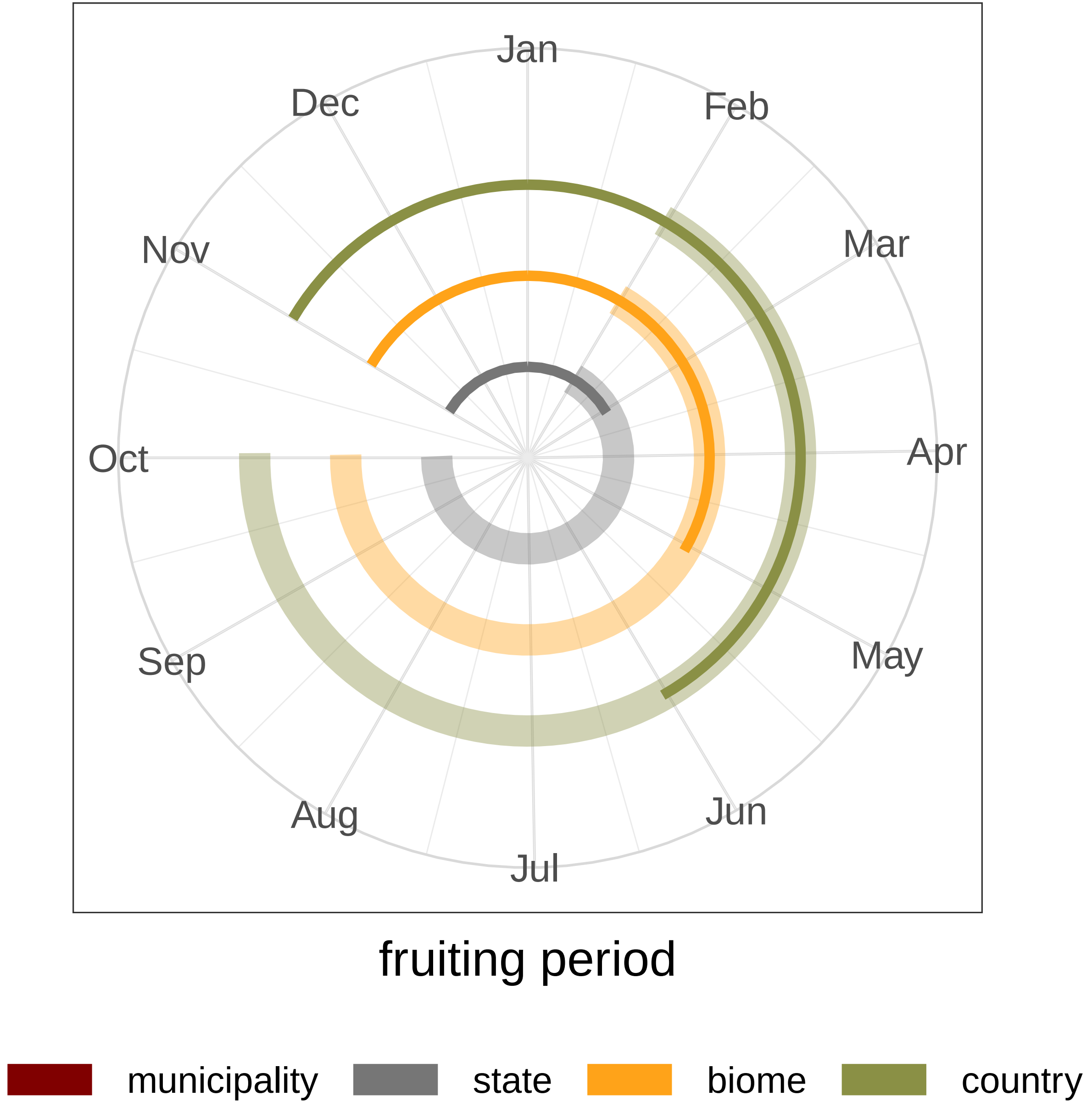

### *Banisteriopsis laevifolia*

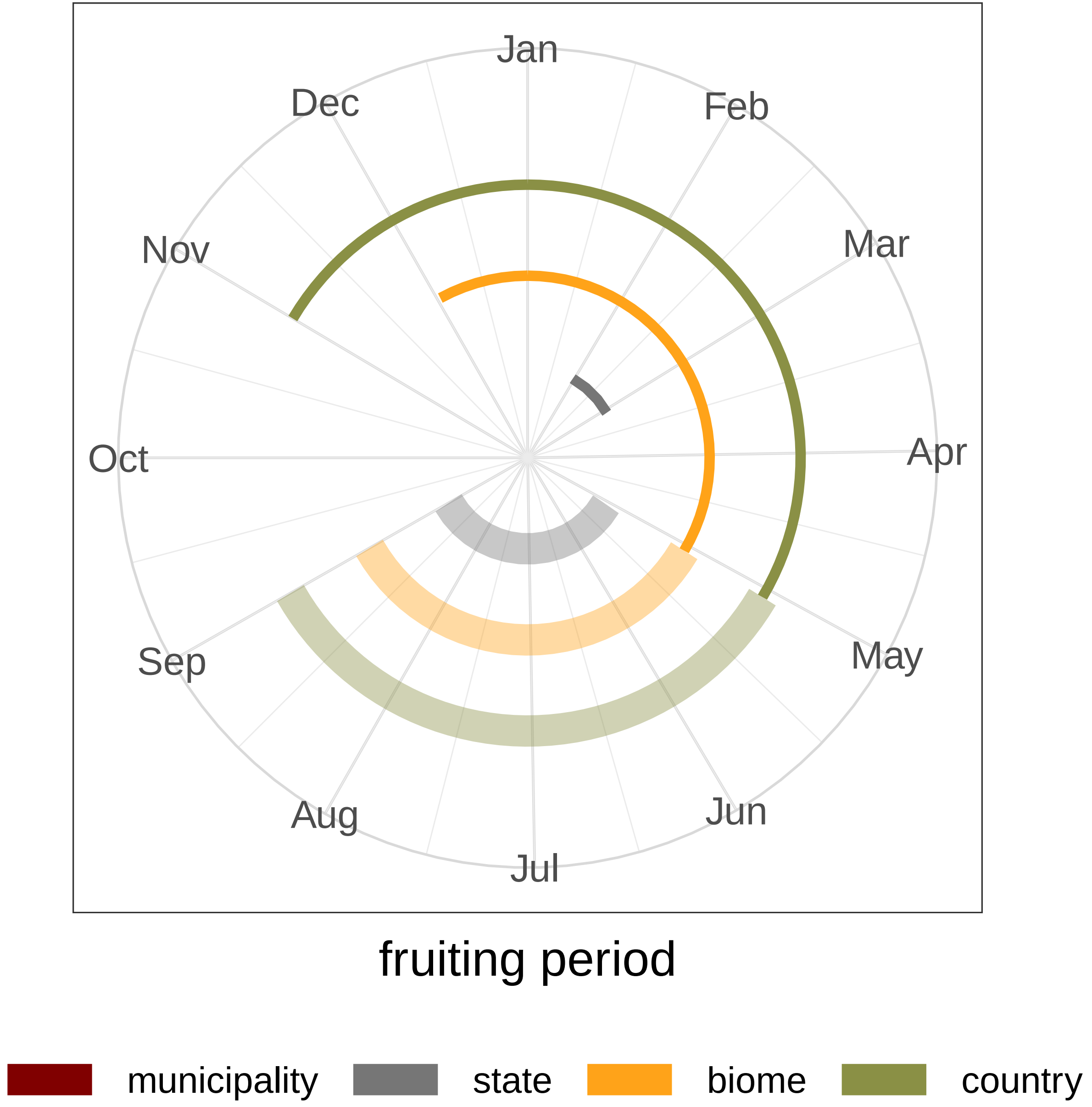

### *Banisteriopsis muricata*

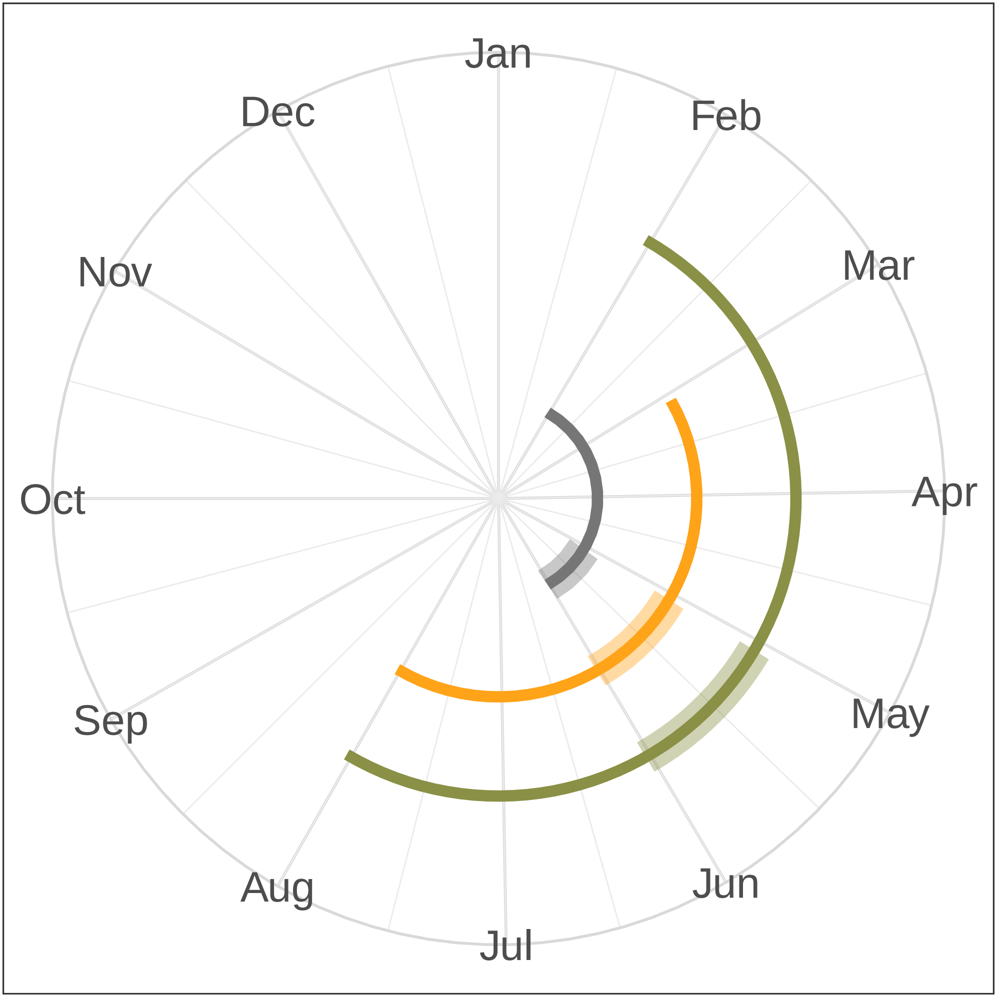


fruiting period

### *Banisteriopsis stellaris*

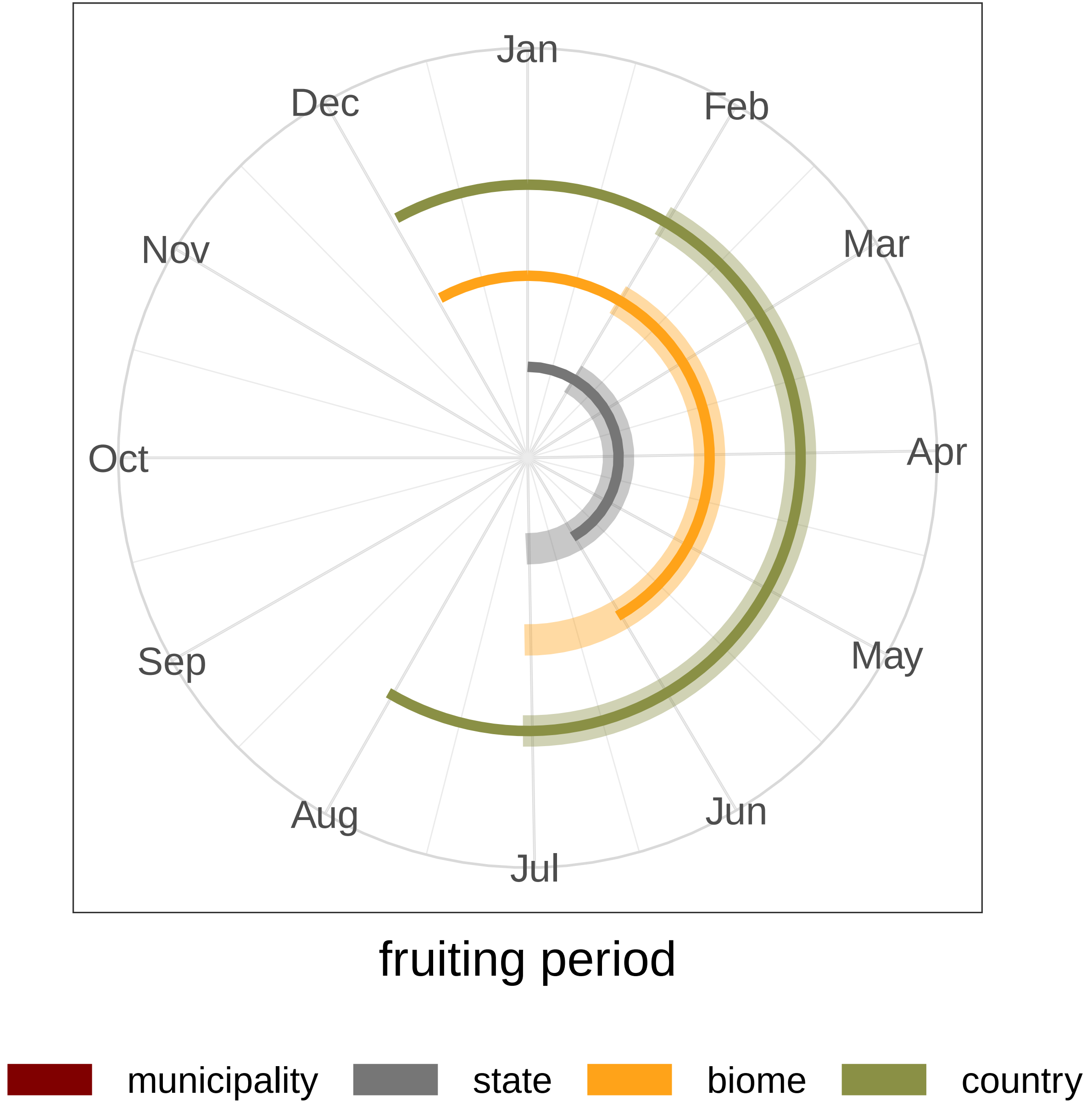

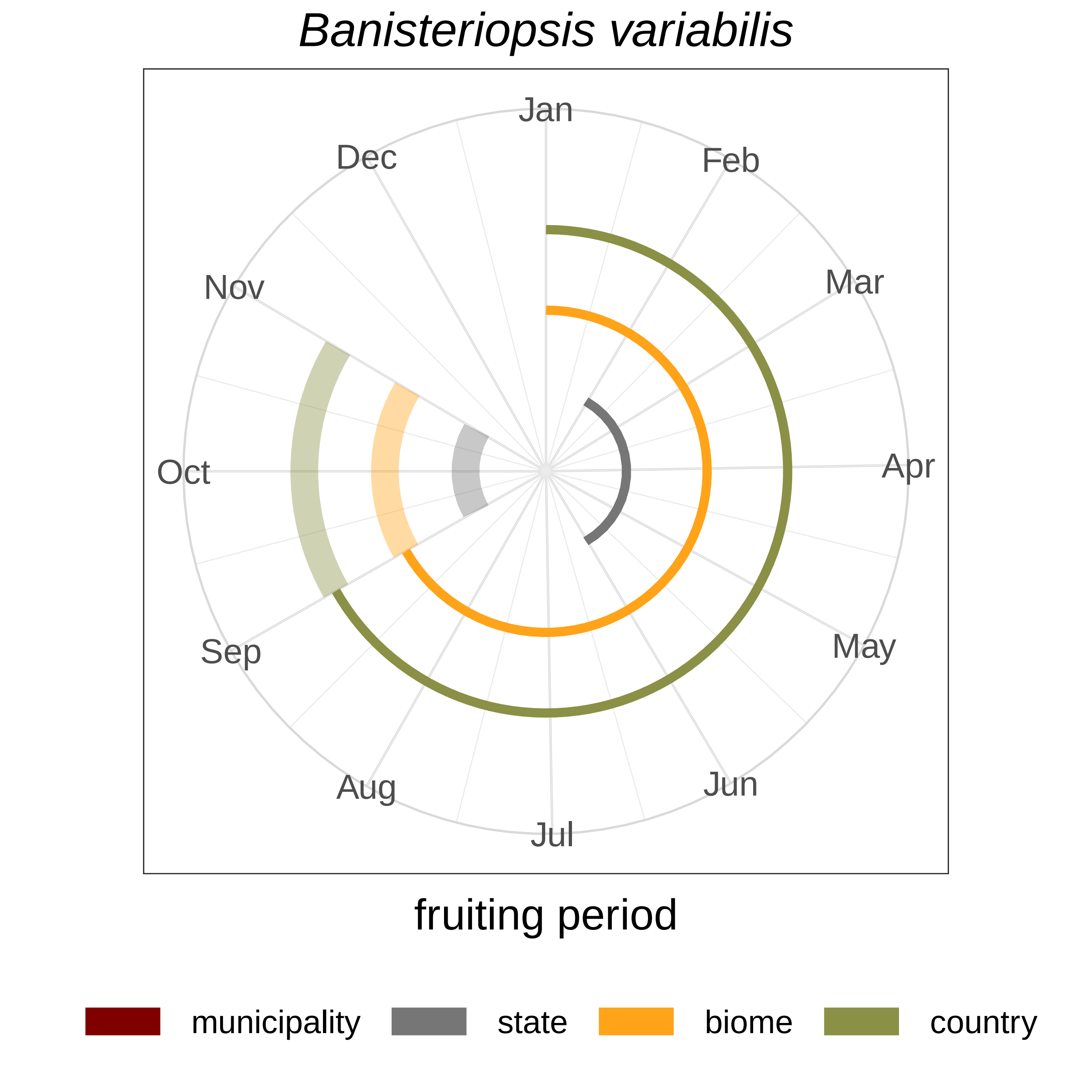
*Banisteriopsis variabilis*
Jan
Feb
Mar
Apr
May
Jun
Jul
Aug
Sep
Oct
Nov
Dec
fruiting period
municipality
state
biome
country

### *Chamaecrista desvauxii*

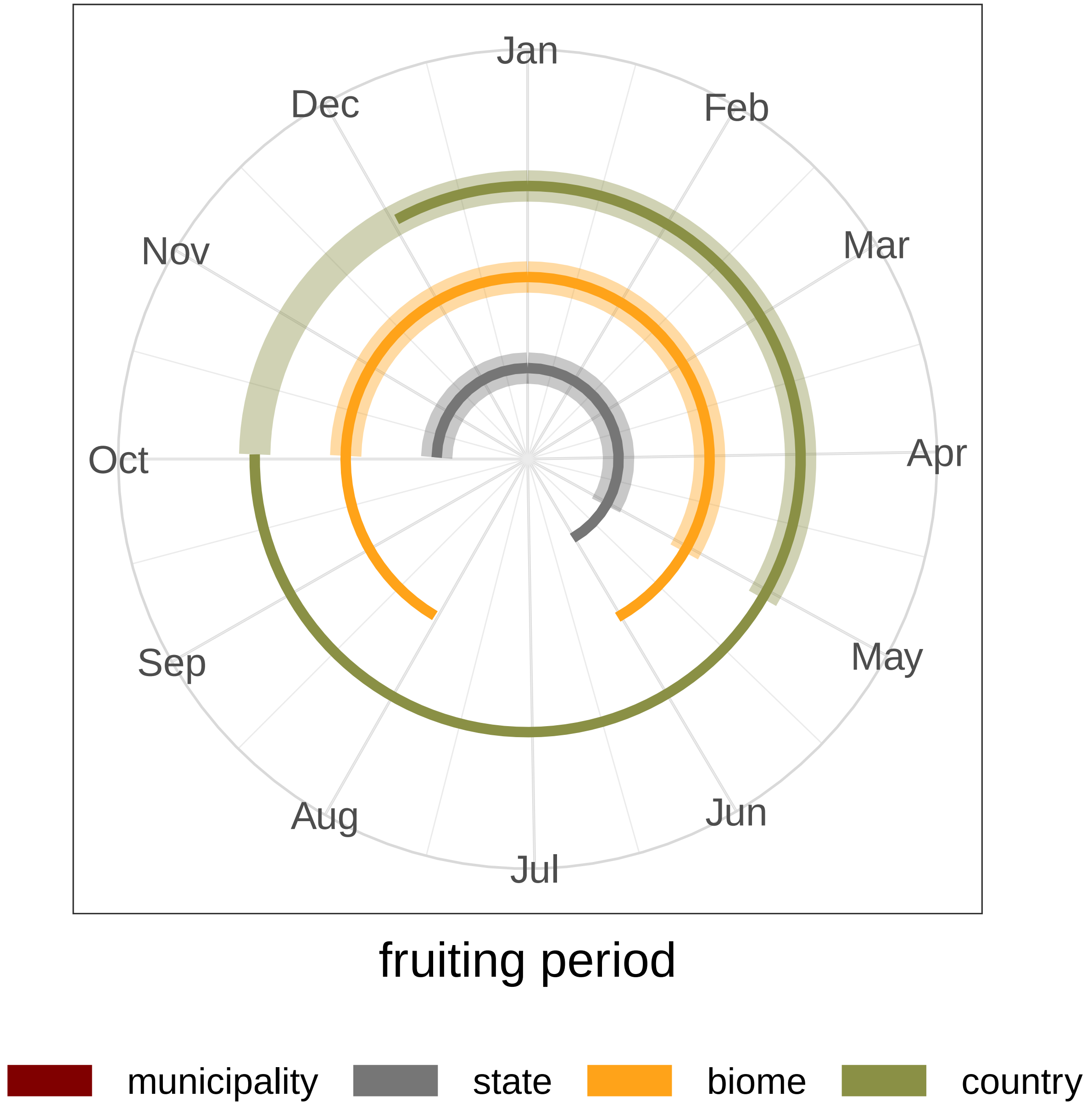

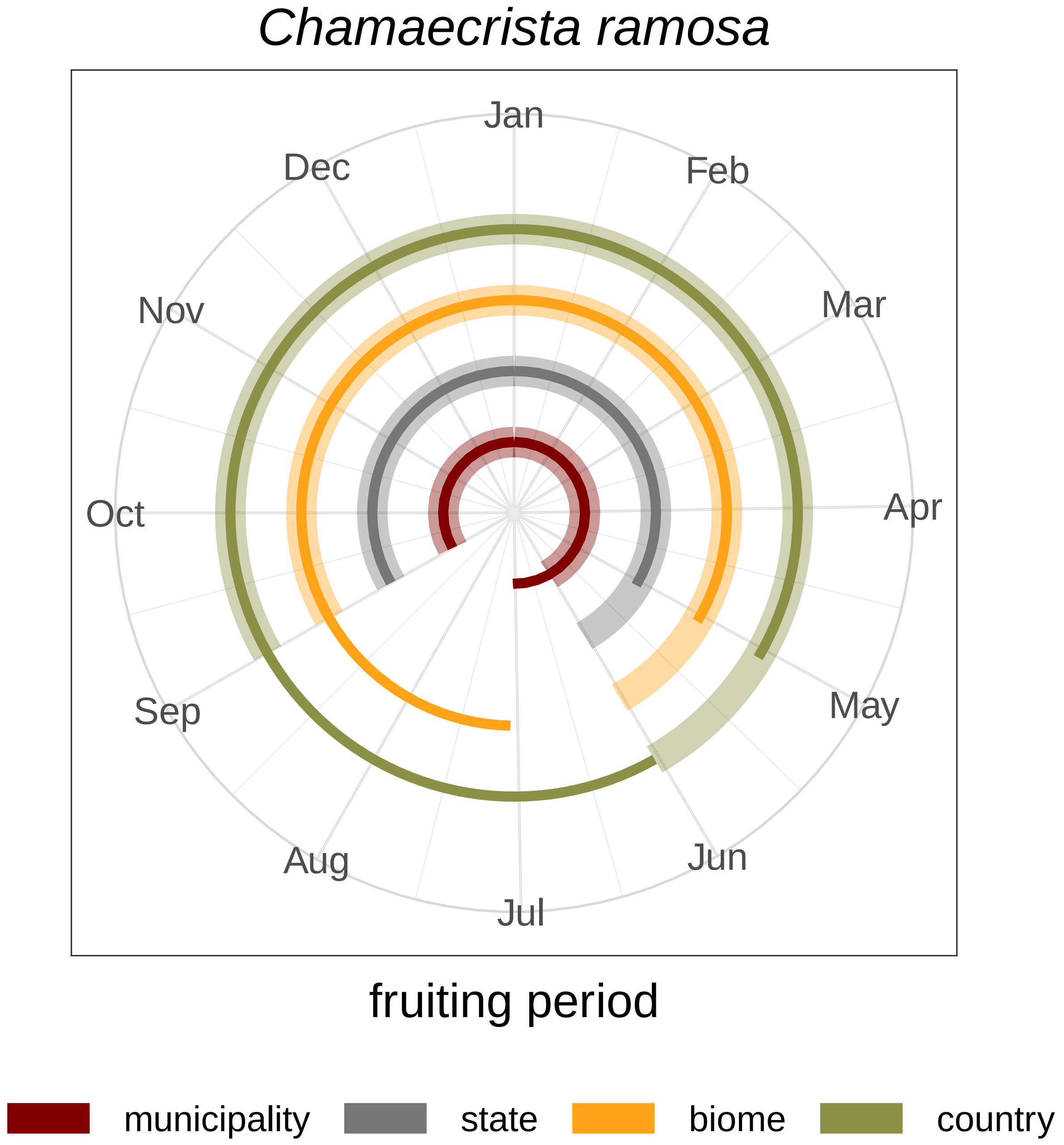
Chamaecrista ramosa
Jan
Feb
Mar
Apr
May
Jun
Jul
Aug
Sep
Oct
Nov
Dec
fruiting period
municipality
state
biome
country

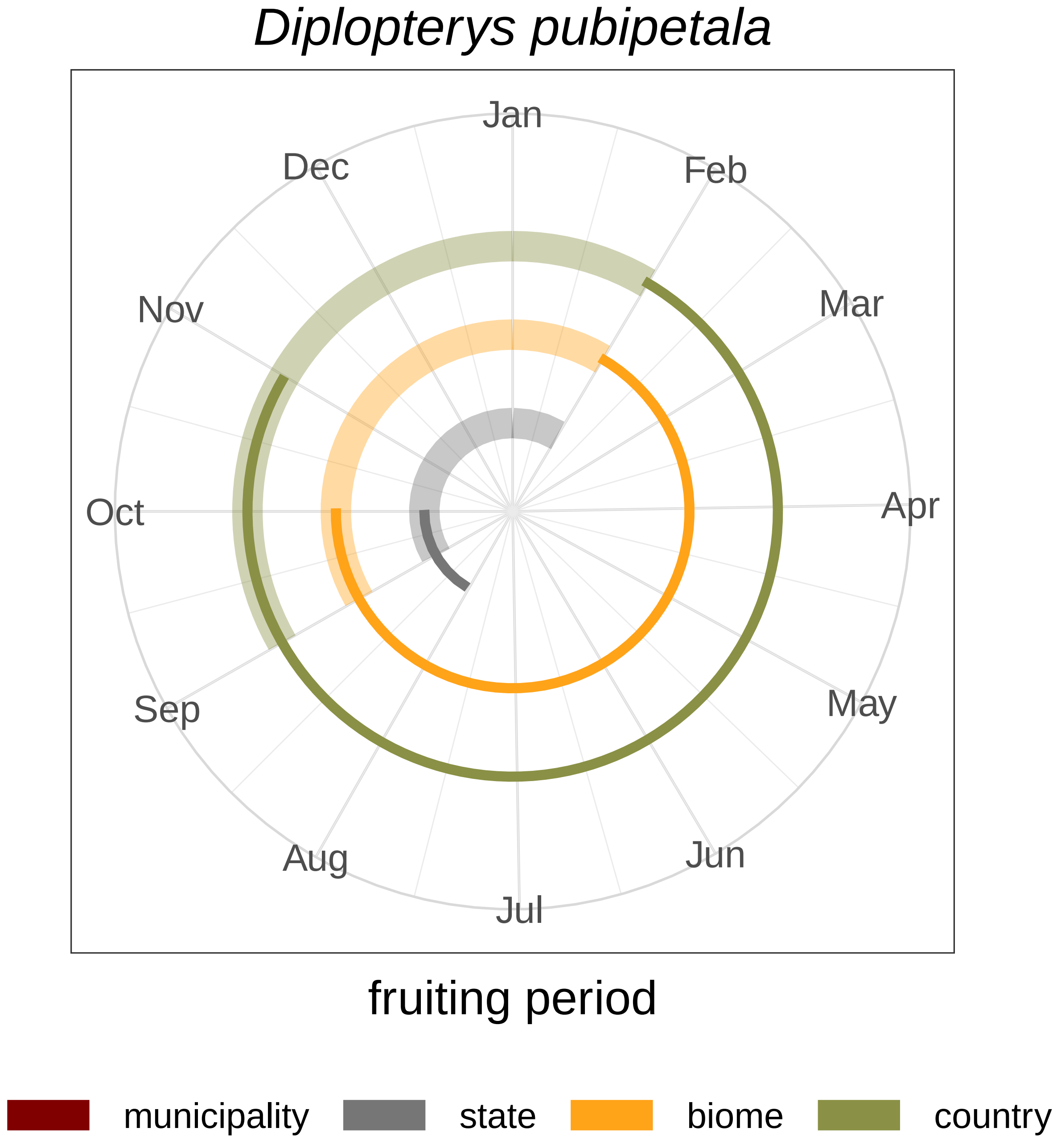

Diplopterys pubipetala
Jan
Feb
Mar
Apr
May
Jun
Jul
Aug
Sep
Oct
Nov
Dec
fruiting period
municipality
state
biome
country

### *Dolichandra unguis−cati*

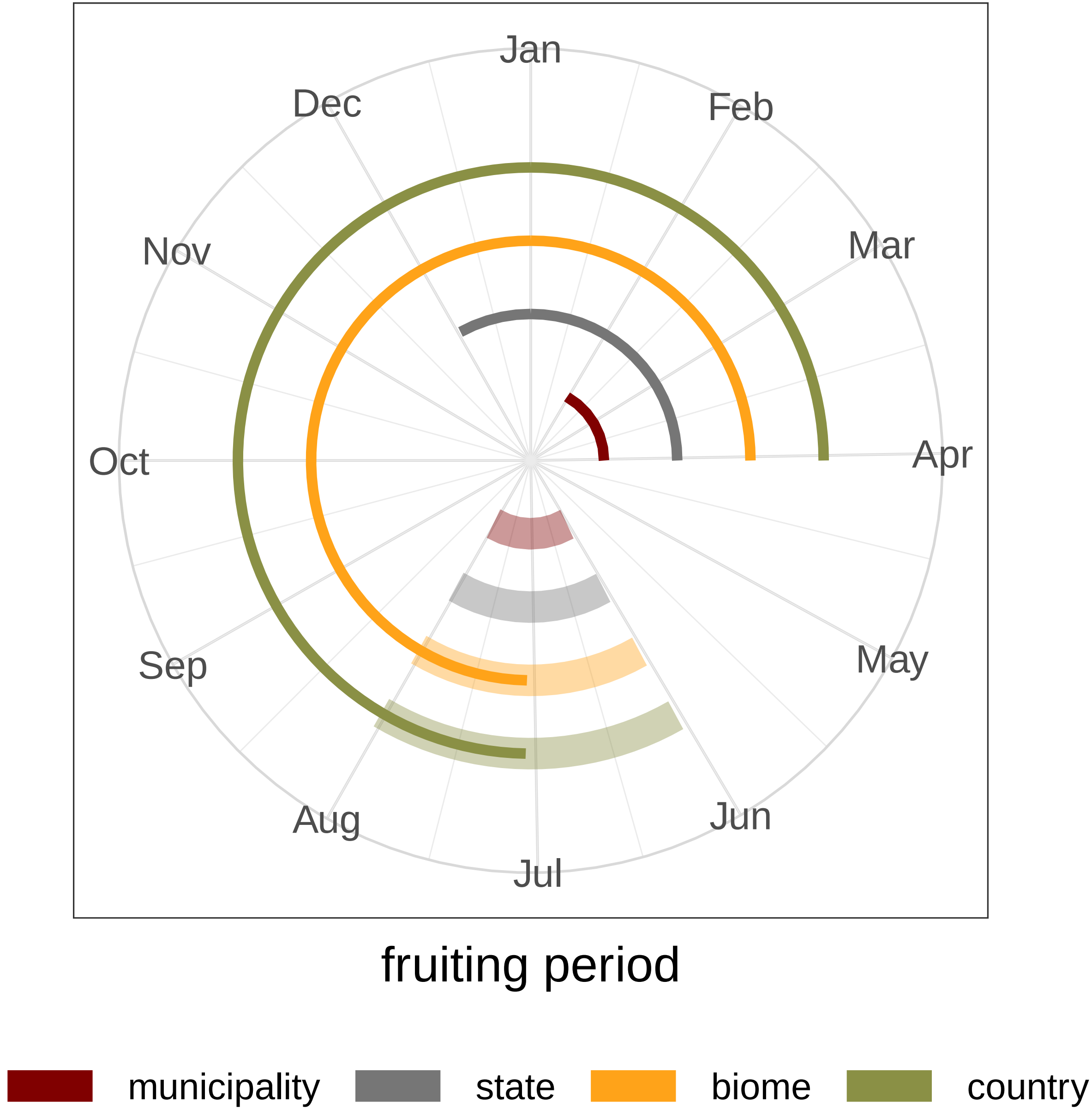

*Heteropterys byrsonimifolia*

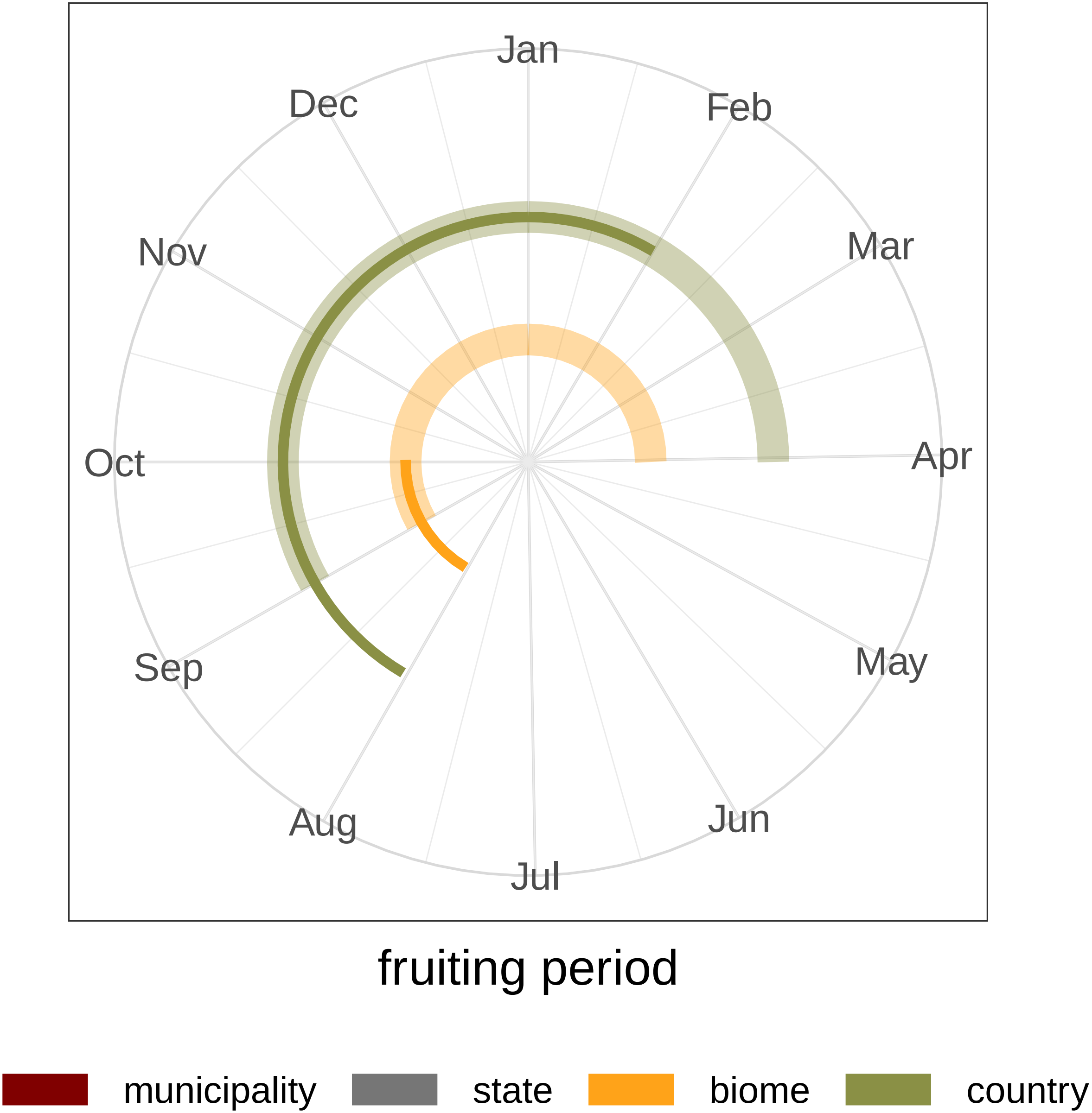


fruiting period

### *Heteropterys umbellata*

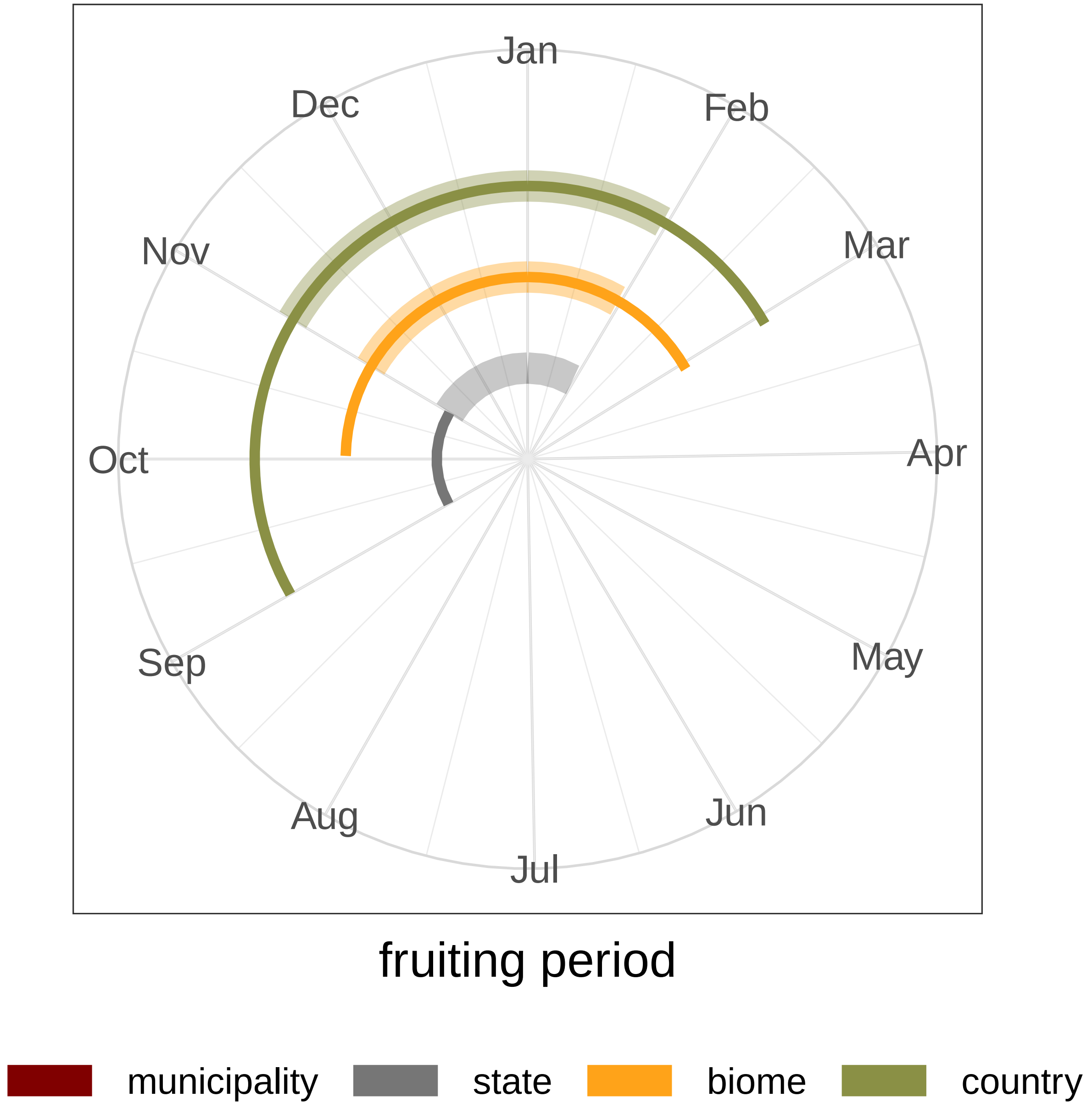

### *Jacaranda caroba*

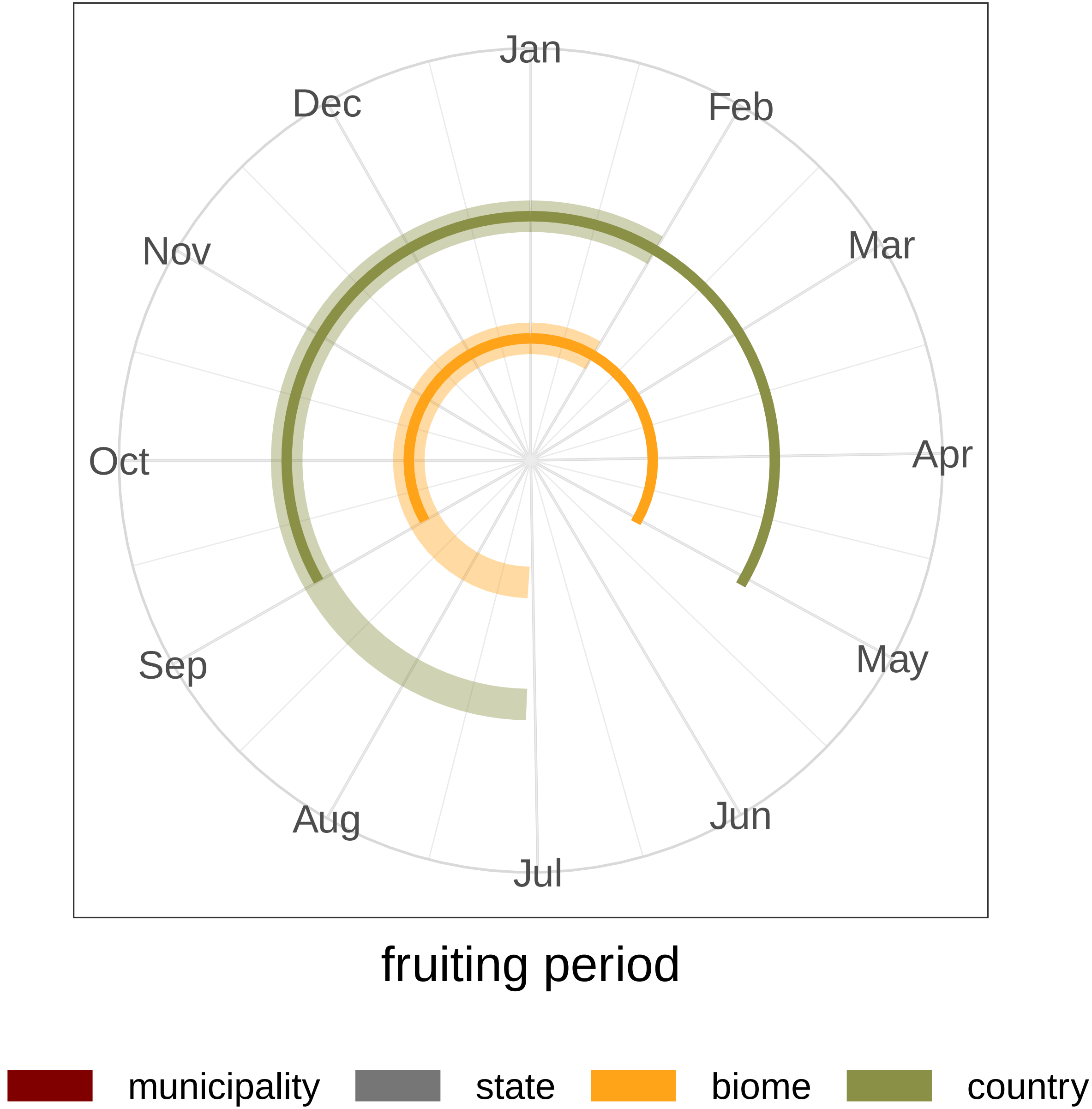

### *Licania heteromorpha*

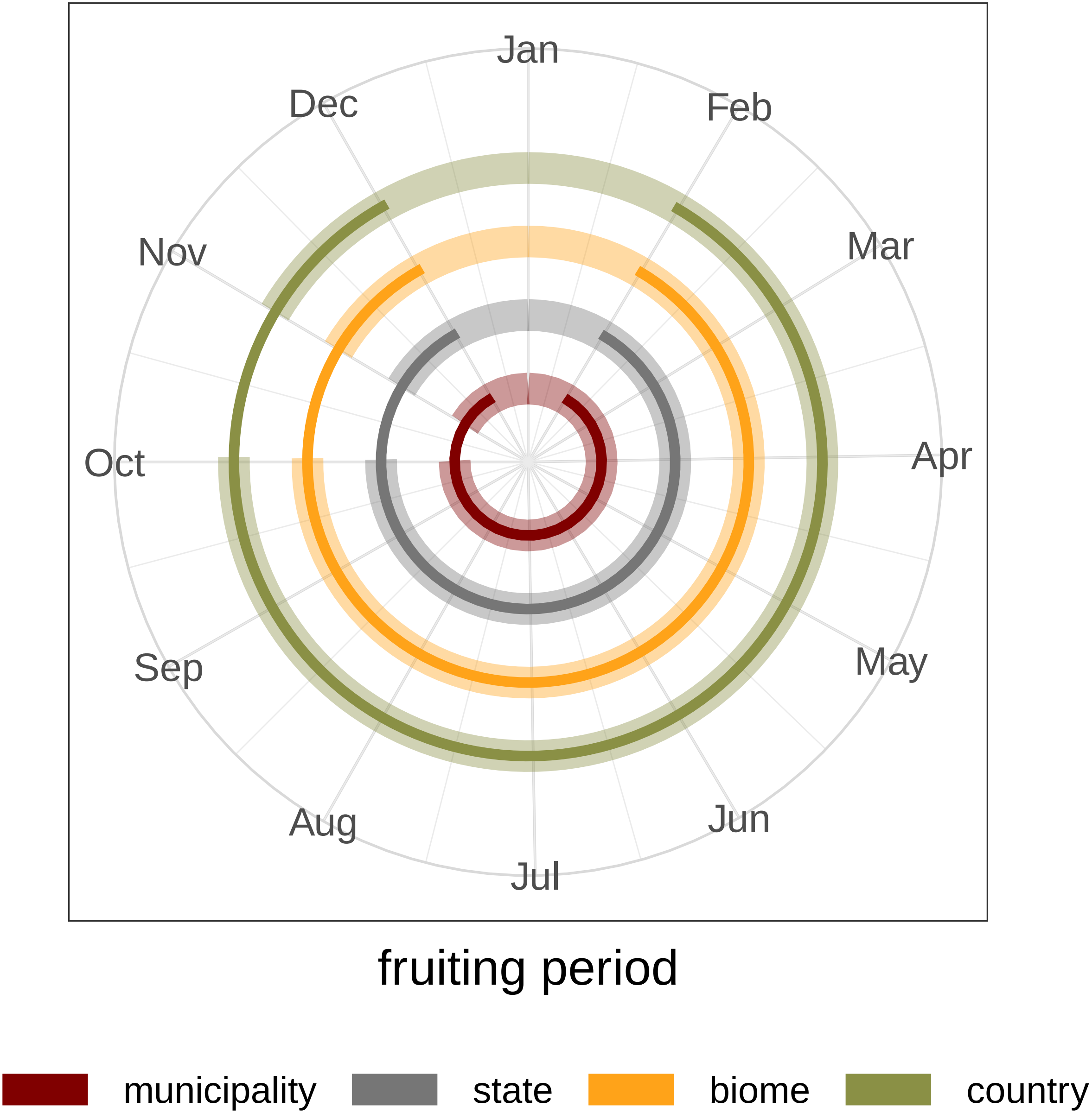

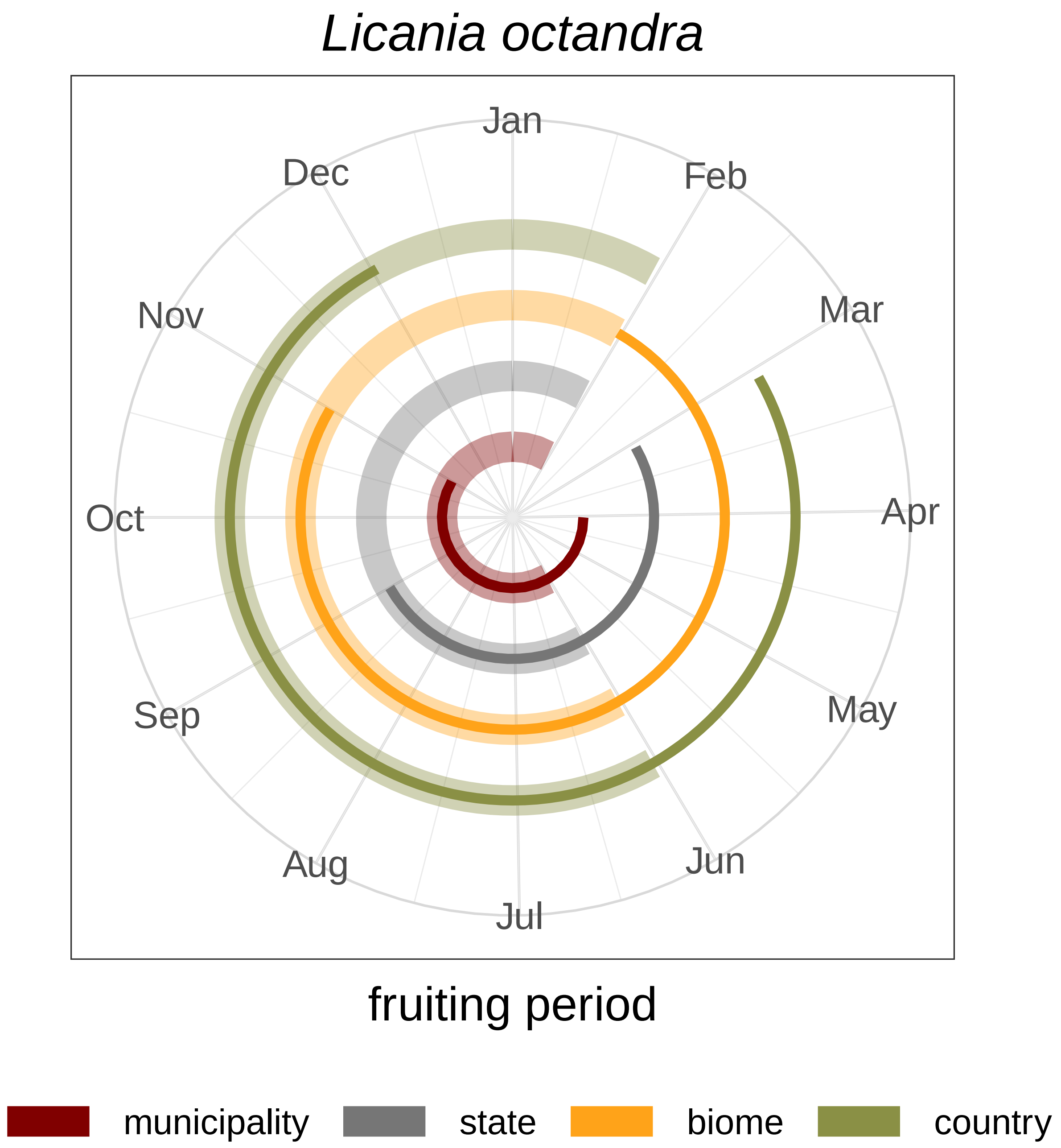
Licania octandra
Jan
Feb
Mar
Apr
May
Jun
Jul
Aug
Sep
Oct
Nov
Dec
fruiting period
municipality
state
biome
country

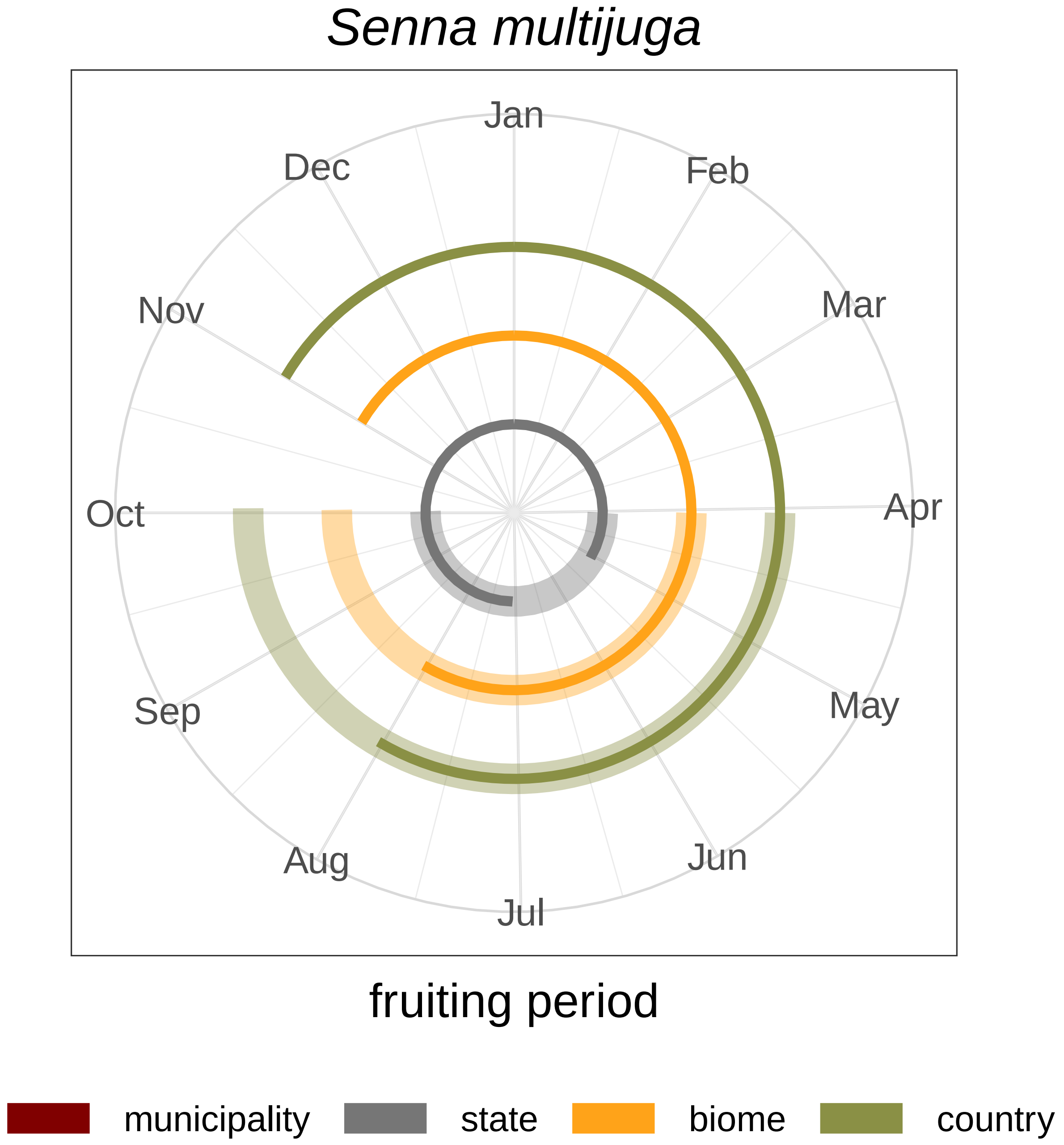
Senna multijuga
Jan
Feb
Mar
Apr
May
Jun
Jul
Aug
Sep
Oct
Nov
Dec
fruiting period
municipality
state
biome
country
