## Supplementary material for "Herbarium records provide reliable phenology estimates in the understudied tropics": Table S1

**Table S1**. Tropical plant phenological observations from the literature. Zeros indicate absences, ones presences, and blanks missing data.

| family | binomial_species | phenophase | Jan | Feb | Mar | April | May | Jun | July | Aug | Sept | Oct | Nov | Dec | Latitude | Longitude | state | Municipality | Biome | Year | study_duration (years) | Area_of_phenological_study | Reference |
|---|---|---|---|---|---|---|---|---|---|---|---|---|---|---|---|---|---|---|---|---|---|---|---|
| Bignoniaceae | Amphilophium_crucigerum | Flowering | | | | 0 | 0 | 0 | 0 | 0 | 0 | 1 | 1 | 1 | -25.41666667 | -49.31666667 | PR | Curitiba | Mata Atlantica | 1996 | 2 | Araucaria_forest_Southern_Brazil | Marques_et_al_2004_Plant_Ecology_173_203_213 |
| Bignoniaceae | Amphilophium_crucigerum | Fruiting | | | | 1 | 1 | 1 | 1 | 1 | 1 | 0 | 0 | 0 | -25.41666667 | -49.31666667 | PR | Curitiba | Mata Atlantica | 1996 | 2 | Araucaria_forest_Southern_Brazil | Marques_et_al_2004_Plant_Ecology_173_203_213 |
| Bignoniaceae | Amphilophium_crucigerum | Leaf_fall | | | | 1 | 1 | 1 | 1 | 1 | 1 | 0 | 0 | 0 | -25.41666667 | -49.31666667 | PR | Curitiba | Mata Atlantica | 1996 | 2 | Araucaria_forest_Southern_Brazil | Marques_et_al_2004_Plant_Ecology_173_203_213 |
| Bignoniaceae | Amphilophium_crucigerum | Flushing | | | | 0 | 0 | 0 | 0 | 0 | 1 | 1 | 1 | 1 | -25.41666667 | -49.31666667 | PR | Curitiba | Mata Atlantica | 1996 | 2 | Araucaria_forest_Southern_Brazil | Marques_et_al_2004_Plant_Ecology_173_203_213 |
| Bignoniaceae | Amphilophium_crucigerum | Flowering | 0 | 0 | 0 | 0 | 0 | 0 | 0 | 0 | 0 | 1 | 1 | 1 | -25.41666667 | -49.31666667 | PR | Curitiba | Mata Atlantica | 1997 | 2 | Araucaria_forest_Southern_Brazil | Marques_et_al_2004_Plant_Ecology_173_203_213 |
| Bignoniaceae | Amphilophium_crucigerum | Fruiting | 0 | 1 | 1 | 1 | 1 | 1 | 1 | 1 | 1 | 0 | 0 | 0 | -25.41666667 | -49.31666667 | PR | Curitiba | Mata Atlantica | 1997 | 2 | Araucaria_forest_Southern_Brazil | Marques_et_al_2004_Plant_Ecology_173_203_213 |
| Bignoniaceae | Amphilophium_crucigerum | Leaf_fall | 0 | 0 | 0 | 1 | 1 | 1 | 1 | 1 | 1 | 0 | 0 | 0 | -25.41666667 | -49.31666667 | PR | Curitiba | Mata Atlantica | 1997 | 2 | Araucaria_forest_Southern_Brazil | Marques_et_al_2004_Plant_Ecology_173_203_213 |
| Bignoniaceae | Amphilophium_crucigerum | Flushing | 1 | 1 | 1 | 0 | 0 | 0 | 0 | 0 | 1 | 1 | 1 | 1 | -25.41666667 | -49.31666667 | PR | Curitiba | Mata Atlantica | 1997 | 2 | Araucaria_forest_Southern_Brazil | Marques_et_al_2004_Plant_Ecology_173_203_213 |
| Bignoniaceae | Amphilophium_crucigerum | Flowering | 0 | 0 | 0 | | | | | | | | | | -25.41666667 | -49.31666667 | PR | Curitiba | Mata Atlantica | 1998 | 2 | Araucaria_forest_Southern_Brazil | Marques_et_al_2004_Plant_Ecology_173_203_213 |
| Bignoniaceae | Amphilophium_crucigerum | Fruiting | 0 | 1 | 1 | | | | | | | | | | -25.41666667 | -49.31666667 | PR | Curitiba | Mata Atlantica | 1998 | 2 | Araucaria_forest_Southern_Brazil | Marques_et_al_2004_Plant_Ecology_173_203_213 |
| Bignoniaceae | Amphilophium_crucigerum | Leaf_fall | 0 | 0 | 0 | | | | | | | | | | -25.41666667 | -49.31666667 | PR | Curitiba | Mata Atlantica | 1998 | 2 | Araucaria_forest_Southern_Brazil | Marques_et_al_2004_Plant_Ecology_173_203_213 |
| Bignoniaceae | Amphilophium_crucigerum | Flushing | 1 | 1 | 1 | | | | | | | | | | -25.41666667 | -49.31666667 | PR | Curitiba | Mata Atlantica | 1998 | 2 | Araucaria_forest_Southern_Brazil | Marques_et_al_2004_Plant_Ecology_173_203_213 |
| Bignoniaceae | Amphilophium_crucigerum | Flowering | | | 0 | 0 | 0 | 0 | 0 | 0 | 0 | 0 | 0 | 1 | -22.82916667 | -47.10916667 | SP | Campinas | Mata Atlantica | 1988 | 3 | Semideciduous_forest_Southeastern_Brazil | Morellato_e_Leitão_Filho_1996_Biotropica_282_180_19 |
| Bignoniaceae | Amphilophium_crucigerum | Fruiting | | | 0 | 0 | 0 | 0 | 1 | 1 | 1 | 0 | 0 | 0 | -22.82916667 | -47.10916667 | SP | Campinas | Mata Atlantica | 1988 | 3 | Semideciduous_forest_Southeastern_Brazil | Morellato_e_Leitão_Filho_1996_Biotropica_282_180_19 |
| Bignoniaceae | Amphilophium_crucigerum | Leaf_fall | | | 0 | 0 | 0 | 0 | 0 | 0 | 0 | 0 | 0 | 0 | -22.82916667 | -47.10916667 | SP | Campinas | Mata Atlantica | 1988 | 3 | Semideciduous_forest_Southeastern_Brazil | Morellato_e_Leitão_Filho_1996_Biotropica_282_180_19 |
| Bignoniaceae | Amphilophium_crucigerum | Flushing | | | 0 | 0 | 0 | 0 | 0 | 0 | 0 | 0 | 0 | 0 | -22.82916667 | -47.10916667 | SP | Campinas | Mata Atlantica | 1988 | 3 | Semideciduous_forest_Southeastern_Brazil | Morellato_e_Leitão_Filho_1996_Biotropica_282_180_19 |
| Bignoniaceae | Amphilophium_crucigerum | Flowering | 1 | 1 | 0 | 0 | 0 | 0 | 0 | 0 | 0 | 0 | 0 | 1 | -22.82916667 | -47.10916667 | SP | Campinas | Mata Atlantica | 1989 | 3 | Semideciduous_forest_Southeastern_Brazil | Morellato_e_Leitão_Filho_1996_Biotropica_282_180_19 |
| Bignoniaceae | Amphilophium_crucigerum | Fruiting | 0 | 0 | 0 | 0 | 0 | 0 | 1 | 1 | 1 | 0 | 0 | 0 | -22.82916667 | -47.10916667 | SP | Campinas | Mata Atlantica | 1989 | 3 | Semideciduous_forest_Southeastern_Brazil | Morellato_e_Leitão_Filho_1996_Biotropica_282_180_19 |
| Bignoniaceae | Amphilophium_crucigerum | Leaf_fall | | | | | | | | | | | | | -22.82916667 | -47.10916667 | SP | Campinas | Mata Atlantica | 1989 | 3 | Semideciduous_forest_Southeastern_Brazil | Morellato_e_Leitão_Filho_1996_Biotropica_282_180_19 |
| Bignoniaceae | Amphilophium_crucigerum | Flushing | | | | | | | | | | | | | -22.82916667 | -47.10916667 | SP | Campinas | Mata Atlantica | 1989 | 3 | Semideciduous_forest_Southeastern_Brazil | Morellato_e_Leitão_Filho_1996_Biotropica_282_180_19 |
| Bignoniaceae | Amphilophium_crucigerum | Flowering | 1 | 1 | 0 | 0 | 0 | 0 | 0 | 0 | 0 | 0 | 0 | 1 | -22.82916667 | -47.10916667 | SP | Campinas | Mata Atlantica | 1990 | 3 | Semideciduous_forest_Southeastern_Brazil | Morellato_e_Leitão_Filho_1996_Biotropica_282_180_19 |
| Bignoniaceae | Amphilophium_crucigerum | Fruiting | 0 | 0 | 0 | 0 | 0 | 0 | 1 | 1 | 1 | 0 | 0 | 0 | -22.82916667 | -47.10916667 | SP | Campinas | Mata Atlantica | 1990 | 3 | Semideciduous_forest_Southeastern_Brazil | Morellato_e_Leitão_Filho_1996_Biotropica_282_180_19 |
| Bignoniaceae | Amphilophium_crucigerum | Leaf_fall | | | | | | | | | | | | | -22.82916667 | -47.10916667 | SP | Campinas | Mata Atlantica | 1990 | 3 | Semideciduous_forest_Southeastern_Brazil | Morellato_e_Leitão_Filho_1996_Biotropica_282_180_19 |
| Bignoniaceae | Amphilophium_crucigerum | Flushing | | | | | | | | | | | | | -22.82916667 | -47.10916667 | SP | Campinas | Mata Atlantica | 1990 | 3 | Semideciduous_forest_Southeastern_Brazil | Morellato_e_Leitão_Filho_1996_Biotropica_282_180_19 |
| Bignoniaceae | Amphilophium_crucigerum | Flowering | 1 | 1 | | | | | | | | | | | -22.82916667 | -47.10916667 | SP | Campinas | Mata Atlantica | 1991 | 3 | Semideciduous_forest_Southeastern_Brazil | Morellato_e_Leitão_Filho_1996_Biotropica_282_180_19 |
| Bignoniaceae | Amphilophium_crucigerum | Fruiting | | | | | | | | | | | | | -22.82916667 | -47.10916667 | SP | Campinas | Mata Atlantica | 1991 | 3 | Semideciduous_forest_Southeastern_Brazil | Morellato_e_Leitão_Filho_1996_Biotropica_282_180_19 |
| Bignoniaceae | Amphilophium_crucigerum | Leaf_fall | | | | | | | | | | | | | -22.82916667 | -47.10916667 | SP | Campinas | Mata Atlantica | 1991 | 3 | Semideciduous_forest_Southeastern_Brazil | Morellato_e_Leitão_Filho_1996_Biotropica_282_180_19 |
| Bignoniaceae | Amphilophium_crucigerum | Flushing | | | | | | | | | | | | | -22.82916667 | -47.10916667 | SP | Campinas | Mata Atlantica | 1991 | 3 | Semideciduous_forest_Southeastern_Brazil | Morellato_e_Leitão_Filho_1996_Biotropica_282_180_19 |
| Bignoniaceae | Anemopaegma_chamberlaynii | Flowering | | | | | | | | | 0 | 0 | 0 | 0 | -21.61083333 | -47.61055556 | SP | Santa Rita do Passa Quatro | Cerrado | 1995 | 2 | Cerrado_Pé-de-Gigante_Reserve_SP_BRAZIL | Batalha_e_Mantovani_2000_Rev_Brasil_Biol_601_129_145 |
| Bignoniaceae | Anemopaegma_chamberlaynii | Fruiting | | | | | | | | | 0 | 1 | 0 | 0 | -21.61083333 | -47.61055556 | SP | Santa Rita do Passa Quatro | Cerrado | 1995 | 2 | Cerrado_Pé-de-Gigante_Reserve_SP_BRAZIL | Batalha_e_Mantovani_2000_Rev_Brasil_Biol_601_129_145 |
| Bignoniaceae | Anemopaegma_chamberlaynii | Leaf_fall | | | | | | | | | | | | | -21.61083333 | -47.61055556 | SP | Santa Rita do Passa Quatro | Cerrado | 1995 | 2 | Cerrado_Pé-de-Gigante_Reserve_SP_BRAZIL | Batalha_e_Mantovani_2000_Rev_Brasil_Biol_601_129_145 |
| Bignoniaceae | Anemopaegma_chamberlaynii | Flushing | | | | | | | | | | | | | -21.61083333 | -47.61055556 | SP | Santa Rita do Passa Quatro | Cerrado | 1995 | 2 | Cerrado_Pé-de-Gigante_Reserve_SP_BRAZIL | Batalha_e_Mantovani_2000_Rev_Brasil_Biol_601_129_145 |
| Bignoniaceae | Anemopaegma_chamberlaynii | Flowering | 0 | 0 | 0 | 0 | 0 | 0 | 0 | 0 | 0 | 0 | 0 | 0 | -21.61083333 | -47.61055556 | SP | Santa Rita do Passa Quatro | Cerrado | 1996 | 2 | Cerrado_Pé-de-Gigante_Reserve_SP_BRAZIL | Batalha_e_Mantovani_2000_Rev_Brasil_Biol_601_129_145 |
| Bignoniaceae | Anemopaegma_chamberlaynii | Fruiting | 0 | 0 | 0 | 0 | 0 | 0 | 0 | 0 | 0 | 1 | 0 | 0 | -21.61083333 | -47.61055556 | SP | Santa Rita do Passa Quatro | Cerrado | 1996 | 2 | Cerrado_Pé-de-Gigante_Reserve_SP_BRAZIL | Batalha_e_Mantovani_2000_Rev_Brasil_Biol_601_129_145 |
| Bignoniaceae | Anemopaegma_chamberlaynii | Leaf_fall | | | | | | | | | | | | | -21.61083333 | -47.61055556 | SP | Santa Rita do Passa Quatro | Cerrado | 1996 | 2 | Cerrado_Pé-de-Gigante_Reserve_SP_BRAZIL | Batalha_e_Mantovani_2000_Rev_Brasil_Biol_601_129_145 |
| Bignoniaceae | Anemopaegma_chamberlaynii | Flushing | | | | | | | | | | | | | -21.61083333 | -47.61055556 | SP | Santa Rita do Passa Quatro | Cerrado | 1996 | 2 | Cerrado_Pé-de-Gigante_Reserve_SP_BRAZIL | Batalha_e_Mantovani_2000_Rev_Brasil_Biol_601_129_145 |
| Bignoniaceae | Anemopaegma_chamberlaynii | Flowering | 0 | 0 | | | | | | | | | | | -21.61083333 | -47.61055556 | SP | Santa Rita do Passa Quatro | Cerrado | 1997 | 2 | Cerrado_Pé-de-Gigante_Reserve_SP_BRAZIL | Batalha_e_Mantovani_2000_Rev_Brasil_Biol_601_129_145 |
| Bignoniaceae | Anemopaegma_chamberlaynii | Fruiting | 0 | 0 | | | | | | | | | | | -21.61083333 | -47.61055556 | SP | Santa Rita do Passa Quatro | Cerrado | 1997 | 2 | Cerrado_Pé-de-Gigante_Reserve_SP_BRAZIL | Batalha_e_Mantovani_2000_Rev_Brasil_Biol_601_129_145 |
| Bignoniaceae | Anemopaegma_chamberlaynii | Leaf_fall | | | | | | | | | | | | | -21.61083333 | -47.61055556 | SP | Santa Rita do Passa Quatro | Cerrado | 1997 | 2 | Cerrado_Pé-de-Gigante_Reserve_SP_BRAZIL | Batalha_e_Mantovani_2000_Rev_Brasil_Biol_601_129_145 |
| Bignoniaceae | Anemopaegma_chamberlaynii | Flushing | | | | | | | | | | | | | -21.61083333 | -47.61055556 | SP | Santa Rita do Passa Quatro | Cerrado | 1997 | 2 | Cerrado_Pé-de-Gigante_Reserve_SP_BRAZIL | Batalha_e_Mantovani_2000_Rev_Brasil_Biol_601_129_145 |
| Bignoniaceae | Dolichandra_unguis_cati | Flowering | | | 0 | 0 | 0 | 0 | 0 | 0 | 1 | 1 | 1 | 0 | -22.82916667 | -47.10916667 | SP | Campinas | Mata Atlantica | 1988 | 3 | Semideciduous_forest_Southeastern_Brazil | Morellato_e_Leitão_Filho_1996_Biotropica_282_180_19 |
| Bignoniaceae | Dolichandra_unguis_cati | Fruiting | | | 0 | 0 | 0 | 1 | 1 | 1 | 0 | 0 | 0 | 0 | -22.82916667 | -47.10916667 | SP | Campinas | Mata Atlantica | 1988 | 3 | Semideciduous_forest_Southeastern_Brazil | Morellato_e_Leitão_Filho_1996_Biotropica_282_180_19 |
| Bignoniaceae | Dolichandra_unguis_cati | Leaf_fall | | | | | | | | | | | | | -22.82916667 | -47.10916667 | SP | Campinas | Mata Atlantica | 1988 | 3 | Semideciduous_forest_Southeastern_Brazil | Morellato_e_Leitão_Filho_1996_Biotropica_282_180_19 |
| Bignoniaceae | Dolichandra_unguis_cati | Flushing | | | | | | | | | | | | | -22.82916667 | -47.10916667 | SP | Campinas | Mata Atlantica | 1988 | 3 | Semideciduous_forest_Southeastern_Brazil | Morellato_e_Leitão_Filho_1996_Biotropica_282_180_19 |
| Bignoniaceae | Dolichandra_unguis_cati | Flowering | 0 | 0 | 0 | 0 | 0 | 0 | 0 | 0 | 1 | 1 | 1 | 0 | -22.82916667 | -47.10916667 | SP | Campinas | Mata Atlantica | 1989 | 3 | Semideciduous_forest_Southeastern_Brazil | Morellato_e_Leitão_Filho_1996_Biotropica_282_180_19 |
| Bignoniaceae | Dolichandra_unguis_cati | Fruiting | 0 | 0 | 0 | 0 | 0 | 1 | 1 | 1 | 0 | 0 | 0 | 0 | -22.82916667 | -47.10916667 | SP | Campinas | Mata Atlantica | 1989 | 3 | Semideciduous_forest_Southeastern_Brazil | Morellato_e_Leitão_Filho_1996_Biotropica_282_180_19 |
| Bignoniaceae | Dolichandra_unguis_cati | Leaf_fall | | | | | | | | | | | | | -22.82916667 | -47.10916667 | SP | Campinas | Mata Atlantica | 1989 | 3 | Semideciduous_forest_Southeastern_Brazil | Morellato_e_Leitão_Filho_1996_Biotropica_282_180_19 |
| Bignoniaceae | Dolichandra_unguis_cati | Flushing | | | | | | | | | | | | | -22.82916667 | -47.10916667 | SP | Campinas | Mata Atlantica | 1989 | 3 | Semideciduous_forest_Southeastern_Brazil | Morellato_e_Leitão_Filho_1996_Biotropica_282_180_19 |
| Bignoniaceae | Dolichandra_unguis_cati | Flowering | 0 | 0 | 0 | 0 | 0 | 0 | 0 | 0 | 1 | 1 | 1 | 0 | -22.82916667 | -47.10916667 | SP | Campinas | Mata Atlantica | 1990 | 3 | Semideciduous_forest_Southeastern_Brazil | Morellato_e_Leitão_Filho_1996_Biotropica_282_180_19 |
| Bignoniaceae | Dolichandra_unguis_cati | Fruiting | 0 | 0 | 0 | 0 | 0 | 1 | 1 | 1 | 0 | 0 | 0 | 0 | -22.82916667 | -47.10916667 | SP | Campinas | Mata Atlantica | 1990 | 3 | Semideciduous_forest_Southeastern_Brazil | Morellato_e_Leitão_Filho_1996_Biotropica_282_180_19 |
| Bignoniaceae | Dolichandra_unguis_cati | Leaf_fall | | | | | | | | | | | | | -22.82916667 | -47.10916667 | SP | Campinas | Mata Atlantica | 1990 | 3 | Semideciduous_forest_Southeastern_Brazil | Morellato_e_Leitão_Filho_1996_Biotropica_282_180_19 |
| Bignoniaceae | Dolichandra_unguis_cati | Flushing | | | | | | | | | | | | | -22.82916667 | -47.10916667 | SP | Campinas | Mata Atlantica | 1990 | 3 | Semideciduous_forest_Southeastern_Brazil | Morellato_e_Leitão_Filho_1996_Biotropica_282_180_19 |
| Bignoniaceae | Dolichandra_unguis_cati | Flowering | 0 | 0 | | | | | | | | | | | -22.82916667 | -47.10916667 | SP | Campinas | Mata Atlantica | 1991 | 3 | Semideciduous_forest_Southeastern_Brazil | Morellato_e_Leitão_Filho_1996_Biotropica_282_180_19 |
| Bignoniaceae | Dolichandra_unguis_cati | Fruiting | 0 | 0 | | | | | | | | | | | -22.82916667 | -47.10916667 | SP | Campinas | Mata Atlantica | 1991 | 3 | Semideciduous_forest_Southeastern_Brazil | Morellato_e_Leitão_Filho_1996_Biotropica_282_180_19 |
| Bignoniaceae | Dolichandra_unguis_cati | Leaf_fall | | | | | | | | | | | | | -22.82916667 | -47.10916667 | SP | Campinas | Mata Atlantica | 1991 | 3 | Semideciduous_forest_Southeastern_Brazil | Morellato_e_Leitão_Filho_1996_Biotropica_282_180_19 |
| Bignoniaceae | Dolichandra_unguis_cati | Flushing | | | | | | | | | | | | | -22.82916667 | -47.10916667 | SP | Campinas | Mata Atlantica | 1991 | 3 | Semideciduous_forest_Southeastern_Brazil | Morellato_e_Leitão_Filho_1996_Biotropica_282_180_19 |
| Bignoniaceae | Jacaranda_caroba | Flowering | | | | | | | | | 1 | 1 | 0 | 0 | -21.61083333 | -47.61055556 | SP | Santa Rita do Passa Quatro | Cerrado | 1995 | 2 | Cerrado_Pé-de-Gigante_Reserve_SP_BRAZIL | Batalha_e_Mantovani_2000_Rev_Brasil_Biol_601_129_145 |
| Bignoniaceae | Jacaranda_caroba | Fruiting | | | | | | | | | 1 | 1 | 1 | 1 | -21.61083333 | -47.61055556 | SP | Santa Rita do Passa Quatro | Cerrado | 1995 | 2 | Cerrado_Pé-de-Gigante_Reserve_SP_BRAZIL | Batalha_e_Mantovani_2000_Rev_Brasil_Biol_601_129_145 |
| Bignoniaceae | Jacaranda_caroba | Leaf_fall | | | | | | | | | | | | | -21.61083333 | -47.61055556 | SP | Santa Rita do Passa Quatro | Cerrado | 1995 | 2 | Cerrado_Pé-de-Gigante_Reserve_SP_BRAZIL | Batalha_e_Mantovani_2000_Rev_Brasil_Biol_601_129_145 |
| Bignoniaceae | Jacaranda_caroba | Flushing | | | | | | | | | | | | | -21.61083333 | -47.61055556 | SP | Santa Rita do Passa Quatro | Cerrado | 1995 | 2 | Cerrado_Pé-de-Gigante_Reserve_SP_BRAZIL | Batalha_e_Mantovani_2000_Rev_Brasil_Biol_601_129_145 |
| Bignoniaceae | Jacaranda_caroba | Flowering | 0 | 0 | 1 | 1 | 1 | 1 | 1 | 1 | 1 | 1 | 0 | 0 | -21.61083333 | -47.61055556 | SP | Santa Rita do Passa Quatro | Cerrado | 1996 | 2 | Cerrado_Pé-de-Gigante_Reserve_SP_BRAZIL | Batalha_e_Mantovani_2000_Rev_Brasil_Biol_601_129_145 |
| Bignoniaceae | Jacaranda_caroba | Fruiting | 1 | 0 | 0 | 0 | 0 | 0 | 1 | 1 | 1 | 1 | 1 | 1 | -21.61083333 | -47.61055556 | SP | Santa Rita do Passa Quatro | Cerrado | 1996 | 2 | Cerrado_Pé-de-Gigante_Reserve_SP_BRAZIL | Batalha_e_Mantovani_2000_Rev_Brasil_Biol_601_129_145 |
| Bignoniaceae | Jacaranda_caroba | Leaf_fall | | | | | | | | | | | | | -21.61083333 | -47.61055556 | SP | Santa Rita do Passa Quatro | Cerrado | 1996 | 2 | Cerrado_Pé-de-Gigante_Reserve_SP_BRAZIL | Batalha_e_Mantovani_2000_Rev_Brasil_Biol_601_129_145 |
| Bignoniaceae | Jacaranda_caroba | Flushing | | | | | | | | | | | | | -21.61083333 | -47.61055556 | SP | Santa Rita do Passa Quatro | Cerrado | 1996 | 2 | Cerrado_Pé-de-Gigante_Reserve_SP_BRAZIL | Batalha_e_Mantovani_2000_Rev_Brasil_Biol_601_129_145 |
| Bignoniaceae | Jacaranda_caroba | Flowering | 0 | 0 | | | | | | | | | | | -21.61083333 | -47.61055556 | SP | Santa Rita do Passa Quatro | Cerrado | 1997 | 2 | Cerrado_Pé-de-Gigante_Reserve_SP_BRAZIL | Batalha_e_Mantovani_2000_Rev_Brasil_Biol_601_129_145 |
| Bignoniaceae | Jacaranda_caroba | Fruiting | 1 | 0 | | | | | | | | | | | -21.61083333 | -47.61055556 | SP | Santa Rita do Passa Quatro | Cerrado | 1997 | 2 | Cerrado_Pé-de-Gigante_Reserve_SP_BRAZIL | Batalha_e_Mantovani_2000_Rev_Brasil_Biol_601_129_145 |
| Bignoniaceae | Jacaranda_caroba | Leaf_fall | | | | | | | | | | | | | -21.61083333 | -47.61055556 | SP | Santa Rita do Passa Quatro | Cerrado | 1997 | 2 | Cerrado_Pé-de-Gigante_Reserve_SP_BRAZIL | Batalha_e_Mantovani_2000_Rev_Brasil_Biol_601_129_145 |
| Bignoniaceae | Jacaranda_caroba | Flushing | | | | | | | | | | | | | -21.61083333 | -47.61055556 | SP | Santa Rita do Passa Quatro | Cerrado | 1997 | 2 | Cerrado_Pé-de-Gigante_Reserve_SP_BRAZIL | Batalha_e_Mantovani_2000_Rev_Brasil_Biol_601_129_145 |
| Bignoniaceae | Tabebuia_aurea | Flowering | | | | | | | | | 1 | 1 | 0 | 0 | -21.61083333 | -47.61055556 | SP | Santa Rita do Passa Quatro | Cerrado | 1995 | 2 | Cerrado_Pé-de-Gigante_Reserve_SP_BRAZIL | Batalha_e_Mantovani_2000_Rev_Brasil_Biol_601_129_145 |
| Bignoniaceae | Tabebuia_aurea | Fruiting | | | | | | | | | | | | | -21.61083333 | -47.61055556 | SP | Santa Rita do Passa Quatro | Cerrado | 1995 | 2 | Cerrado_Pé-de-Gigante_Reserve_SP_BRAZIL | Batalha_e_Mantovani_2000_Rev_Brasil_Biol_601_129_145 |
| Bignoniaceae | Tabebuia_aurea | Leaf_fall | | | | | | | | | | | | | -21.61083333 | -47.61055556 | SP | Santa Rita do Passa Quatro | Cerrado | 1995 | 2 | Cerrado_Pé-de-Gigante_Reserve_SP_BRAZIL | Batalha_e_Mantovani_2000_Rev_Brasil_Biol_601_129_145 |
| Bignoniaceae | Tabebuia_aurea | Flushing | | | | | | | | | | | | | -21.61083333 | -47.61055556 | SP | Santa Rita do Passa Quatro | Cerrado | 1995 | 2 | Cerrado_Pé-de-Gigante_Reserve_SP_BRAZIL | Batalha_e_Mantovani_2000_Rev_Brasil_Biol_601_129_145 |
| Bignoniaceae | Tabebuia_aurea | Flowering | 0 | 0 | 0 | 0 | 0 | 0 | 0 | 0 | 1 | 1 | 0 | 0 | -21.61083333 | -47.61055556 | SP | Santa Rita do Passa Quatro | Cerrado | 1996 | 2 | Cerrado_Pé-de-Gigante_Reserve_SP_BRAZIL | Batalha_e_Mantovani_2000_Rev_Brasil_Biol_601_129_145 |
| Bignoniaceae | Tabebuia_aurea | Fruiting | | | | | | | | | | | | | -21.61083333 | -47.61055556 | SP | Santa Rita do Passa Quatro | Cerrado | 1996 | 2 | Cerrado_Pé-de-Gigante_Reserve_SP_BRAZIL | Batalha_e_Mantovani_2000_Rev_Brasil_Biol_601_129_145 |
| Bignoniaceae | Tabebuia_aurea | Leaf_fall | | | | | | | | | | | | | -21.61083333 | -47.61055556 | SP | Santa Rita do Passa Quatro | Cerrado | 1996 | 2 | Cerrado_Pé-de-Gigante_Reserve_SP_BRAZIL | Batalha_e_Mantovani_2000_Rev_Brasil_Biol_601_129_145 |
| Bignoniaceae | Tabebuia_aurea | Flushing | | | | | | | | | | | | | -21.61083333 | -47.61055556 | SP | Santa Rita do Passa Quatro | Cerrado | 1996 | 2 | Cerrado_Pé-de-Gigante_Reserve_SP_BRAZIL | Batalha_e_Mantovani_2000_Rev_Brasil_Biol_601_129_145 |
| Bignoniaceae | Tabebuia_aurea | Flowering | 0 | 0 | | | | | | | | | | | -21.61083333 | -47.61055556 | SP | Santa Rita do Passa Quatro | Cerrado | 1997 | 2 | Cerrado_Pé-de-Gigante_Reserve_SP_BRAZIL | Batalha_e_Mantovani_2000_Rev_Brasil_Biol_601_129_145 |
| Bignoniaceae | Tabebuia_aurea | Fruiting | | | | | | | | | | | | | -21.61083333 | -47.61055556 | SP | Santa Rita do Passa Quatro | Cerrado | 1997 | 2 | Cerrado_Pé-de-Gigante_Reserve_SP_BRAZIL | Batalha_e_Mantovani_2000_Rev_Brasil_Biol_601_129_145 |
| Bignoniaceae | Tabebuia_aurea | Leaf_fall | | | | | | | | | | | | | -21.61083333 | -47.61055556 | SP | Santa Rita do Passa Quatro | Cerrado | 1997 | 2 | Cerrado_Pé-de-Gigante_Reserve_SP_BRAZIL | Batalha_e_Mantovani_2000_Rev_Brasil_Biol_601_129_145 |

| | | | | | | | | | | | | | | | | | | | | | | |
|---|---|---|---|---|---|---|---|---|---|---|---|---|---|---|---|---|---|---|---|---|---|---|---|
| Bignoniaceae | Tabebuia_aurea | Flushing | | | | | | | | | | | | | -21.61083333 | -47.61055556 | SP | Santa Rita do Passa Quatro | Cerrado | 1997 | 2 | Cerrado_Pé-de-Gigante_Reserve_SP_BRAZIL | Batalha_e_Mantovani_2000_Rev_Brasil_Biol_601_129_145 |
| Chrysobalanaceae | Acioa_longipendula | Flowering | 0 | 0 | 0 | 0 | 0 | 1 | 1 | 1 | 1 | 1 | 1 | 0 | -3.13333333 | -59.87777777 | AM | Manaus | Amazonia | 1970_1994 | 25 | Tropical_rainforest_Northern_Brazil | Ruiz_and_Alencar_1999_Acta_Amazonica_29_223_242 |
| Chrysobalanaceae | Acioa_longipendula | Flower_buds | 0 | 0 | 0 | 0 | 0 | 1 | 1 | 1 | 1 | 1 | 1 | 0 | -3.13333333 | -59.87777777 | AM | Manaus | Amazonia | 1970_1994 | 25 | Tropical_rainforest_Northern_Brazil | Ruiz_and_Alencar_1999_Acta_Amazonica_29_223_242 |
| Chrysobalanaceae | Acioa_longipendula | Fruiting | 1 | 1 | 1 | 1 | 0 | 0 | 0 | 1 | 1 | 1 | 1 | 1 | -3.13333333 | -59.87777777 | AM | Manaus | Amazonia | 1970_1994 | 25 | Tropical_rainforest_Northern_Brazil | Ruiz_and_Alencar_1999_Acta_Amazonica_29_223_242 |
| Chrysobalanaceae | Acioa_longipendula | Leaf_fall | 0 | 0 | 1 | 0 | 0 | 0 | 0 | 0 | 0 | 0 | 0 | 0 | -3.13333333 | -59.87777777 | AM | Manaus | Amazonia | 1970_1994 | 25 | Tropical_rainforest_Northern_Brazil | Ruiz_and_Alencar_1999_Acta_Amazonica_29_223_242 |
| Chrysobalanaceae | Acioa_longipendula | Flushing | 1 | 1 | 1 | 1 | 1 | 1 | 1 | 1 | 1 | 1 | 1 | 1 | -3.13333333 | -59.87777777 | AM | Manaus | Amazonia | 1970_1994 | 25 | Tropical_rainforest_Northern_Brazil | Ruiz_and_Alencar_1999_Acta_Amazonica_29_223_242 |
| Chrysobalanaceae | Licania_heteromorpha | Flowering | 1 | 1 | 1 | 1 | 1 | 1 | 1 | 0 | 1 | 1 | 1 | 1 | -3.13333333 | -59.87777777 | AM | Manaus | Amazonia | 1970_1994 | 25 | Tropical_rainforest_Northern_Brazil | Ruiz_and_Alencar_1999_Acta_Amazonica_29_223_242 |
| Chrysobalanaceae | Licania_heteromorpha | Flower_buds | 1 | 1 | 0 | 0 | 1 | 1 | 0 | 1 | 1 | 1 | 1 | 1 | -3.13333333 | -59.87777777 | AM | Manaus | Amazonia | 1970_1994 | 25 | Tropical_rainforest_Northern_Brazil | Ruiz_and_Alencar_1999_Acta_Amazonica_29_223_242 |
| Chrysobalanaceae | Licania_heteromorpha | Fruiting | 1 | 1 | 1 | 1 | 1 | 1 | 1 | 1 | 1 | 1 | 1 | 1 | -3.13333333 | -59.87777777 | AM | Manaus | Amazonia | 1970_1994 | 25 | Tropical_rainforest_Northern_Brazil | Ruiz_and_Alencar_1999_Acta_Amazonica_29_223_242 |
| Chrysobalanaceae | Licania_heteromorpha | Leaf_fall | 0 | 0 | 0 | 0 | 0 | 0 | 0 | 0 | 0 | 0 | 0 | 0 | -3.13333333 | -59.87777777 | AM | Manaus | Amazonia | 1970_1994 | 25 | Tropical_rainforest_Northern_Brazil | Ruiz_and_Alencar_1999_Acta_Amazonica_29_223_242 |
| Chrysobalanaceae | Licania_heteromorpha | Flushing | 0 | 0 | 1 | 1 | 1 | 0 | 1 | 1 | 1 | 1 | 1 | 1 | -3.13333333 | -59.87777777 | AM | Manaus | Amazonia | 1970_1994 | 25 | Tropical_rainforest_Northern_Brazil | Ruiz_and_Alencar_1999_Acta_Amazonica_29_223_242 |
| Chrysobalanaceae | Licania_octandra | Flowering | 0 | 0 | 0 | 0 | 0 | 1 | 1 | 1 | 0 | 0 | 1 | 0 | -3.13333333 | -59.87777777 | AM | Manaus | Amazonia | 1970_1994 | 25 | Tropical_rainforest_Northern_Brazil | Ruiz_and_Alencar_1999_Acta_Amazonica_29_223_242 |
| Chrysobalanaceae | Licania_octandra | Flower_buds | 0 | 1 | 0 | 1 | 1 | 1 | 1 | 1 | 1 | 1 | 1 | 0 | -3.13333333 | -59.87777777 | AM | Manaus | Amazonia | 1970_1994 | 25 | Tropical_rainforest_Northern_Brazil | Ruiz_and_Alencar_1999_Acta_Amazonica_29_223_242 |
| Chrysobalanaceae | Licania_octandra | Fruiting | 1 | 1 | 0 | 0 | 0 | 1 | 1 | 1 | 1 | 1 | 1 | 1 | -3.13333333 | -59.87777777 | AM | Manaus | Amazonia | 1970_1994 | 25 | Tropical_rainforest_Northern_Brazil | Ruiz_and_Alencar_1999_Acta_Amazonica_29_223_242 |
| Chrysobalanaceae | Licania_octandra | Leaf_fall | 0 | 0 | 0 | 0 | 0 | 0 | 0 | 0 | 0 | 0 | 1 | 0 | -3.13333333 | -59.87777777 | AM | Manaus | Amazonia | 1970_1994 | 25 | Tropical_rainforest_Northern_Brazil | Ruiz_and_Alencar_1999_Acta_Amazonica_29_223_242 |
| Chrysobalanaceae | Licania_octandra | Flushing | 0 | 0 | 0 | 0 | 0 | 1 | 1 | 1 | 1 | 1 | 0 | 0 | -3.13333333 | -59.87777777 | AM | Manaus | Amazonia | 1970_1994 | 25 | Tropical_rainforest_Northern_Brazil | Ruiz_and_Alencar_1999_Acta_Amazonica_29_223_242 |
| Fabaceae | Bauhinia_acuruana | Flowering | | | | 1 | 1 | 1 | 1 | 1 | 0 | 0 | 1 | 0 | -5.14583333 | -40.92861111 | CE | Crateus | Caatinga | 2004 | 1 | Carrasco_vegetation_Serra_das_Almas_Reserve | Vasconcelos_et_al_2010_Biodivers_Conserv_19_2263_2289 |
| Fabaceae | Bauhinia_acuruana | Fruiting | | | | 1 | 1 | 1 | 1 | 1 | 1 | 1 | 1 | 0 | -5.14583333 | -40.92861111 | CE | Crateus | Caatinga | 2004 | 1 | Carrasco_vegetation_Serra_das_Almas_Reserve | Vasconcelos_et_al_2010_Biodivers_Conserv_19_2263_2289 |
| Fabaceae | Bauhinia_acuruana | Leaf_fall | | | | | | | | | | | | | -5.14583333 | -40.92861111 | CE | Crateus | Caatinga | 2004 | 1 | Carrasco_vegetation_Serra_das_Almas_Reserve | Vasconcelos_et_al_2010_Biodivers_Conserv_19_2263_2289 |
| Fabaceae | Bauhinia_acuruana | Flushing | | | | | | | | | | | | | -5.14583333 | -40.92861111 | CE | Crateus | Caatinga | 2004 | 1 | Carrasco_vegetation_Serra_das_Almas_Reserve | Vasconcelos_et_al_2010_Biodivers_Conserv_19_2263_2289 |
| Fabaceae | Bauhinia_acuruana | Flowering | 1 | 1 | 1 | | | | | | | | | | -5.14583333 | -40.92861111 | CE | Crateus | Caatinga | 2005 | 1 | Carrasco_vegetation_Serra_das_Almas_Reserve | Vasconcelos_et_al_2010_Biodivers_Conserv_19_2263_2289 |
| Fabaceae | Bauhinia_acuruana | Fruiting | 1 | 1 | 1 | | | | | | | | | | -5.14583333 | -40.92861111 | CE | Crateus | Caatinga | 2005 | 1 | Carrasco_vegetation_Serra_das_Almas_Reserve | Vasconcelos_et_al_2010_Biodivers_Conserv_19_2263_2289 |
| Fabaceae | Bauhinia_acuruana | Leaf_fall | | | | | | | | | | | | | -5.14583333 | -40.92861111 | CE | Crateus | Caatinga | 2005 | 1 | Carrasco_vegetation_Serra_das_Almas_Reserve | Vasconcelos_et_al_2010_Biodivers_Conserv_19_2263_2289 |
| Fabaceae | Bauhinia_acuruana | Flushing | | | | | | | | | | | | | -5.14583333 | -40.92861111 | CE | Crateus | Caatinga | 2005 | 1 | Carrasco_vegetation_Serra_das_Almas_Reserve | Vasconcelos_et_al_2010_Biodivers_Conserv_19_2263_2289 |
| Fabaceae | Chamaecrista_nictitans | Flowering | 1 | 1 | 0 | 1 | 1 | 0 | 1 | 0 | 1 | 0 | 0 | 0 | -7.47916667 | -36.905 | PB | Sao Jose dos Cordeiros | Caatinga | 2003 | 2 | Arboreal_and_shrub_Caatinga | Quirino_and_Machad0_Braz. J. Biol_74_62_71 |
| Fabaceae | Chamaecrista_nictitans | Fruiting | | | | | | | | | | | | | -7.47916667 | -36.905 | PB | Sao Jose dos Cordeiros | Caatinga | 2003 | 2 | Arboreal_and_shrub_Caatinga | Quirino_and_Machad0_Braz. J. Biol_74_62_71 |
| Fabaceae | Chamaecrista_nictitans | Leaf_fall | | | | | | | | | | | | | -7.47916667 | -36.905 | PB | Sao Jose dos Cordeiros | Caatinga | 2003 | 2 | Arboreal_and_shrub_Caatinga | Quirino_and_Machad0_Braz. J. Biol_74_62_71 |
| Fabaceae | Chamaecrista_nictitans | Flushing | | | | | | | | | | | | | -7.47916667 | -36.905 | PB | Sao Jose dos Cordeiros | Caatinga | 2003 | 2 | Arboreal_and_shrub_Caatinga | Quirino_and_Machad0_Braz. J. Biol_74_62_71 |
| Fabaceae | Chamaecrista_nictitans | Flowering | 0 | 1 | 1 | 0 | 1 | 1 | 1 | 1 | 1 | 0 | 0 | 0 | -7.47916667 | -36.905 | PB | Sao Jose dos Cordeiros | Caatinga | 2004 | 2 | Arboreal_and_shrub_Caatinga | Quirino_and_Machad0_Braz. J. Biol_74_62_71 |
| Fabaceae | Chamaecrista_nictitans | Fruiting | | | | | | | | | | | | | -7.47916667 | -36.905 | PB | Sao Jose dos Cordeiros | Caatinga | 2004 | 2 | Arboreal_and_shrub_Caatinga | Quirino_and_Machad0_Braz. J. Biol_74_62_71 |
| Fabaceae | Chamaecrista_nictitans | Leaf_fall | | | | | | | | | | | | | -7.47916667 | -36.905 | PB | Sao Jose dos Cordeiros | Caatinga | 2004 | 2 | Arboreal_and_shrub_Caatinga | Quirino_and_Machad0_Braz. J. Biol_74_62_71 |
| Fabaceae | Chamaecrista_nictitans | Flushing | | | | | | | | | | | | | -7.47916667 | -36.905 | PB | Sao Jose dos Cordeiros | Caatinga | 2004 | 2 | Arboreal_and_shrub_Caatinga | Quirino_and_Machad0_Braz. J. Biol_74_62_71 |
| Fabaceae | Chamaecrista_desvauxii | Flowering | 1 | 1 | 1 | 1 | 0 | 0 | 0 | 0 | 0 | 0 | 1 | 1 | -23.98583333 | -46.49583333 | SP | Sao Vicente | Mata Atlantica | 2006 | 2 | Coastal_plain_degraded_by_sand_mining_in_São_Paulo_State_Brazil | Fidalgo_et_al_2018_Hoehnea_451_93_102 |
| Fabaceae | Chamaecrista_desvauxii | Fruiting | 1 | 1 | 1 | 1 | 0 | 0 | 0 | 0 | 0 | 1 | 1 | 1 | -23.98583333 | -46.49583333 | SP | Sao Vicente | Mata Atlantica | 2006 | 2 | Coastal_plain_degraded_by_sand_mining_in_São_Paulo_State_Brazil | Fidalgo_et_al_2018_Hoehnea_451_93_102 |
| Fabaceae | Chamaecrista_desvauxii | Leaf_fall | | | | | | | | | | | | | -23.98583333 | -46.49583333 | SP | Sao Vicente | Mata Atlantica | 2006 | 2 | Coastal_plain_degraded_by_sand_mining_in_São_Paulo_State_Brazil | Fidalgo_et_al_2018_Hoehnea_451_93_102 |
| Fabaceae | Chamaecrista_desvauxii | Flushing | | | | | | | | | | | | | -23.98583333 | -46.49583333 | SP | Sao Vicente | Mata Atlantica | 2006 | 2 | Coastal_plain_degraded_by_sand_mining_in_São_Paulo_State_Brazil | Fidalgo_et_al_2018_Hoehnea_451_93_102 |
| Fabaceae | Chamaecrista_desvauxii | Flowering | 1 | 1 | 1 | 1 | 0 | 0 | 0 | 0 | 0 | 0 | 1 | 1 | -23.98583333 | -46.49583333 | SP | Sao Vicente | Mata Atlantica | 2007 | 2 | Coastal_plain_degraded_by_sand_mining_in_São_Paulo_State_Brazil | Fidalgo_et_al_2018_Hoehnea_451_93_102 |
| Fabaceae | Chamaecrista_desvauxii | Fruiting | 1 | 1 | 1 | 1 | 1 | 0 | 0 | 0 | 0 | 0 | 1 | 1 | -23.98583333 | -46.49583333 | SP | Sao Vicente | Mata Atlantica | 2007 | 2 | Coastal_plain_degraded_by_sand_mining_in_São_Paulo_State_Brazil | Fidalgo_et_al_2018_Hoehnea_451_93_102 |
| Fabaceae | Chamaecrista_desvauxii | Leaf_fall | | | | | | | | | | | | | -23.98583333 | -46.49583333 | SP | Sao Vicente | Mata Atlantica | 2007 | 2 | Coastal_plain_degraded_by_sand_mining_in_São_Paulo_State_Brazil | Fidalgo_et_al_2018_Hoehnea_451_93_102 |
| Fabaceae | Chamaecrista_desvauxii | Flushing | | | | | | | | | | | | | -23.98583333 | -46.49583333 | SP | Sao Vicente | Mata Atlantica | 2007 | 2 | Coastal_plain_degraded_by_sand_mining_in_São_Paulo_State_Brazil | Fidalgo_et_al_2018_Hoehnea_451_93_102 |
| Fabaceae | Chamaecrista_desvauxii | Flowering | 0 | 0 | 1 | 1 | 0 | | | | | | | | -23.98583333 | -46.49583333 | SP | Sao Vicente | Mata Atlantica | 2008 | 2 | Coastal_plain_degraded_by_sand_mining_in_São_Paulo_State_Brazil | Fidalgo_et_al_2018_Hoehnea_451_93_102 |
| Fabaceae | Chamaecrista_desvauxii | Fruiting | 1 | 0 | 1 | 0 | 0 | | | | | | | | -23.98583333 | -46.49583333 | SP | Sao Vicente | Mata Atlantica | 2008 | 2 | Coastal_plain_degraded_by_sand_mining_in_São_Paulo_State_Brazil | Fidalgo_et_al_2018_Hoehnea_451_93_102 |
| Fabaceae | Chamaecrista_desvauxii | Leaf_fall | | | | | | | | | | | | | -23.98583333 | -46.49583333 | SP | Sao Vicente | Mata Atlantica | 2008 | 2 | Coastal_plain_degraded_by_sand_mining_in_São_Paulo_State_Brazil | Fidalgo_et_al_2018_Hoehnea_451_93_102 |
| Fabaceae | Chamaecrista_desvauxii | Flushing | | | | | | | | | | | | | -23.98583333 | -46.49583333 | SP | Sao Vicente | Mata Atlantica | 2008 | 2 | Coastal_plain_degraded_by_sand_mining_in_São_Paulo_State_Brazil | Fidalgo_et_al_2018_Hoehnea_451_93_102 |
| Fabaceae | Chamaecrista_ramosa | Flowering | 1 | 1 | 1 | 1 | 1 | 1 | 0 | 0 | 0 | 1 | 1 | 1 | -19.365 | -43.53305556 | MG | Santana do Riacho | Cerrado | 1996_1997 | 2 | Campo_rupestre_Serra_do_Cipó_MG_southeast_Brazil | Madeira_&_Fernandes_1999_Journal_of_Tropical_Ecology_15463_479 |
| Fabaceae | Chamaecrista_ramosa | Fruiting | 1 | 1 | 1 | 1 | 1 | 1 | 0 | 0 | 1 | 1 | 1 | 1 | -19.365 | -43.53305556 | MG | Santana do Riacho | Cerrado | 1996_1997 | 2 | Campo_rupestre_Serra_do_Cipó_MG_southeast_Brazil | Madeira_&_Fernandes_1999_Journal_of_Tropical_Ecology_15463_479 |
| Fabaceae | Chamaecrista_ramosa | Leaf_fall | | | | | | | | | | | | | -19.365 | -43.53305556 | MG | Santana do Riacho | Cerrado | 1996_1997 | 2 | Campo_rupestre_Serra_do_Cipó_MG_southeast_Brazil | Madeira_&_Fernandes_1999_Journal_of_Tropical_Ecology_15463_479 |
| Fabaceae | Chamaecrista_ramosa | Flushing | | | | | | | | | | | | | -19.365 | -43.53305556 | MG | Santana do Riacho | Cerrado | 1996_1997 | 2 | Campo_rupestre_Serra_do_Cipó_MG_southeast_Brazil | Madeira_&_Fernandes_1999_Journal_of_Tropical_Ecology_15463_479 |
| Fabaceae | Senna_multijuga | Flowering | 1 | 1 | 1 | 0 | 0 | 0 | 0 | 0 | 0 | 0 | 0 | 0 | -25.32083333 | -45.70666667 | PR | Antonina | Mata Atlantica | 2007_2008 | 2 | Atlantic_forest_Rio_Cachoeira_Reserve_PR_southern_coast_of_Brazil | Cardoso_et_al_2012_Plant_Soil_354269_281 |
| Fabaceae | Senna_multijuga | Fruiting | 0 | 0 | 0 | 1 | 1 | 1 | 1 | 1 | 1 | 1 | 0 | 0 | -25.32083333 | -45.70666667 | PR | Antonina | Mata Atlantica | 2007_2008 | 2 | Atlantic_forest_Rio_Cachoeira_Reserve_PR_southern_coast_of_Brazil | Cardoso_et_al_2012_Plant_Soil_354269_281 |
| Fabaceae | Senna_multijuga | Leaf_fall | 0 | 0 | 0 | 1 | 1 | 1 | 1 | 0 | 0 | 0 | 0 | 0 | -25.32083333 | -45.70666667 | PR | Antonina | Mata Atlantica | 2007_2008 | 2 | Atlantic_forest_Rio_Cachoeira_Reserve_PR_southern_coast_of_Brazil | Cardoso_et_al_2012_Plant_Soil_354269_281 |
| Fabaceae | Senna_multijuga | Flushing | 1 | 0 | 0 | 0 | 0 | 0 | 0 | 1 | 1 | 1 | 1 | 1 | -25.32083333 | -45.70666667 | PR | Antonina | Mata Atlantica | 2007_2008 | 2 | Atlantic_forest_Rio_Cachoeira_Reserve_PR_southern_coast_of_Brazil | Cardoso_et_al_2012_Plant_Soil_354269_281 |
| Fabaceae | Senna_multijuga | Flowering | 1 | 1 | 1 | 0 | 0 | 0 | 0 | 0 | 0 | 0 | 0 | 0 | -25.32083333 | -45.70666667 | PR | Antonina | Mata Atlantica | 2007_2008 | 2 | Atlantic_forest_Rio_Cachoeira_Reserve_PR_southern_coast_of_Brazil | Cardoso_et_al_2012_Plant_Soil_354269_281 |
| Fabaceae | Senna_multijuga | Fruiting | 0 | 0 | 0 | 1 | 1 | 1 | 1 | 1 | 1 | 1 | 0 | 0 | -25.32083333 | -45.70666667 | PR | Antonina | Mata Atlantica | 2007_2008 | 2 | Atlantic_forest_Rio_Cachoeira_Reserve_PR_southern_coast_of_Brazil | Cardoso_et_al_2012_Plant_Soil_354269_281 |
| Fabaceae | Senna_multijuga | Leaf_fall | 0 | 0 | 0 | 1 | 1 | 1 | 1 | 0 | 0 | 0 | 0 | 0 | -25.32083333 | -45.70666667 | PR | Antonina | Mata Atlantica | 2007_2008 | 2 | Atlantic_forest_Rio_Cachoeira_Reserve_PR_southern_coast_of_Brazil | Cardoso_et_al_2012_Plant_Soil_354269_281 |
| Fabaceae | Senna_multijuga | Flushing | 1 | 0 | 0 | 0 | 0 | 0 | 0 | 1 | 1 | 1 | 1 | 1 | -25.32083333 | -45.70666667 | PR | Antonina | Mata Atlantica | 2007_2008 | 2 | Atlantic_forest_Rio_Cachoeira_Reserve_PR_southern_coast_of_Brazil | Cardoso_et_al_2012_Plant_Soil_354269_281 |
| Malpighiaceae | Banisteriopsis_argyrophylla | Flowering | | | | | | | | | 0 | 0 | 0 | 0 | -21.61083333 | -47.61055556 | SP | Santa Rita do Passa Quatro | Cerrado | 1995 | 2 | Cerrado_Pé-de-Gigante_Reserve_SP_BRAZIL | Batalha_e_Mantovani_2000_Rev_Brasil_Biol_601_129_145 |
| Malpighiaceae | Banisteriopsis_argyrophylla | Fruiting | | | | | | | | | | | | | -21.61083333 | -47.61055556 | SP | Santa Rita do Passa Quatro | Cerrado | 1995 | 2 | Cerrado_Pé-de-Gigante_Reserve_SP_BRAZIL | Batalha_e_Mantovani_2000_Rev_Brasil_Biol_601_129_145 |
| Malpighiaceae | Banisteriopsis_argyrophylla | Leaf_fall | | | | | | | | | | | | | -21.61083333 | -47.61055556 | SP | Santa Rita do Passa Quatro | Cerrado | 1995 | 2 | Cerrado_Pé-de-Gigante_Reserve_SP_BRAZIL | Batalha_e_Mantovani_2000_Rev_Brasil_Biol_601_129_145 |
| Malpighiaceae | Banisteriopsis_argyrophylla | Flushing | | | | | | | | | | | | | -21.61083333 | -47.61055556 | SP | Santa Rita do Passa Quatro | Cerrado | 1995 | 2 | Cerrado_Pé-de-Gigante_Reserve_SP_BRAZIL | Batalha_e_Mantovani_2000_Rev_Brasil_Biol_601_129_145 |
| Malpighiaceae | Banisteriopsis_argyrophylla | Flowering | 0 | 0 | 0 | 1 | 1 | 0 | 0 | 0 | 0 | 0 | 0 | 0 | -21.61083333 | -47.61055556 | SP | Santa Rita do Passa Quatro | Cerrado | 1996 | 2 | Cerrado_Pé-de-Gigante_Reserve_SP_BRAZIL | Batalha_e_Mantovani_2000_Rev_Brasil_Biol_601_129_145 |
| Malpighiaceae | Banisteriopsis_argyrophylla | Fruiting | | | | | | | | | | | | | -21.61083333 | -47.61055556 | SP | Santa Rita do Passa Quatro | Cerrado | 1996 | 2 | Cerrado_Pé-de-Gigante_Reserve_SP_BRAZIL | Batalha_e_Mantovani_2000_Rev_Brasil_Biol_601_129_145 |
| Malpighiaceae | Banisteriopsis_argyrophylla | Leaf_fall | | | | | | | | | | | | | -21.61083333 | -47.61055556 | SP | Santa Rita do Passa Quatro | Cerrado | 1996 | 2 | Cerrado_Pé-de-Gigante_Reserve_SP_BRAZIL | Batalha_e_Mantovani_2000_Rev_Brasil_Biol_601_129_145 |
| Malpighiaceae | Banisteriopsis_argyrophylla | Flushing | | | | | | | | | | | | | -21.61083333 | -47.61055556 | SP | Santa Rita do Passa Quatro | Cerrado | 1996 | 2 | Cerrado_Pé-de-Gigante_Reserve_SP_BRAZIL | Batalha_e_Mantovani_2000_Rev_Brasil_Biol_601_129_145 |
| Malpighiaceae | Banisteriopsis_argyrophylla | Flowering | 0 | 0 | | | | | | | | | | | -21.61083333 | -47.61055556 | SP | Santa Rita do Passa Quatro | Cerrado | 1997 | 2 | Cerrado_Pé-de-Gigante_Reserve_SP_BRAZIL | Batalha_e_Mantovani_2000_Rev_Brasil_Biol_601_129_145 |
| Malpighiaceae | Banisteriopsis_argyrophylla | Fruiting | | | | | | | | | | | | | -21.61083333 | -47.61055556 | SP | Santa Rita do Passa Quatro | Cerrado | 1997 | 2 | Cerrado_Pé-de-Gigante_Reserve_SP_BRAZIL | Batalha_e_Mantovani_2000_Rev_Brasil_Biol_601_129_145 |
| Malpighiaceae | Banisteriopsis_argyrophylla | Leaf_fall | | | | | | | | | | | | | -21.61083333 | -47.61055556 | SP | Santa Rita do Passa Quatro | Cerrado | 1997 | 2 | Cerrado_Pé-de-Gigante_Reserve_SP_BRAZIL | Batalha_e_Mantovani_2000_Rev_Brasil_Biol_601_129_145 |
| Malpighiaceae | Banisteriopsis_argyrophylla | Flushing | | | | | | | | | | | | | -21.61083333 | -47.61055556 | SP | Santa Rita do Passa Quatro | Cerrado | 1997 | 2 | Cerrado_Pé-de-Gigante_Reserve_SP_BRAZIL | Batalha_e_Mantovani_2000_Rev_Brasil_Biol_601_129_145 |
| Malpighiaceae | Banisteriopsis_argyrophylla | Flowering | | | 1 | 1 | 1 | 0 | 0 | 0 | 0 | 0 | 0 | 0 | -22.82916667 | -47.10916667 | SP | Campinas | Mata Atlantica | 1988 | 3 | Semideciduous_forest_Southeastern_Brazil | Morellato_e_Leitão_Filho_1996_Biotropica_282_180_19 |
| Malpighiaceae | Banisteriopsis_argyrophylla | Fruiting | | | 0 | 0 | 0 | 1 | 1 | 0 | 0 | 0 | 0 | 0 | -22.82916667 | -47.10916667 | SP | Campinas | Mata Atlantica | 1988 | 3 | Semideciduous_forest_Southeastern_Brazil | Morellato_e_Leitão_Filho_1996_Biotropica_282_180_19 |
| Malpighiaceae | Banisteriopsis_argyrophylla | Leaf_fall | | | | | | | | | | | | | -22.82916667 | -47.10916667 | SP | Campinas | Mata Atlantica | 1988 | 3 | Semideciduous_forest_Southeastern_Brazil | Morellato_e_Leitão_Filho_1996_Biotropica_282_180_19 |
| Malpighiaceae | Banisteriopsis_argyrophylla | Flushing | | | | | | | | | | | | | -22.82916667 | -47.10916667 | SP | Campinas | Mata Atlantica | 1988 | 3 | Semideciduous_forest_Southeastern_Brazil | Morellato_e_Leitão_Filho_1996_Biotropica_282_180_19 |
| Malpighiaceae | Banisteriopsis_argyrophylla | Flowering | 0 | 0 | 1 | 1 | 1 | 0 | 0 | 0 | 0 | 0 | 0 | 0 | -22.82916667 | -47.10916667 | SP | Campinas | Mata Atlantica | 1989 | 3 | Semideciduous_forest_Southeastern_Brazil | Morellato_e_Leitão_Filho_1996_Biotropica_282_180_19 |
| Malpighiaceae | Banisteriopsis_argyrophylla | Fruiting | 0 | 0 | 0 | 0 | 0 | 1 | 1 | 0 | 0 | 0 | 0 | 0 | -22.82916667 | -47.10916667 | SP | Campinas | Mata Atlantica | 1989 | 3 | Semideciduous_forest_Southeastern_Brazil | Morellato_e_Leitão_Filho_1996_Biotropica_282_180_19 |
| Malpighiaceae | Banisteriopsis_argyrophylla | Leaf_fall | | | | | | | | | | | | | -22.82916667 | -47.10916667 | SP | Campinas | Mata Atlantica | 1989 | 3 | Semideciduous_forest_Southeastern_Brazil | Morellato_e_Leitão_Filho_1996_Biotropica_282_180_19 |
| Malpighiaceae | Banisteriopsis_argyrophylla | Flushing | | | | | | | | | | | | | -22.82916667 | -47.10916667 | SP | Campinas | Mata Atlantica | 1989 | 3 | Semideciduous_forest_Southeastern_Brazil | Morellato_e_Leitão_Filho_1996_Biotropica_282_180_19 |
| Malpighiaceae | Banisteriopsis_argyrophylla | Flowering | 0 | 0 | 1 | 1 | 1 | 0 | 0 | 0 | 0 | 0 | 0 | 0 | -22.82916667 | -47.10916667 | SP | Campinas | Mata Atlantica | 1990 | 3 | Semideciduous_forest_Southeastern_Brazil | Morellato_e_Leitão_Filho_1996_Biotropica_282_180_19 |
| Malpighiaceae | Banisteriopsis_argyrophylla | Fruiting | 0 | 0 | 0 | 0 | 0 | 1 | 1 | 0 | 0 | 0 | 0 | 0 | -22.82916667 | -47.10916667 | SP | Campinas | Mata Atlantica | 1990 | 3 | Semideciduous_forest_Southeastern_Brazil | Morellato_e_Leitão_Filho_1996_Biotropica_282_180_19 |
| Malpighiaceae | Banisteriopsis_argyrophylla | Leaf_fall | | | | | | | | | | | | | -22.82916667 | -47.10916667 | SP | Campinas | Mata Atlantica | 1990 | 3 | Semideciduous_forest_Southeastern_Brazil | Morellato_e_Leitão_Filho_1996_Biotropica_282_180_19 |
| Malpighiaceae | Banisteriopsis_argyrophylla | Flushing | | | | | | | | | | | | | -22.82916667 | -47.10916667 | SP | Campinas | Mata Atlantica | 1990 | 3 | Semideciduous_forest_Southeastern_Brazil | Morellato_e_Leitão_Filho_1996_Biotropica_282_180_19 |
| Malpighiaceae | Banisteriopsis_argyrophylla | Flowering | 0 | 0 | | | | | | | | | | | -22.82916667 | -47.10916667 | SP | Campinas | Mata Atlantica | 1991 | 3 | Semideciduous_forest_Southeastern_Brazil | Morellato_e_Leitão_Filho_1996_Biotropica_282_180_19 |

| | | | | | | | | | | | | | | | | | | | | | | | |
|---|---|---|---|---|---|---|---|---|---|---|---|---|---|---|---|---|---|---|---|---|---|---|---|
| Malpighiaceae | Banisteriopsis_argyrophylla | Fruiting | 0 | 0 | | | | | | | | | | | -22.82916667 | -47.10916667 | SP | Campinas | Mata Atlantica | 1991 | 3 | Semideciduous_forest_Southeastern_Brazil | Morellato_e_Leitão_Filho_1996_Biotropica_282_180_19 |
| Malpighiaceae | Banisteriopsis_argyrophylla | Leaf_fall | | | | | | | | | | | | | -22.82916667 | -47.10916667 | SP | Campinas | Mata Atlantica | 1991 | 3 | Semideciduous_forest_Southeastern_Brazil | Morellato_e_Leitão_Filho_1996_Biotropica_282_180_19 |
| Malpighiaceae | Banisteriopsis_argyrophylla | Flushing | | | | | | | | | | | | | -22.82916667 | -47.10916667 | SP | Campinas | Mata Atlantica | 1991 | 3 | Semideciduous_forest_Southeastern_Brazil | Morellato_e_Leitão_Filho_1996_Biotropica_282_180_19 |
| Malpighiaceae | Banisteriopsis_campestris | Flowering | | | | | | | | | 0 | 0 | 0 | 0 | -21.61083333 | -47.61055556 | SP | Santa Rita do Passa Quatro | Cerrado | 1995 | 2 | Cerrado_Pé-de-Gigante_Reserve_SP_BRAZIL | Batalha_e_Mantovani_2000_Rev_Brasil_Biol_601_129_145 |
| Malpighiaceae | Banisteriopsis_campestris | Fruiting | | | | | | | | | 1 | 1 | 0 | 0 | -21.61083333 | -47.61055556 | SP | Santa Rita do Passa Quatro | Cerrado | 1995 | 2 | Cerrado_Pé-de-Gigante_Reserve_SP_BRAZIL | Batalha_e_Mantovani_2000_Rev_Brasil_Biol_601_129_145 |
| Malpighiaceae | Banisteriopsis_campestris | Leaf_fall | | | | | | | | | | | | | -21.61083333 | -47.61055556 | SP | Santa Rita do Passa Quatro | Cerrado | 1995 | 2 | Cerrado_Pé-de-Gigante_Reserve_SP_BRAZIL | Batalha_e_Mantovani_2000_Rev_Brasil_Biol_601_129_145 |
| Malpighiaceae | Banisteriopsis_campestris | Flushing | | | | | | | | | | | | | -21.61083333 | -47.61055556 | SP | Santa Rita do Passa Quatro | Cerrado | 1995 | 2 | Cerrado_Pé-de-Gigante_Reserve_SP_BRAZIL | Batalha_e_Mantovani_2000_Rev_Brasil_Biol_601_129_145 |
| Malpighiaceae | Banisteriopsis_campestris | Flowering | 1 | 1 | 1 | 1 | 1 | 1 | 0 | 0 | 0 | 0 | 0 | 0 | -21.61083333 | -47.61055556 | SP | Santa Rita do Passa Quatro | Cerrado | 1996 | 2 | Cerrado_Pé-de-Gigante_Reserve_SP_BRAZIL | Batalha_e_Mantovani_2000_Rev_Brasil_Biol_601_129_145 |
| Malpighiaceae | Banisteriopsis_campestris | Fruiting | 0 | 1 | 1 | 1 | 1 | 1 | 1 | 1 | 1 | 1 | 0 | 0 | -21.61083333 | -47.61055556 | SP | Santa Rita do Passa Quatro | Cerrado | 1996 | 2 | Cerrado_Pé-de-Gigante_Reserve_SP_BRAZIL | Batalha_e_Mantovani_2000_Rev_Brasil_Biol_601_129_145 |
| Malpighiaceae | Banisteriopsis_campestris | Leaf_fall | | | | | | | | | | | | | -21.61083333 | -47.61055556 | SP | Santa Rita do Passa Quatro | Cerrado | 1996 | 2 | Cerrado_Pé-de-Gigante_Reserve_SP_BRAZIL | Batalha_e_Mantovani_2000_Rev_Brasil_Biol_601_129_145 |
| Malpighiaceae | Banisteriopsis_campestris | Flushing | | | | | | | | | | | | | -21.61083333 | -47.61055556 | SP | Santa Rita do Passa Quatro | Cerrado | 1996 | 2 | Cerrado_Pé-de-Gigante_Reserve_SP_BRAZIL | Batalha_e_Mantovani_2000_Rev_Brasil_Biol_601_129_145 |
| Malpighiaceae | Banisteriopsis_campestris | Flowering | 1 | 1 | | | | | | | | | | | -21.61083333 | -47.61055556 | SP | Santa Rita do Passa Quatro | Cerrado | 1997 | 2 | Cerrado_Pé-de-Gigante_Reserve_SP_BRAZIL | Batalha_e_Mantovani_2000_Rev_Brasil_Biol_601_129_145 |
| Malpighiaceae | Banisteriopsis_campestris | Fruiting | 0 | 1 | | | | | | | | | | | -21.61083333 | -47.61055556 | SP | Santa Rita do Passa Quatro | Cerrado | 1997 | 2 | Cerrado_Pé-de-Gigante_Reserve_SP_BRAZIL | Batalha_e_Mantovani_2000_Rev_Brasil_Biol_601_129_145 |
| Malpighiaceae | Banisteriopsis_campestris | Leaf_fall | | | | | | | | | | | | | -21.61083333 | -47.61055556 | SP | Santa Rita do Passa Quatro | Cerrado | 1997 | 2 | Cerrado_Pé-de-Gigante_Reserve_SP_BRAZIL | Batalha_e_Mantovani_2000_Rev_Brasil_Biol_601_129_145 |
| Malpighiaceae | Banisteriopsis_campestris | Flushing | | | | | | | | | | | | | -21.61083333 | -47.61055556 | SP | Santa Rita do Passa Quatro | Cerrado | 1997 | 2 | Cerrado_Pé-de-Gigante_Reserve_SP_BRAZIL | Batalha_e_Mantovani_2000_Rev_Brasil_Biol_601_129_145 |
| Malpighiaceae | Banisteriopsis_laevifolia | Flowering | | | | | | | | | 0 | 0 | 0 | 1 | -21.61083333 | -47.61055556 | SP | Santa Rita do Passa Quatro | Cerrado | 1995 | 2 | Cerrado_Pé-de-Gigante_Reserve_SP_BRAZIL | Batalha_e_Mantovani_2000_Rev_Brasil_Biol_601_129_145 |
| Malpighiaceae | Banisteriopsis_laevifolia | Fruiting | | | | | | | | | 1 | 0 | 0 | 0 | -21.61083333 | -47.61055556 | SP | Santa Rita do Passa Quatro | Cerrado | 1995 | 2 | Cerrado_Pé-de-Gigante_Reserve_SP_BRAZIL | Batalha_e_Mantovani_2000_Rev_Brasil_Biol_601_129_145 |
| Malpighiaceae | Banisteriopsis_laevifolia | Leaf_fall | | | | | | | | | | | | | -21.61083333 | -47.61055556 | SP | Santa Rita do Passa Quatro | Cerrado | 1995 | 2 | Cerrado_Pé-de-Gigante_Reserve_SP_BRAZIL | Batalha_e_Mantovani_2000_Rev_Brasil_Biol_601_129_145 |
| Malpighiaceae | Banisteriopsis_laevifolia | Flushing | | | | | | | | | | | | | -21.61083333 | -47.61055556 | SP | Santa Rita do Passa Quatro | Cerrado | 1995 | 2 | Cerrado_Pé-de-Gigante_Reserve_SP_BRAZIL | Batalha_e_Mantovani_2000_Rev_Brasil_Biol_601_129_145 |
| Malpighiaceae | Banisteriopsis_laevifolia | Flowering | 1 | 1 | 1 | 1 | 1 | 0 | 0 | 0 | 0 | 0 | 0 | 1 | -21.61083333 | -47.61055556 | SP | Santa Rita do Passa Quatro | Cerrado | 1996 | 2 | Cerrado_Pé-de-Gigante_Reserve_SP_BRAZIL | Batalha_e_Mantovani_2000_Rev_Brasil_Biol_601_129_145 |
| Malpighiaceae | Banisteriopsis_laevifolia | Fruiting | 0 | 0 | 0 | 0 | 1 | 1 | 1 | 1 | 1 | 0 | 0 | 0 | -21.61083333 | -47.61055556 | SP | Santa Rita do Passa Quatro | Cerrado | 1996 | 2 | Cerrado_Pé-de-Gigante_Reserve_SP_BRAZIL | Batalha_e_Mantovani_2000_Rev_Brasil_Biol_601_129_145 |
| Malpighiaceae | Banisteriopsis_laevifolia | Leaf_fall | | | | | | | | | | | | | -21.61083333 | -47.61055556 | SP | Santa Rita do Passa Quatro | Cerrado | 1996 | 2 | Cerrado_Pé-de-Gigante_Reserve_SP_BRAZIL | Batalha_e_Mantovani_2000_Rev_Brasil_Biol_601_129_145 |
| Malpighiaceae | Banisteriopsis_laevifolia | Flushing | | | | | | | | | | | | | -21.61083333 | -47.61055556 | SP | Santa Rita do Passa Quatro | Cerrado | 1996 | 2 | Cerrado_Pé-de-Gigante_Reserve_SP_BRAZIL | Batalha_e_Mantovani_2000_Rev_Brasil_Biol_601_129_145 |
| Malpighiaceae | Banisteriopsis_laevifolia | Flowering | 1 | 1 | | | | | | | | | | | -21.61083333 | -47.61055556 | SP | Santa Rita do Passa Quatro | Cerrado | 1997 | 2 | Cerrado_Pé-de-Gigante_Reserve_SP_BRAZIL | Batalha_e_Mantovani_2000_Rev_Brasil_Biol_601_129_145 |
| Malpighiaceae | Banisteriopsis_laevifolia | Fruiting | 0 | 0 | | | | | | | | | | | -21.61083333 | -47.61055556 | SP | Santa Rita do Passa Quatro | Cerrado | 1997 | 2 | Cerrado_Pé-de-Gigante_Reserve_SP_BRAZIL | Batalha_e_Mantovani_2000_Rev_Brasil_Biol_601_129_145 |
| Malpighiaceae | Banisteriopsis_laevifolia | Leaf_fall | | | | | | | | | | | | | -21.61083333 | -47.61055556 | SP | Santa Rita do Passa Quatro | Cerrado | 1997 | 2 | Cerrado_Pé-de-Gigante_Reserve_SP_BRAZIL | Batalha_e_Mantovani_2000_Rev_Brasil_Biol_601_129_145 |
| Malpighiaceae | Banisteriopsis_laevifolia | Flushing | | | | | | | | | | | | | -21.61083333 | -47.61055556 | SP | Santa Rita do Passa Quatro | Cerrado | 1997 | 2 | Cerrado_Pé-de-Gigante_Reserve_SP_BRAZIL | Batalha_e_Mantovani_2000_Rev_Brasil_Biol_601_129_145 |
| Malpighiaceae | Banisteriopsis_muricata | Flowering | | | | | | | | | 0 | 0 | 0 | 0 | -21.61083333 | -47.61055556 | SP | Santa Rita do Passa Quatro | Cerrado | 1995 | 2 | Cerrado_Pé-de-Gigante_Reserve_SP_BRAZIL | Batalha_e_Mantovani_2000_Rev_Brasil_Biol_601_129_145 |
| Malpighiaceae | Banisteriopsis_muricata | Fruiting | | | | | | | | | 0 | 0 | 0 | 0 | -21.61083333 | -47.61055556 | SP | Santa Rita do Passa Quatro | Cerrado | 1995 | 2 | Cerrado_Pé-de-Gigante_Reserve_SP_BRAZIL | Batalha_e_Mantovani_2000_Rev_Brasil_Biol_601_129_145 |
| Malpighiaceae | Banisteriopsis_muricata | Leaf_fall | | | | | | | | | | | | | -21.61083333 | -47.61055556 | SP | Santa Rita do Passa Quatro | Cerrado | 1995 | 2 | Cerrado_Pé-de-Gigante_Reserve_SP_BRAZIL | Batalha_e_Mantovani_2000_Rev_Brasil_Biol_601_129_145 |
| Malpighiaceae | Banisteriopsis_muricata | Flushing | | | | | | | | | | | | | -21.61083333 | -47.61055556 | SP | Santa Rita do Passa Quatro | Cerrado | 1995 | 2 | Cerrado_Pé-de-Gigante_Reserve_SP_BRAZIL | Batalha_e_Mantovani_2000_Rev_Brasil_Biol_601_129_145 |
| Malpighiaceae | Banisteriopsis_muricata | Flowering | 0 | 1 | 1 | 1 | 0 | 0 | 0 | 0 | 0 | 0 | 0 | 0 | -21.61083333 | -47.61055556 | SP | Santa Rita do Passa Quatro | Cerrado | 1996 | 2 | Cerrado_Pé-de-Gigante_Reserve_SP_BRAZIL | Batalha_e_Mantovani_2000_Rev_Brasil_Biol_601_129_145 |
| Malpighiaceae | Banisteriopsis_muricata | Fruiting | 0 | 0 | 0 | 0 | 1 | 1 | 0 | 0 | 0 | 0 | 0 | 0 | -21.61083333 | -47.61055556 | SP | Santa Rita do Passa Quatro | Cerrado | 1996 | 2 | Cerrado_Pé-de-Gigante_Reserve_SP_BRAZIL | Batalha_e_Mantovani_2000_Rev_Brasil_Biol_601_129_145 |
| Malpighiaceae | Banisteriopsis_muricata | Leaf_fall | | | | | | | | | | | | | -21.61083333 | -47.61055556 | SP | Santa Rita do Passa Quatro | Cerrado | 1996 | 2 | Cerrado_Pé-de-Gigante_Reserve_SP_BRAZIL | Batalha_e_Mantovani_2000_Rev_Brasil_Biol_601_129_145 |
| Malpighiaceae | Banisteriopsis_muricata | Flushing | | | | | | | | | | | | | -21.61083333 | -47.61055556 | SP | Santa Rita do Passa Quatro | Cerrado | 1996 | 2 | Cerrado_Pé-de-Gigante_Reserve_SP_BRAZIL | Batalha_e_Mantovani_2000_Rev_Brasil_Biol_601_129_145 |
| Malpighiaceae | Banisteriopsis_muricata | Flowering | 0 | 1 | | | | | | | | | | | -21.61083333 | -47.61055556 | SP | Santa Rita do Passa Quatro | Cerrado | 1997 | 2 | Cerrado_Pé-de-Gigante_Reserve_SP_BRAZIL | Batalha_e_Mantovani_2000_Rev_Brasil_Biol_601_129_145 |
| Malpighiaceae | Banisteriopsis_muricata | Fruiting | 0 | 0 | | | | | | | | | | | -21.61083333 | -47.61055556 | SP | Santa Rita do Passa Quatro | Cerrado | 1997 | 2 | Cerrado_Pé-de-Gigante_Reserve_SP_BRAZIL | Batalha_e_Mantovani_2000_Rev_Brasil_Biol_601_129_145 |
| Malpighiaceae | Banisteriopsis_muricata | Leaf_fall | | | | | | | | | | | | | -21.61083333 | -47.61055556 | SP | Santa Rita do Passa Quatro | Cerrado | 1997 | 2 | Cerrado_Pé-de-Gigante_Reserve_SP_BRAZIL | Batalha_e_Mantovani_2000_Rev_Brasil_Biol_601_129_145 |
| Malpighiaceae | Banisteriopsis_muricata | Flushing | | | | | | | | | | | | | -21.61083333 | -47.61055556 | SP | Santa Rita do Passa Quatro | Cerrado | 1997 | 2 | Cerrado_Pé-de-Gigante_Reserve_SP_BRAZIL | Batalha_e_Mantovani_2000_Rev_Brasil_Biol_601_129_145 |
| Malpighiaceae | Banisteriopsis_stellaris | Flowering | | | | | | | | | 0 | 0 | 0 | 1 | -21.61083333 | -47.61055556 | SP | Santa Rita do Passa Quatro | Cerrado | 1995 | 2 | Cerrado_Pé-de-Gigante_Reserve_SP_BRAZIL | Batalha_e_Mantovani_2000_Rev_Brasil_Biol_601_129_145 |
| Malpighiaceae | Banisteriopsis_stellaris | Fruiting | | | | | | | | | 0 | 0 | 0 | 0 | -21.61083333 | -47.61055556 | SP | Santa Rita do Passa Quatro | Cerrado | 1995 | 2 | Cerrado_Pé-de-Gigante_Reserve_SP_BRAZIL | Batalha_e_Mantovani_2000_Rev_Brasil_Biol_601_129_145 |
| Malpighiaceae | Banisteriopsis_stellaris | Leaf_fall | | | | | | | | | | | | | -21.61083333 | -47.61055556 | SP | Santa Rita do Passa Quatro | Cerrado | 1995 | 2 | Cerrado_Pé-de-Gigante_Reserve_SP_BRAZIL | Batalha_e_Mantovani_2000_Rev_Brasil_Biol_601_129_145 |
| Malpighiaceae | Banisteriopsis_stellaris | Flushing | | | | | | | | | | | | | -21.61083333 | -47.61055556 | SP | Santa Rita do Passa Quatro | Cerrado | 1995 | 2 | Cerrado_Pé-de-Gigante_Reserve_SP_BRAZIL | Batalha_e_Mantovani_2000_Rev_Brasil_Biol_601_129_145 |
| Malpighiaceae | Banisteriopsis_stellaris | Flowering | 1 | 1 | 1 | 1 | 1 | 1 | 1 | 0 | 0 | 0 | 0 | 1 | -21.61083333 | -47.61055556 | SP | Santa Rita do Passa Quatro | Cerrado | 1996 | 2 | Cerrado_Pé-de-Gigante_Reserve_SP_BRAZIL | Batalha_e_Mantovani_2000_Rev_Brasil_Biol_601_129_145 |
| Malpighiaceae | Banisteriopsis_stellaris | Fruiting | 0 | 1 | 1 | 1 | 1 | 1 | 1 | 0 | 0 | 0 | 0 | 0 | -21.61083333 | -47.61055556 | SP | Santa Rita do Passa Quatro | Cerrado | 1996 | 2 | Cerrado_Pé-de-Gigante_Reserve_SP_BRAZIL | Batalha_e_Mantovani_2000_Rev_Brasil_Biol_601_129_145 |
| Malpighiaceae | Banisteriopsis_stellaris | Leaf_fall | | | | | | | | | | | | | -21.61083333 | -47.61055556 | SP | Santa Rita do Passa Quatro | Cerrado | 1996 | 2 | Cerrado_Pé-de-Gigante_Reserve_SP_BRAZIL | Batalha_e_Mantovani_2000_Rev_Brasil_Biol_601_129_145 |
| Malpighiaceae | Banisteriopsis_stellaris | Flushing | | | | | | | | | | | | | -21.61083333 | -47.61055556 | SP | Santa Rita do Passa Quatro | Cerrado | 1996 | 2 | Cerrado_Pé-de-Gigante_Reserve_SP_BRAZIL | Batalha_e_Mantovani_2000_Rev_Brasil_Biol_601_129_145 |
| Malpighiaceae | Banisteriopsis_stellaris | Flowering | 1 | 1 | | | | | | | | | | | -21.61083333 | -47.61055556 | SP | Santa Rita do Passa Quatro | Cerrado | 1997 | 2 | Cerrado_Pé-de-Gigante_Reserve_SP_BRAZIL | Batalha_e_Mantovani_2000_Rev_Brasil_Biol_601_129_145 |
| Malpighiaceae | Banisteriopsis_stellaris | Fruiting | 0 | 1 | | | | | | | | | | | -21.61083333 | -47.61055556 | SP | Santa Rita do Passa Quatro | Cerrado | 1997 | 2 | Cerrado_Pé-de-Gigante_Reserve_SP_BRAZIL | Batalha_e_Mantovani_2000_Rev_Brasil_Biol_601_129_145 |
| Malpighiaceae | Banisteriopsis_stellaris | Leaf_fall | | | | | | | | | | | | | -21.61083333 | -47.61055556 | SP | Santa Rita do Passa Quatro | Cerrado | 1997 | 2 | Cerrado_Pé-de-Gigante_Reserve_SP_BRAZIL | Batalha_e_Mantovani_2000_Rev_Brasil_Biol_601_129_145 |
| Malpighiaceae | Banisteriopsis_stellaris | Flushing | | | | | | | | | | | | | -21.61083333 | -47.61055556 | SP | Santa Rita do Passa Quatro | Cerrado | 1997 | 2 | Cerrado_Pé-de-Gigante_Reserve_SP_BRAZIL | Batalha_e_Mantovani_2000_Rev_Brasil_Biol_601_129_145 |
| Malpighiaceae | Banisteriopsis_variabilis | Flowering | | | | | | | | | 1 | 0 | 0 | 0 | -21.61083333 | -47.61055556 | SP | Santa Rita do Passa Quatro | Cerrado | 1995 | 2 | Cerrado_Pé-de-Gigante_Reserve_SP_BRAZIL | Batalha_e_Mantovani_2000_Rev_Brasil_Biol_601_129_145 |
| Malpighiaceae | Banisteriopsis_variabilis | Fruiting | | | | | | | | | 1 | 1 | 1 | 0 | -21.61083333 | -47.61055556 | SP | Santa Rita do Passa Quatro | Cerrado | 1995 | 2 | Cerrado_Pé-de-Gigante_Reserve_SP_BRAZIL | Batalha_e_Mantovani_2000_Rev_Brasil_Biol_601_129_145 |
| Malpighiaceae | Banisteriopsis_variabilis | Leaf_fall | | | | | | | | | | | | | -21.61083333 | -47.61055556 | SP | Santa Rita do Passa Quatro | Cerrado | 1995 | 2 | Cerrado_Pé-de-Gigante_Reserve_SP_BRAZIL | Batalha_e_Mantovani_2000_Rev_Brasil_Biol_601_129_145 |
| Malpighiaceae | Banisteriopsis_variabilis | Flushing | | | | | | | | | | | | | -21.61083333 | -47.61055556 | SP | Santa Rita do Passa Quatro | Cerrado | 1995 | 2 | Cerrado_Pé-de-Gigante_Reserve_SP_BRAZIL | Batalha_e_Mantovani_2000_Rev_Brasil_Biol_601_129_145 |
| Malpighiaceae | Banisteriopsis_variabilis | Flowering | 0 | 0 | 0 | 0 | 0 | 0 | 0 | 0 | 1 | 0 | 0 | 0 | -21.61083333 | -47.61055556 | SP | Santa Rita do Passa Quatro | Cerrado | 1996 | 2 | Cerrado_Pé-de-Gigante_Reserve_SP_BRAZIL | Batalha_e_Mantovani_2000_Rev_Brasil_Biol_601_129_145 |
| Malpighiaceae | Banisteriopsis_variabilis | Fruiting | 0 | 0 | 0 | 0 | 0 | 0 | 0 | 0 | 1 | 1 | 1 | 0 | -21.61083333 | -47.61055556 | SP | Santa Rita do Passa Quatro | Cerrado | 1996 | 2 | Cerrado_Pé-de-Gigante_Reserve_SP_BRAZIL | Batalha_e_Mantovani_2000_Rev_Brasil_Biol_601_129_145 |
| Malpighiaceae | Banisteriopsis_variabilis | Leaf_fall | | | | | | | | | | | | | -21.61083333 | -47.61055556 | SP | Santa Rita do Passa Quatro | Cerrado | 1996 | 2 | Cerrado_Pé-de-Gigante_Reserve_SP_BRAZIL | Batalha_e_Mantovani_2000_Rev_Brasil_Biol_601_129_145 |
| Malpighiaceae | Banisteriopsis_variabilis | Flushing | | | | | | | | | | | | | -21.61083333 | -47.61055556 | SP | Santa Rita do Passa Quatro | Cerrado | 1996 | 2 | Cerrado_Pé-de-Gigante_Reserve_SP_BRAZIL | Batalha_e_Mantovani_2000_Rev_Brasil_Biol_601_129_145 |
| Malpighiaceae | Banisteriopsis_variabilis | Flowering | 0 | 0 | | | | | | | | | | | -21.61083333 | -47.61055556 | SP | Santa Rita do Passa Quatro | Cerrado | 1997 | 2 | Cerrado_Pé-de-Gigante_Reserve_SP_BRAZIL | Batalha_e_Mantovani_2000_Rev_Brasil_Biol_601_129_145 |
| Malpighiaceae | Banisteriopsis_variabilis | Fruiting | 0 | 0 | | | | | | | | | | | -21.61083333 | -47.61055556 | SP | Santa Rita do Passa Quatro | Cerrado | 1997 | 2 | Cerrado_Pé-de-Gigante_Reserve_SP_BRAZIL | Batalha_e_Mantovani_2000_Rev_Brasil_Biol_601_129_145 |
| Malpighiaceae | Banisteriopsis_variabilis | Leaf_fall | | | | | | | | | | | | | -21.61083333 | -47.61055556 | SP | Santa Rita do Passa Quatro | Cerrado | 1997 | 2 | Cerrado_Pé-de-Gigante_Reserve_SP_BRAZIL | Batalha_e_Mantovani_2000_Rev_Brasil_Biol_601_129_145 |
| Malpighiaceae | Banisteriopsis_variabilis | Flushing | | | | | | | | | | | | | -21.61083333 | -47.61055556 | SP | Santa Rita do Passa Quatro | Cerrado | 1997 | 2 | Cerrado_Pé-de-Gigante_Reserve_SP_BRAZIL | Batalha_e_Mantovani_2000_Rev_Brasil_Biol_601_129_145 |
| Malpighiaceae | Diplopterys_pubipetala | Flowering | | | | | | | | | 1 | 1 | 1 | 1 | -21.61083333 | -47.61055556 | SP | Santa Rita do Passa Quatro | Cerrado | 1995 | 2 | Cerrado_Pé-de-Gigante_Reserve_SP_BRAZIL | Batalha_e_Mantovani_2000_Rev_Brasil_Biol_601_129_145 |
| Malpighiaceae | Diplopterys_pubipetala | Fruiting | | | | | | | | | 1 | 1 | 1 | 1 | -21.61083333 | -47.61055556 | SP | Santa Rita do Passa Quatro | Cerrado | 1995 | 2 | Cerrado_Pé-de-Gigante_Reserve_SP_BRAZIL | Batalha_e_Mantovani_2000_Rev_Brasil_Biol_601_129_145 |
| Malpighiaceae | Diplopterys_pubipetala | Leaf_fall | | | | | | | | | | | | | -21.61083333 | -47.61055556 | SP | Santa Rita do Passa Quatro | Cerrado | 1995 | 2 | Cerrado_Pé-de-Gigante_Reserve_SP_BRAZIL | Batalha_e_Mantovani_2000_Rev_Brasil_Biol_601_129_145 |
| Malpighiaceae | Diplopterys_pubipetala | Flushing | | | | | | | | | | | | | -21.61083333 | -47.61055556 | SP | Santa Rita do Passa Quatro | Cerrado | 1995 | 2 | Cerrado_Pé-de-Gigante_Reserve_SP_BRAZIL | Batalha_e_Mantovani_2000_Rev_Brasil_Biol_601_129_145 |
| Malpighiaceae | Diplopterys_pubipetala | Flowering | 1 | 0 | 0 | 0 | 0 | 0 | 1 | 1 | 1 | 1 | 1 | 1 | -21.61083333 | -47.61055556 | SP | Santa Rita do Passa Quatro | Cerrado | 1996 | 2 | Cerrado_Pé-de-Gigante_Reserve_SP_BRAZIL | Batalha_e_Mantovani_2000_Rev_Brasil_Biol_601_129_145 |
| Malpighiaceae | Diplopterys_pubipetala | Fruiting | 1 | 0 | 0 | 0 | 0 | 0 | 0 | 0 | 1 | 1 | 1 | 1 | -21.61083333 | -47.61055556 | SP | Santa Rita do Passa Quatro | Cerrado | 1996 | 2 | Cerrado_Pé-de-Gigante_Reserve_SP_BRAZIL | Batalha_e_Mantovani_2000_Rev_Brasil_Biol_601_129_145 |
| Malpighiaceae | Diplopterys_pubipetala | Leaf_fall | | | | | | | | | | | | | -21.61083333 | -47.61055556 | SP | Santa Rita do Passa Quatro | Cerrado | 1996 | 2 | Cerrado_Pé-de-Gigante_Reserve_SP_BRAZIL | Batalha_e_Mantovani_2000_Rev_Brasil_Biol_601_129_145 |
| Malpighiaceae | Diplopterys_pubipetala | Flushing | | | | | | | | | | | | | -21.61083333 | -47.61055556 | SP | Santa Rita do Passa Quatro | Cerrado | 1996 | 2 | Cerrado_Pé-de-Gigante_Reserve_SP_BRAZIL | Batalha_e_Mantovani_2000_Rev_Brasil_Biol_601_129_145 |
| Malpighiaceae | Diplopterys_pubipetala | Flowering | 1 | 0 | | | | | | | | | | | -21.61083333 | -47.61055556 | SP | Santa Rita do Passa Quatro | Cerrado | 1997 | 2 | Cerrado_Pé-de-Gigante_Reserve_SP_BRAZIL | Batalha_e_Mantovani_2000_Rev_Brasil_Biol_601_129_145 |
| Malpighiaceae | Diplopterys_pubipetala | Fruiting | 1 | 0 | | | | | | | | | | | -21.61083333 | -47.61055556 | SP | Santa Rita do Passa Quatro | Cerrado | 1997 | 2 | Cerrado_Pé-de-Gigante_Reserve_SP_BRAZIL | Batalha_e_Mantovani_2000_Rev_Brasil_Biol_601_129_145 |
| Malpighiaceae | Diplopterys_pubipetala | Leaf_fall | | | | | | | | | | | | | -21.61083333 | -47.61055556 | SP | Santa Rita do Passa Quatro | Cerrado | 1997 | 2 | Cerrado_Pé-de-Gigante_Reserve_SP_BRAZIL | Batalha_e_Mantovani_2000_Rev_Brasil_Biol_601_129_145 |
| Malpighiaceae | Diplopterys_pubipetala | Flushing | | | | | | | | | | | | | -21.61083333 | -47.61055556 | SP | Santa Rita do Passa Quatro | Cerrado | 1997 | 2 | Cerrado_Pé-de-Gigante_Reserve_SP_BRAZIL | Batalha_e_Mantovani_2000_Rev_Brasil_Biol_601_129_145 |
| Malpighiaceae | Diplopterys_pubipetala | Flowering | | | 0 | 0 | 0 | 0 | 0 | 0 | 0 | 1 | 0 | 0 | -22.82916667 | -47.10916667 | SP | Campinas | Mata Atlantica | 1988 | 3 | Semideciduous_forest_Southeastern_Brazil | Morellato_e_Leitão_Filho_1996_Biotropica_282_180_19 |
| Malpighiaceae | Diplopterys_pubipetala | Fruiting | | | | | | | | | | | | | -22.82916667 | -47.10916667 | SP | Campinas | Mata Atlantica | 1988 | 3 | Semideciduous_forest_Southeastern_Brazil | Morellato_e_Leitão_Filho_1996_Biotropica_282_180_19 |
| Malpighiaceae | Diplopterys_pubipetala | Leaf_fall | | | | | | | | | | | | | -22.82916667 | -47.10916667 | SP | Campinas | Mata Atlantica | 1988 | 3 | Semideciduous_forest_Southeastern_Brazil | Morellato_e_Leitão_Filho_1996_Biotropica_282_180_19 |
| Malpighiaceae | Diplopterys_pubipetala | Flushing | | | | | | | | | | | | | -22.82916667 | -47.10916667 | SP | Campinas | Mata Atlantica | 1988 | 3 | Semideciduous_forest_Southeastern_Brazil | Morellato_e_Leitão_Filho_1996_Biotropica_282_180_19 |
| Malpighiaceae | Diplopterys_pubipetala | Flowering | 0 | 0 | 0 | 0 | 0 | 0 | 0 | 0 | 0 | 1 | 0 | 0 | -22.82916667 | -47.10916667 | SP | Campinas | Mata Atlantica | 1989 | 3 | Semideciduous_forest_Southeastern_Brazil | Morellato_e_Leitão_Filho_1996_Biotropica_282_180_19 |
| Malpighiaceae | Diplopterys_pubipetala | Fruiting | | | | | | | | | | | | | -22.82916667 | -47.10916667 | SP | Campinas | Mata Atlantica | 1989 | 3 | Semideciduous_forest_Southeastern_Brazil | Morellato_e_Leitão_Filho_1996_Biotropica_282_180_19 |

| | | | | | | | | | | | | | | | | | | | | | | |
|---|---|---|---|---|---|---|---|---|---|---|---|---|---|---|---|---|---|---|---|---|---|---|
| Malpighiaceae | Diplopterys_pubipetala | Leaf_fall | | | | | | | | | | | | | -22.82916667 | -47.10916667 | SP | Campinas | Mata Atlantica | 1989 | 3 | Semideciduous_forest_Southeastern_Brazil | Morellato_e_Leitão_Filho_1996_Biotropica_282_180_19 |
| Malpighiaceae | Diplopterys_pubipetala | Flushing | | | | | | | | | | | | | -22.82916667 | -47.10916667 | SP | Campinas | Mata Atlantica | 1989 | 3 | Semideciduous_forest_Southeastern_Brazil | Morellato_e_Leitão_Filho_1996_Biotropica_282_180_19 |
| Malpighiaceae | Diplopterys_pubipetala | Flowering | 0 | 0 | 0 | 0 | 0 | 0 | 0 | 0 | 0 | 1 | 0 | 0 | -22.82916667 | -47.10916667 | SP | Campinas | Mata Atlantica | 1990 | 3 | Semideciduous_forest_Southeastern_Brazil | Morellato_e_Leitão_Filho_1996_Biotropica_282_180_19 |
| Malpighiaceae | Diplopterys_pubipetala | Fruiting | | | | | | | | | | | | | -22.82916667 | -47.10916667 | SP | Campinas | Mata Atlantica | 1990 | 3 | Semideciduous_forest_Southeastern_Brazil | Morellato_e_Leitão_Filho_1996_Biotropica_282_180_19 |
| Malpighiaceae | Diplopterys_pubipetala | Leaf_fall | | | | | | | | | | | | | -22.82916667 | -47.10916667 | SP | Campinas | Mata Atlantica | 1990 | 3 | Semideciduous_forest_Southeastern_Brazil | Morellato_e_Leitão_Filho_1996_Biotropica_282_180_19 |
| Malpighiaceae | Diplopterys_pubipetala | Flushing | | | | | | | | | | | | | -22.82916667 | -47.10916667 | SP | Campinas | Mata Atlantica | 1990 | 3 | Semideciduous_forest_Southeastern_Brazil | Morellato_e_Leitão_Filho_1996_Biotropica_282_180_19 |
| Malpighiaceae | Diplopterys_pubipetala | Flowering | 0 | 0 | | | | | | | | | | | -22.82916667 | -47.10916667 | SP | Campinas | Mata Atlantica | 1991 | 3 | Semideciduous_forest_Southeastern_Brazil | Morellato_e_Leitão_Filho_1996_Biotropica_282_180_19 |
| Malpighiaceae | Diplopterys_pubipetala | Fruiting | | | | | | | | | | | | | -22.82916667 | -47.10916667 | SP | Campinas | Mata Atlantica | 1991 | 3 | Semideciduous_forest_Southeastern_Brazil | Morellato_e_Leitão_Filho_1996_Biotropica_282_180_19 |
| Malpighiaceae | Diplopterys_pubipetala | Leaf_fall | | | | | | | | | | | | | -22.82916667 | -47.10916667 | SP | Campinas | Mata Atlantica | 1991 | 3 | Semideciduous_forest_Southeastern_Brazil | Morellato_e_Leitão_Filho_1996_Biotropica_282_180_19 |
| Malpighiaceae | Diplopterys_pubipetala | Flushing | | | | | | | | | | | | | -22.82916667 | -47.10916667 | SP | Campinas | Mata Atlantica | 1991 | 3 | Semideciduous_forest_Southeastern_Brazil | Morellato_e_Leitão_Filho_1996_Biotropica_282_180_19 |
| Malpighiaceae | Heteropterys_byrsonimifolia | Flowering | | | | | | | | | 1 | 1 | 1 | 1 | -21.61083333 | -47.61055556 | SP | Santa Rita do Passa Quatro | Cerrado | 1995 | 2 | Cerrado_Pé-de-Gigante_Reserve_SP_BRAZIL | Batalha_e_Mantovani_2000_Rev_Brasil_Biol_601_129_145 |
| Malpighiaceae | Heteropterys_byrsonimifolia | Fruiting | | | | | | | | | 1 | 1 | 1 | 1 | -21.61083333 | -47.61055556 | SP | Santa Rita do Passa Quatro | Cerrado | 1995 | 2 | Cerrado_Pé-de-Gigante_Reserve_SP_BRAZIL | Batalha_e_Mantovani_2000_Rev_Brasil_Biol_601_129_145 |
| Malpighiaceae | Heteropterys_byrsonimifolia | Leaf_fall | | | | | | | | | | | | | -21.61083333 | -47.61055556 | SP | Santa Rita do Passa Quatro | Cerrado | 1995 | 2 | Cerrado_Pé-de-Gigante_Reserve_SP_BRAZIL | Batalha_e_Mantovani_2000_Rev_Brasil_Biol_601_129_145 |
| Malpighiaceae | Heteropterys_byrsonimifolia | Flushing | | | | | | | | | | | | | -21.61083333 | -47.61055556 | SP | Santa Rita do Passa Quatro | Cerrado | 1995 | 2 | Cerrado_Pé-de-Gigante_Reserve_SP_BRAZIL | Batalha_e_Mantovani_2000_Rev_Brasil_Biol_601_129_145 |
| Malpighiaceae | Heteropterys_byrsonimifolia | Flowering | 1 | 1 | 0 | 0 | 0 | 0 | 1 | 1 | 1 | 1 | 1 | 1 | -21.61083333 | -47.61055556 | SP | Santa Rita do Passa Quatro | Cerrado | 1996 | 2 | Cerrado_Pé-de-Gigante_Reserve_SP_BRAZIL | Batalha_e_Mantovani_2000_Rev_Brasil_Biol_601_129_145 |
| Malpighiaceae | Heteropterys_byrsonimifolia | Fruiting | 1 | 1 | 1 | 1 | 0 | 0 | 0 | 0 | 1 | 1 | 1 | 1 | -21.61083333 | -47.61055556 | SP | Santa Rita do Passa Quatro | Cerrado | 1996 | 2 | Cerrado_Pé-de-Gigante_Reserve_SP_BRAZIL | Batalha_e_Mantovani_2000_Rev_Brasil_Biol_601_129_145 |
| Malpighiaceae | Heteropterys_byrsonimifolia | Leaf_fall | | | | | | | | | | | | | -21.61083333 | -47.61055556 | SP | Santa Rita do Passa Quatro | Cerrado | 1996 | 2 | Cerrado_Pé-de-Gigante_Reserve_SP_BRAZIL | Batalha_e_Mantovani_2000_Rev_Brasil_Biol_601_129_145 |
| Malpighiaceae | Heteropterys_byrsonimifolia | Flushing | | | | | | | | | | | | | -21.61083333 | -47.61055556 | SP | Santa Rita do Passa Quatro | Cerrado | 1996 | 2 | Cerrado_Pé-de-Gigante_Reserve_SP_BRAZIL | Batalha_e_Mantovani_2000_Rev_Brasil_Biol_601_129_145 |
| Malpighiaceae | Heteropterys_byrsonimifolia | Flowering | 1 | 1 | | | | | | | | | | | -21.61083333 | -47.61055556 | SP | Santa Rita do Passa Quatro | Cerrado | 1997 | 2 | Cerrado_Pé-de-Gigante_Reserve_SP_BRAZIL | Batalha_e_Mantovani_2000_Rev_Brasil_Biol_601_129_145 |
| Malpighiaceae | Heteropterys_byrsonimifolia | Fruiting | 1 | 1 | | | | | | | | | | | -21.61083333 | -47.61055556 | SP | Santa Rita do Passa Quatro | Cerrado | 1997 | 2 | Cerrado_Pé-de-Gigante_Reserve_SP_BRAZIL | Batalha_e_Mantovani_2000_Rev_Brasil_Biol_601_129_145 |
| Malpighiaceae | Heteropterys_byrsonimifolia | Leaf_fall | | | | | | | | | | | | | -21.61083333 | -47.61055556 | SP | Santa Rita do Passa Quatro | Cerrado | 1997 | 2 | Cerrado_Pé-de-Gigante_Reserve_SP_BRAZIL | Batalha_e_Mantovani_2000_Rev_Brasil_Biol_601_129_145 |
| Malpighiaceae | Heteropterys_byrsonimifolia | Flushing | | | | | | | | | | | | | -21.61083333 | -47.61055556 | SP | Santa Rita do Passa Quatro | Cerrado | 1997 | 2 | Cerrado_Pé-de-Gigante_Reserve_SP_BRAZIL | Batalha_e_Mantovani_2000_Rev_Brasil_Biol_601_129_145 |
| Malpighiaceae | Heteropterys_umbellata | Flowering | | | | | | | | | 0 | 1 | 1 | 1 | -21.61083333 | -47.61055556 | SP | Santa Rita do Passa Quatro | Cerrado | 1995 | 2 | Cerrado_Pé-de-Gigante_Reserve_SP_BRAZIL | Batalha_e_Mantovani_2000_Rev_Brasil_Biol_601_129_145 |
| Malpighiaceae | Heteropterys_umbellata | Fruiting | | | | | | | | | 0 | 0 | 1 | 1 | -21.61083333 | -47.61055556 | SP | Santa Rita do Passa Quatro | Cerrado | 1995 | 2 | Cerrado_Pé-de-Gigante_Reserve_SP_BRAZIL | Batalha_e_Mantovani_2000_Rev_Brasil_Biol_601_129_145 |
| Malpighiaceae | Heteropterys_umbellata | Leaf_fall | | | | | | | | | | | | | -21.61083333 | -47.61055556 | SP | Santa Rita do Passa Quatro | Cerrado | 1995 | 2 | Cerrado_Pé-de-Gigante_Reserve_SP_BRAZIL | Batalha_e_Mantovani_2000_Rev_Brasil_Biol_601_129_145 |
| Malpighiaceae | Heteropterys_umbellata | Flushing | | | | | | | | | | | | | -21.61083333 | -47.61055556 | SP | Santa Rita do Passa Quatro | Cerrado | 1995 | 2 | Cerrado_Pé-de-Gigante_Reserve_SP_BRAZIL | Batalha_e_Mantovani_2000_Rev_Brasil_Biol_601_129_145 |
| Malpighiaceae | Heteropterys_umbellata | Flowering | 1 | 1 | 0 | 0 | 0 | 0 | 0 | 0 | 0 | 1 | 1 | 1 | -21.61083333 | -47.61055556 | SP | Santa Rita do Passa Quatro | Cerrado | 1996 | 2 | Cerrado_Pé-de-Gigante_Reserve_SP_BRAZIL | Batalha_e_Mantovani_2000_Rev_Brasil_Biol_601_129_145 |
| Malpighiaceae | Heteropterys_umbellata | Fruiting | 1 | 1 | 0 | 0 | 0 | 0 | 0 | 0 | 0 | 0 | 1 | 1 | -21.61083333 | -47.61055556 | SP | Santa Rita do Passa Quatro | Cerrado | 1996 | 2 | Cerrado_Pé-de-Gigante_Reserve_SP_BRAZIL | Batalha_e_Mantovani_2000_Rev_Brasil_Biol_601_129_145 |
| Malpighiaceae | Heteropterys_umbellata | Leaf_fall | | | | | | | | | | | | | -21.61083333 | -47.61055556 | SP | Santa Rita do Passa Quatro | Cerrado | 1996 | 2 | Cerrado_Pé-de-Gigante_Reserve_SP_BRAZIL | Batalha_e_Mantovani_2000_Rev_Brasil_Biol_601_129_145 |
| Malpighiaceae | Heteropterys_umbellata | Flushing | | | | | | | | | | | | | -21.61083333 | -47.61055556 | SP | Santa Rita do Passa Quatro | Cerrado | 1996 | 2 | Cerrado_Pé-de-Gigante_Reserve_SP_BRAZIL | Batalha_e_Mantovani_2000_Rev_Brasil_Biol_601_129_145 |
| Malpighiaceae | Heteropterys_umbellata | Flowering | 1 | 1 | | | | | | | | | | | -21.61083333 | -47.61055556 | SP | Santa Rita do Passa Quatro | Cerrado | 1997 | 2 | Cerrado_Pé-de-Gigante_Reserve_SP_BRAZIL | Batalha_e_Mantovani_2000_Rev_Brasil_Biol_601_129_145 |
| Malpighiaceae | Heteropterys_umbellata | Fruiting | 1 | 1 | | | | | | | | | | | -21.61083333 | -47.61055556 | SP | Santa Rita do Passa Quatro | Cerrado | 1997 | 2 | Cerrado_Pé-de-Gigante_Reserve_SP_BRAZIL | Batalha_e_Mantovani_2000_Rev_Brasil_Biol_601_129_145 |
| Malpighiaceae | Heteropterys_umbellata | Leaf_fall | | | | | | | | | | | | | -21.61083333 | -47.61055556 | SP | Santa Rita do Passa Quatro | Cerrado | 1997 | 2 | Cerrado_Pé-de-Gigante_Reserve_SP_BRAZIL | Batalha_e_Mantovani_2000_Rev_Brasil_Biol_601_129_145 |
| Malpighiaceae | Heteropterys_umbellata | Flushing | | | | | | | | | | | | | -21.61083333 | -47.61055556 | SP | Santa Rita do Passa Quatro | Cerrado | 1997 | 2 | Cerrado_Pé-de-Gigante_Reserve_SP_BRAZIL | Batalha_e_Mantovani_2000_Rev_Brasil_Biol_601_129_145 |
| Fabaceae | Inga_edulis | Flowering | | | | | | | | | | 1 | 1 | 0 | -9 | -35.86666667 | AL | Ibateguara | Mata Atlantica | 2005 | 2 | Coimbra_Forest | Cruz_Neto_et_al_2011_Biodivers_Conserv_20_751_765 |
| Fabaceae | Inga_edulis | Fruiting | | | | | | | | | | | | | -9 | -35.86666667 | AL | Ibateguara | Mata Atlantica | 2005 | 2 | Coimbra_Forest | Cruz_Neto_et_al_2011_Biodivers_Conserv_20_751_765 |
| Fabaceae | Inga_edulis | Leaf_fall | | | | | | | | | | | | | -9 | -35.86666667 | AL | Ibateguara | Mata Atlantica | 2005 | 2 | Coimbra_Forest | Cruz_Neto_et_al_2011_Biodivers_Conserv_20_751_765 |
| Fabaceae | Inga_edulis | Flushing | | | | | | | | | | | | | -9 | -35.86666667 | AL | Ibateguara | Mata Atlantica | 2005 | 2 | Coimbra_Forest | Cruz_Neto_et_al_2011_Biodivers_Conserv_20_751_765 |
| Fabaceae | Inga_edulis | Flowering | 0 | 0 | 0 | 0 | 0 | 0 | 0 | 1 | 1 | 1 | 1 | 0 | -9 | -35.86666667 | AL | Ibateguara | Mata Atlantica | 2006 | 2 | Coimbra_Forest | Cruz_Neto_et_al_2011_Biodivers_Conserv_20_751_765 |
| Fabaceae | Inga_edulis | Fruiting | | | | | | | | | | | | | -9 | -35.86666667 | AL | Ibateguara | Mata Atlantica | 2006 | 2 | Coimbra_Forest | Cruz_Neto_et_al_2011_Biodivers_Conserv_20_751_765 |
| Fabaceae | Inga_edulis | Leaf_fall | | | | | | | | | | | | | -9 | -35.86666667 | AL | Ibateguara | Mata Atlantica | 2006 | 2 | Coimbra_Forest | Cruz_Neto_et_al_2011_Biodivers_Conserv_20_751_765 |
| Fabaceae | Inga_edulis | Flushing | | | | | | | | | | | | | -9 | -35.86666667 | AL | Ibateguara | Mata Atlantica | 2006 | 2 | Coimbra_Forest | Cruz_Neto_et_al_2011_Biodivers_Conserv_20_751_765 |
| Fabaceae | Inga_edulis | Flowering | 0 | 0 | 0 | 0 | 0 | 1 | 1 | 1 | | | | | -9 | -35.86666667 | AL | Ibateguara | Mata Atlantica | 2007 | 2 | Coimbra_Forest | Cruz_Neto_et_al_2011_Biodivers_Conserv_20_751_765 |
| Fabaceae | Inga_edulis | Fruiting | | | | | | | | | | | | | -9 | -35.86666667 | AL | Ibateguara | Mata Atlantica | 2007 | 2 | Coimbra_Forest | Cruz_Neto_et_al_2011_Biodivers_Conserv_20_751_765 |
| Fabaceae | Inga_edulis | Leaf_fall | | | | | | | | | | | | | -9 | -35.86666667 | AL | Ibateguara | Mata Atlantica | 2007 | 2 | Coimbra_Forest | Cruz_Neto_et_al_2011_Biodivers_Conserv_20_751_765 |
| Fabaceae | Inga_edulis | Flushing | | | | | | | | | | | | | -9 | -35.86666667 | AL | Ibateguara | Mata Atlantica | 2007 | 2 | Coimbra_Forest | Cruz_Neto_et_al_2011_Biodivers_Conserv_20_751_765 |
| Fabaceae | Inga_thibaudiana | Flowering | | | | | | | | | | 1 | 1 | 1 | -9 | -35.86666667 | AL | Ibateguara | Mata Atlantica | 2005 | 2 | Coimbra_Forest | Cruz_Neto_et_al_2011_Biodivers_Conserv_20_751_765 |
| Fabaceae | Inga_thibaudiana | Fruiting | | | | | | | | | | | | | -9 | -35.86666667 | AL | Ibateguara | Mata Atlantica | 2005 | 2 | Coimbra_Forest | Cruz_Neto_et_al_2011_Biodivers_Conserv_20_751_765 |
| Fabaceae | Inga_thibaudiana | Leaf_fall | | | | | | | | | | | | | -9 | -35.86666667 | AL | Ibateguara | Mata Atlantica | 2005 | 2 | Coimbra_Forest | Cruz_Neto_et_al_2011_Biodivers_Conserv_20_751_765 |
| Fabaceae | Inga_thibaudiana | Flushing | | | | | | | | | | | | | -9 | -35.86666667 | AL | Ibateguara | Mata Atlantica | 2005 | 2 | Coimbra_Forest | Cruz_Neto_et_al_2011_Biodivers_Conserv_20_751_765 |
| Fabaceae | Inga_thibaudiana | Flowering | 0 | 0 | 0 | 0 | 1 | 1 | 0 | 1 | 1 | 1 | 1 | 1 | -9 | -35.86666667 | AL | Ibateguara | Mata Atlantica | 2006 | 2 | Coimbra_Forest | Cruz_Neto_et_al_2011_Biodivers_Conserv_20_751_765 |
| Fabaceae | Inga_thibaudiana | Fruiting | | | | | | | | | | | | | -9 | -35.86666667 | AL | Ibateguara | Mata Atlantica | 2006 | 2 | Coimbra_Forest | Cruz_Neto_et_al_2011_Biodivers_Conserv_20_751_765 |
| Fabaceae | Inga_thibaudiana | Leaf_fall | | | | | | | | | | | | | -9 | -35.86666667 | AL | Ibateguara | Mata Atlantica | 2006 | 2 | Coimbra_Forest | Cruz_Neto_et_al_2011_Biodivers_Conserv_20_751_765 |
| Fabaceae | Inga_thibaudiana | Flushing | | | | | | | | | | | | | -9 | -35.86666667 | AL | Ibateguara | Mata Atlantica | 2006 | 2 | Coimbra_Forest | Cruz_Neto_et_al_2011_Biodivers_Conserv_20_751_765 |
| Fabaceae | Inga_thibaudiana | Flowering | 1 | 0 | 1 | 1 | 0 | 1 | 0 | 1 | | | | | -9 | -35.86666667 | AL | Ibateguara | Mata Atlantica | 2007 | 2 | Coimbra_Forest | Cruz_Neto_et_al_2011_Biodivers_Conserv_20_751_765 |
| Fabaceae | Inga_thibaudiana | Fruiting | | | | | | | | | | | | | -9 | -35.86666667 | AL | Ibateguara | Mata Atlantica | 2007 | 2 | Coimbra_Forest | Cruz_Neto_et_al_2011_Biodivers_Conserv_20_751_765 |
| Fabaceae | Inga_thibaudiana | Leaf_fall | | | | | | | | | | | | | -9 | -35.86666667 | AL | Ibateguara | Mata Atlantica | 2007 | 2 | Coimbra_Forest | Cruz_Neto_et_al_2011_Biodivers_Conserv_20_751_765 |
| Fabaceae | Inga_thibaudiana | Flushing | | | | | | | | | | | | | -9 | -35.86666667 | AL | Ibateguara | Mata Atlantica | 2007 | 2 | Coimbra_Forest | Cruz_Neto_et_al_2011_Biodivers_Conserv_20_751_765 |
